# Epigenetic regulation of PAR4-related platelet activation: mechanistic links between environmental exposure and cardiovascular disease

**DOI:** 10.1101/473728

**Authors:** Laura J. Corbin, Amy E. Taylor, Stephen J. White, Christopher M. Williams, Kurt Taylor, Marion T. van den Bosch, Jack E. Teasdale, Matthew Jones, Mark Bond, Matthew T. Harper, Louise Falk, Alix Groom, Georgina G J Hazell, Lavinia Paternoster, Marcus R. Munafò, Børge G. Nordestgaard, Anne Tybjaerg-Hansen, Stig E. Bojesen, Caroline Relton, Josine L. Min, for the GoDMC Consortium, George Davey Smith, Andrew D. Mumford, Alastair W. Poole, Nicholas J. Timpson

## Abstract

Protease-activated receptor 4 (PAR4) is a potent thrombin receptor. Epigenetic control of the *F2RL3* locus (which encodes for PAR4) via DNA methylation is associated with both smoking and cardiovascular disease. We examined the association between DNA hypomethylation at *F2RL3* and risk of cardiovascular disease, focusing on acute myocardial infarction (AMI) (n=853 cases / 2,352 controls). We used *in vitro* cell models to dissect the role of DNA methylation in regulating expression of *F2RL3.* We investigated the interplay between *F2RL3* DNA methylation and platelet function in human (n=41). Lastly, we used Mendelian randomization to unify observational and functional work by assessing evidence for causal relationships using data from UK Biobank (n=407,141) and CARDIoGRAMplusC4D (n=184,305). Observationally, one standard deviation (SD) decrease in DNA methylation at *F2RL3* was associated with a 25% increase in the odds of AMI. *In vitro*, short-term exposure of cells to cigarette smoke reduced *F2RL3* DNA methylation and increased gene expression. Transcriptional assays flagged a role for a CEBP recognition sequence in modulating the enhancer activity of *F2RL3* exon 2. Lower DNA methylation at *F2RL3* was associated with increased platelet reactivity in human. The estimated casual odds ratio of ischaemic heart disease was 1.03 (95% CI: 1.00, 1.07) per 1 SD decrease in *F2RL3* DNA. In conclusion, we show that DNA methylation-dependent platelet activation is part of a complex system of features contributing to cardiovascular health. Tailoring therapeutic intervention to new knowledge of *F2RL3*/PAR4 function should be explored to ameliorate the detrimental effects of this risk factor on cardiovascular health.

**One sentence summary:** DNA methylation-dependent platelet activation is a likely causal contributor to cardiovascular health.

## Introduction

The mechanisms behind the adverse effects of smoking on cardiovascular health remain incompletely understood, but over the last decade increasing evidence suggests that epigenetic modifications might link environmental exposure and pathology. The use of array-based DNA methylation detection technologies has revealed smoking-related differential DNA methylation patterns in DNA extracted from peripheral blood^1–5^. Specifically, DNA methylation at *F2RL3* appears to show a dose-response relationship with smoking and lower DNA methylation at *F2RL3* has been associated with mortality from all causes, cardiovascular disease (CVD) and cancer^6,7^. Little evidence currently exists as to the causal impact of these observational effects nor the potential mechanisms or therapeutic implications of this route to disease.

*F2RL3* codes for protease-activated receptor 4 (PAR4), a G-protein coupled receptor (GPCR) expressed on the surface of a number of cell types including platelets^8^. Together with protease-activated receptor 1 (PAR-1), PAR4 activates platelets in response to thrombin generated at the site of tissue injury. A small number of missense coding variants in *F2RL3* that alter platelet aggregation and function have been described, providing a link between PAR4 and the heritable inter-individual variation in platelet reactivity^9–11^. Little is known about the functional consequences of epigenetic modifications at the *F2RL3* locus and how regulatory shifts in the complex events controlling platelet function may be manifest in realised health outcomes. We aimed to triangulate evidence from multiple sources in order to not only test hypothetical causal relationships between smoking and methylation related regulation, but also to flag possible targets for therapeutic intervention.

## Methods

We used human data, *in vitro* studies and human testing to investigate the functional consequences of differential DNA methylation at *F2RL3*. Our methods are described in detail in the Materials and Methods section in the Supplementary Appendix (under the same subheadings as those used below) and an overview of the different components of the study can be found in Fig. 1A.

**Fig. 1.**
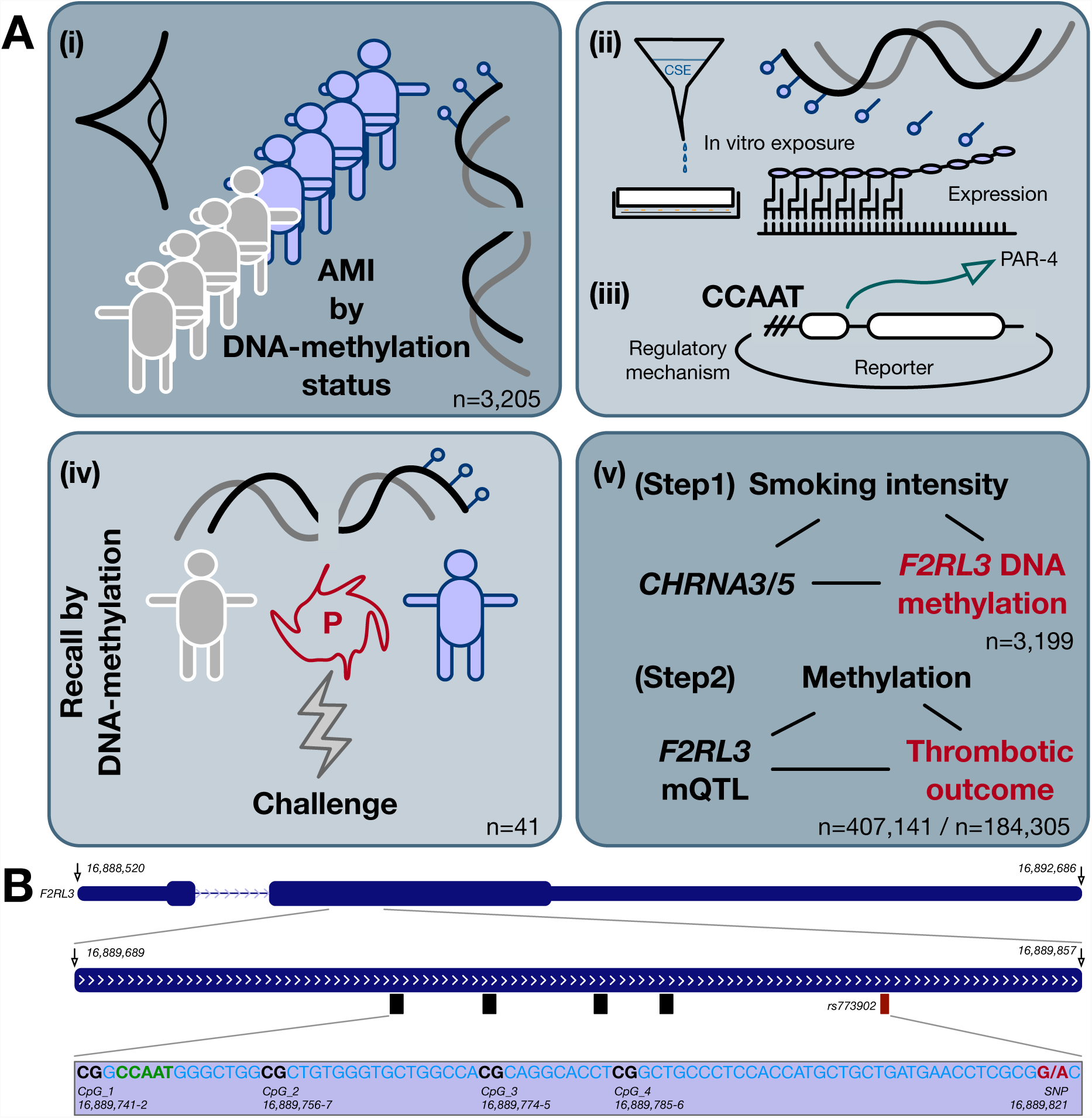
Overview of methods. **(A) Infographic to describe the five components of the study (i)** In a population-based analysis, the observational association between DNA hypomethylation of *F2RL3* and increased risk of cardiovascular disease was quantified, focusing specifically on acute myocardial infarction (AMI). (ii) The effect of exposure to cigarette smoke extract on DNA methylation at *F2RL3* and expression of PAR4 in primary human coronary artery endothelial cells (HCAEC) and acute megakaryocytic leukemia (CMK) cells was tested experimentally. (iii) The impact of a DNA methylation-sensitive CCAAT binding site on *F2RL3* expression was examined to elucidate the mechanism linking DNA methylation to regulation. (iv) The impact of differential DNA methylation at *F2RL3* on platelet function and reactivity in humans was tested in a recall study. (v) A two-step Mendelian randomization strategy was used to assess evidence for causal relationships between smoking, DNA methylation and cardiovascular disease outcomes (data from UK Biobank, n=407,141 and CARDIoGRAMplusC4D, n=184,305). **(B) Schematic to show genomic context of the *F2RL3* DNA methylation sites**. Genomic position and exon structure of *F2RL3* on chromosome 19, with part of exon 2 expanded to show the positions of the four DNA methylation sites CpG_1 to CpG_4 that were assessed by pyrosequencing. Also shown is the single nucleotide polymorphism (SNP) rs773902 (highlighted red) and the CCAAT binding factor recognition sequence (highlighted green). CpG_3 corresponds to the CpG labelled cg03636183 on the Illumina Infinium Human Methylation450 BeadChip (450K) array. Figure produced using UCSC Genome Browser^57^ based on Genome Reference Consortium Human Build 38 patch release 7 (GRCh38.p7).

### (i) *F2RL3* Epidemiology

Using individual participant data in an observational framework, we explored the relationship between smoking, DNA methylation and acute myocardial infarction (AMI). DNA methylation was measured at four CpG sites (CpG_1-4) in exon 2 of *F2RL3* (Fig. 1B) in 853 AMI cases and 2,352 controls from the Copenhagen City Heart Study (Table S1) using pyrosequencing. The CpG_3 at position 16,889,774-5 corresponds to the CpG labelled cg03636183 on the Illumina Infinium BeadChip (27k and 450k). Associations between DNA methylation and self-reported smoking behaviour, incidence of AMI and AMI mortality (within AMI cases) were assessed using linear, logistic and Cox regression, respectively, and adjusted for age, sex and smoking status as appropriate. We also improve the adjustment for smoking exposure in these analyses by using a second smoking related DNA methylation site, the aryl hydrocarbon receptor repressor *(AHRR)*, as a more refined measure of long-term exposure to cigarette smoke.

### (ii) *F2RL3* DNA methylation in a cell model

Two cell types pertinent to CVD were used to evaluate the effect of cigarette smoke on *F2RL3* DNA methylation and mRNA expression. Firstly, in line with previously published work using this model^12,13^, human coronary artery endothelial cells (HCAEC) were exposed to three doses of cigarette smoke extract (CSE) then *F2RL3* DNA methylation and mRNA expression measured^13^. In HCAEC not exposed to CSE, we investigated the impact of global DNA demethylation on *F2RL3* mRNA expression by culture with 5-Azacytidine^14^. In a second set of experiments designed to assess the effect of CSE on a human hematopoietic cell lineage (precursors to platelets) an acute megakaryocytic leukemia cell line (CMK) was used^15^. CMK cells were exposed to four doses of CSE over the course of four days, with *F2RL3* DNA methylation and mRNA expression measured on day five. Expression was also measured in the endogenous control, ribosomal protein lateral stalk subunit P0 (RPLP0).

### (iii) Functional regulation of *F2RL3*

We used a pGL3 reporter vector to test for the presence of an enhancer within a fragment of *F2RL3* exon 2 containing CpG_1 to CpG_4. The potential mechanisms of effects on *F2RL3* expression were explored by transfecting HEK-293 cells with reporter constructs containing different fragments of *F2RL3* to drive expression of luciferase. We began with an expression model in HEK-293 cells with a promoter-less pCpGL reporter vector and in order to set a baseline comparator, the *F2RL3* putative promotor sequence was inserted immediately upstream of the transcription start site adjacent to the luciferase cDNA in pCpGL (pCpGL_*F2RL3*pro). Subsequently, the *F2RL3* exon 2 fragment only was inserted into the pCPGL vector (pCpGL_exon2) and then both the *F2RL3* promoter region and the exon 2 fragment (pCpGL_*F2RL3*pro_exon2). To investigate whether the CCAAT/enhancer binding protein (CEBP) recognition sequence in *F2RL3* exon 2 (see Fig. 1B) was functional and involved in DNA methylation-dependent regulation of *F2RL3,* we assessed luciferase activity again having deleted the CEBP recognition sequence (pCpGL_*F2RL3*pro_exon2 CCAAT deletion). Finally, the HCAEC model was revisited in order to examine the impact of differential DNA methylation on DNA-protein interactions at the locus using chromatin immunoprecipitation (ChIP) in cells cultured with 5-Azacytidine.

### (iv) Differential platelet function in a human experiment

Forty-one never-smoking volunteers (aged 22-24 years) were recruited from the Accessible Resource for Integrated Epigenomic Studies (ARIES) substudy^16^ of the Avon Longitudinal Study of Parents and Children (ALSPAC)^17^ based on having high or low methylation at the *F2RL3* CpG site cg03636183. Recruited individuals provided fresh blood samples that were immediately analysed for platelet reactivity, as assessed by stimulating platelet-rich plasma (PRP) with different concentrations of AYPGKF peptide, a specific agonist of PAR4. Platelet responses were measured using flow cytometry to detect the open conformation of the platelet *α*_IIb_*β*_3_ integrin and platelet surface exposure of P-selectin, both markers of platelet activation during haemostasis. To assess the specificity of any differences observed, the same measurements were made after PRP was treated with SFLLRN, a PAR1 specific agonist. DNA was extracted and targeted pyrosequencing of *F2RL3* carried out to capture the same four positions described previously (Fig. 1B).

Previous studies have shown that the single nucleotide polymorphism (SNP), rs773902, located at 16,889,821 bp on chromosome 19 (Fig. 1B) is associated with platelet function^9,11^. Using existing genetic data in the ALSPAC cohort^17^, we explored the potential impact of this variant both on methylation and platelet reactivity. Methods and results relating to this genetic analysis are described in the Supplementary Appendix (under the subheading ‘Differential platelet function in a human experiment’ in both the Materials and Methods and Additional results sections).

### (v) Mendelian randomization

A two-step epigenetic Mendelian randomization (MR) strategy^18,19^ was used to estimate the causal relationship between smoking, DNA methylation and CVD outcomes (Fig. S1). In the first step, a genetic variant in the *CHRNA5-A3-B4* gene cluster, rs1051730, was used as an instrument for smoking intensity in a one-sample MR design using individual level data from the Copenhagen City Heart Study. Causality was assessed through association of the variant with methylation at CpG_3 in groups stratified by smoking status^20^. In the second step, we carried out a two-sample MR analysis^21,22^ using genetic variants reliably associated with *F2RL3* DNA methylation at CpG_3 in results from the GoDMC Consortium (N=27,750, unpublished) and their effect estimates from association analyses for CVD outcomes in UK Biobank^23,24^ (n=407,141, using individual level data to calculate associations) and CARDIoGRAMplusC4D consortium^25^ (max. n=184,305, using publicly available genome-wide association study summary statistics accessed via MRBase^26^). In UK Biobank, where individual level data was available, three disease outcomes were defined from the most general definition of CVD through ischaemic heart disease (IHD) to the most specific definition of AMI. Approximately equivalent disease definitions were available in CARDIoGRAMplusC4D (myocardial infarction for AMI; coronary heart disease (CHD) for IHD). Sensitivity analyses were performed in UK Biobank to investigate the potential impact of survivorship bias, confounding due to population stratification and smoking on our estimates.

## Results

### (i) *F2RL3* Epidemiology

Observationally, smoking was associated with lower DNA methylation of *F2RL3* across all four CpG sites with the strongest association being seen at CpG_1 (Table S2). The percentage DNA methylation at all four sites was highly correlated in current smokers (pairwise correlations all r>0.77) and former smokers (r between 0.58 and 0.76) and was moderately correlated in those who had never smoked (r between 0.28 and 0.56) (Fig. S2). Amongst smokers, there appeared to be a dose-response relationship such that DNA methylation was lowest in heavier smokers (Table S2).

In a sex-and age-adjusted model, the estimated odds of subsequent AMI observationally was 1.33 (95% CI: 1.21, 1.45) higher per standard deviation (SD) decrease in *F2RL3* DNA methylation (at CpG_1) (Table S3A). This association persisted after adjustment for the potential confounders of active smoking and exposure to passive smoking. Following stratification of samples according to smoking status, a similar association was observed in current or previous smokers (defined as ‘ever smokers’), but not in those who had never smoked, although there was no strong statistical evidence for a difference in estimates between smoking groups (p-value for heterogeneity = 0.36). Adjustment for *AHRR* DNA methylation (a more objective measure of smoking exposure than self-report ^2-4^) did not substantially alter associations between *F2RL3* DNA methylation and AMI, with a fully adjusted model in the full sample yielding an estimated OR of 1.25 (95% CI: 1.11, 1.42) (Table S3A, Fig. S3).

Similar patterns were observed for all-cause mortality in AMI cases assessed over a 23-year period. A hazard ratio of 1.39 (95% CI: 1.22, 1.58) per SD decrease in *F2RL3* DNA methylation in the fully adjusted model suggests that observationally, *F2RL3* DNA methylation is associated with not only risk of AMI, but also with outcome following AMI (Table S4A, Fig. S4).

### (ii) *F2RL3* DNA methylation in a cell model

Exposure of cells to CSE reduced *F2RL3* DNA methylation both in HCAEC (at CpG_1 and CpG_2) and CMK cells (across all CpG sites measured) (Fig. 2A & C). This reduction in DNA methylation was accompanied by a 5.4 (95% CI: 3.9, 7.6, *p*<0.001) fold and 1.7 (95% CI: 1.2, 2.3, *p*<0.01) fold increase in *F2RL3* mRNA levels in HCAEC and CMK cells, respectively (Fig. 2B & D). In CMK cells, expression of the endogenous control, RPLP0, was unchanged. Culture of HCAEC with 5-Azacytidine to induce global DNA hypomethylation resulted in a 14.8 (95% CI: 4.5, 48.3, *p*<0.05) fold increase in *F2RL3* expression over untreated controls while no change in expression was seen in DMSO-treated cells.

**Fig. 2.**
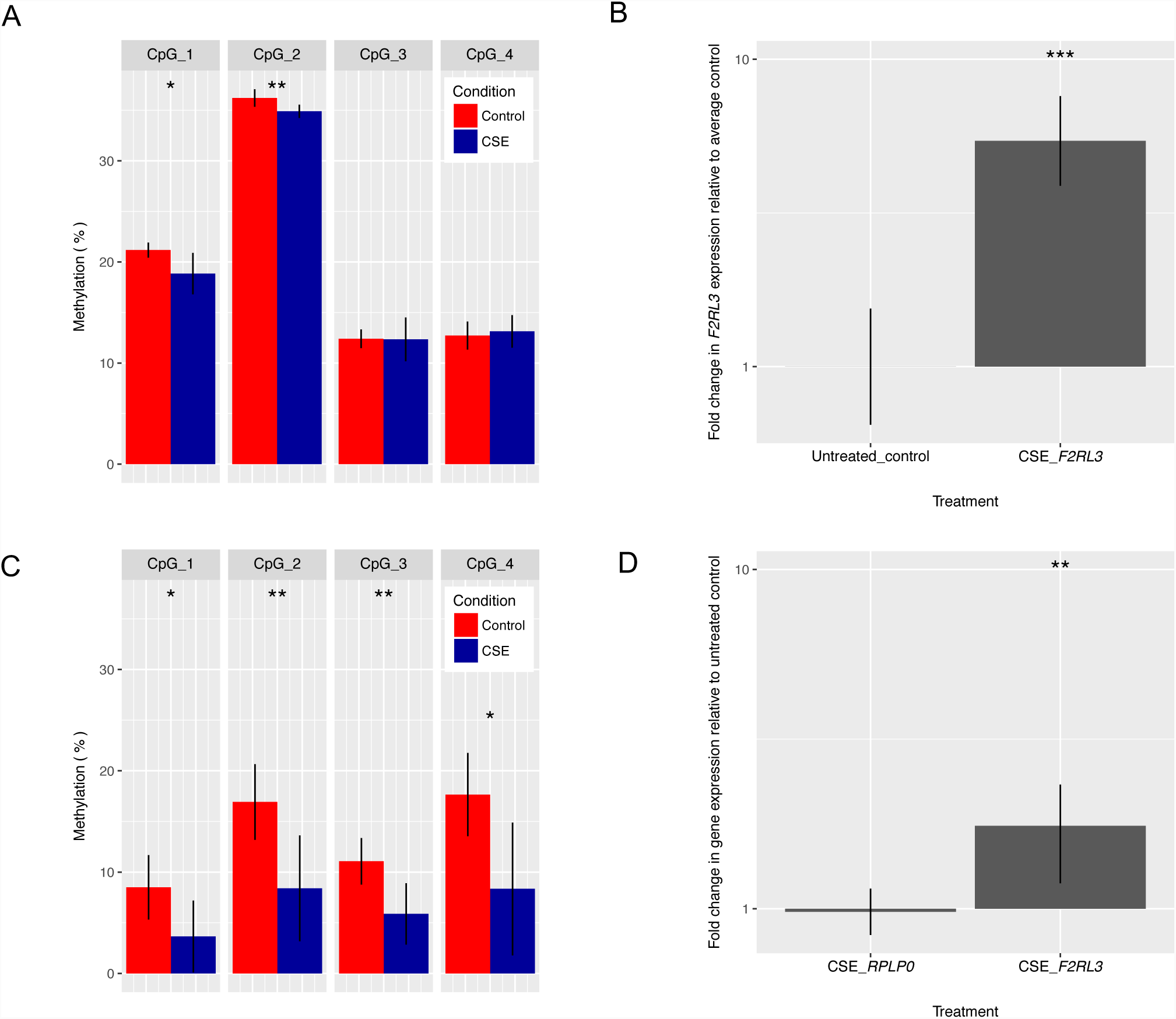
Cigarette smoke exposure and DNA methylation sensitive regulation of *F2RL3* expression in human coronary artery endothelial cells (HCAEC) and acute megakaryocytic leukemia (CMK) cells. **(A)** Exposure of HCAECs to 48 hours of cigarette smoke extract (CSE; blue bars, n=3) reduced DNA methylation at CpG_1 and CpG_2 by approximately 10% compared to untreated controls (red bars, n=3). There were no observed changes in DNA methylation at sites CpG_3 and CpG_4. Data presented are means with 95% confidence intervals and p-values from a two-sample t-test (* = *p*<0.05, ** = *p*<0.01). **(B)** Exposure of HCAECs to 48 hours of CSE led to a 5.4-fold increase in *F2RL3* expression over untreated control cells. Data presented are means with 95% confidence intervals and p-values from a *t*-test (*** = *p*<0.001). **(C)** Exposure of CMKs to 96 hours of CSE (blue bars, n=4) reduced DNA methylation across all CpG sites by approximately 50% compared to untreated controls (red bars, n=4). Data presented are means with 95% confidence intervals and p-values from a two-sample t-test (* = *p*<0.05, ** = *p*<0.01). **(D)** Exposure of CMKs to 96 hours of CSE led to a 1.71-fold increase in *F2RL3* expression over untreated control, whilst expression of the endogenous control *RPLP0* was unchanged (n=5). Data are presented as means with 95% confidence intervals and *p*-values from a *t*-test (** = *p*<0.01).

### (iii) Functional regulation of *F2RL3*

Insertion of a fragment of *F2RL3* exon 2 containing CpG_1 to CpG_4 into a pGL3 reporter vector resulted in a 5.8-fold (95% CI: 3.0, 11.2, *p*=0.007) increase in luciferase activity compared to a pGL3 vector alone, suggesting that this fragment contains a transcriptional enhancer. Combining both the *F2RL3* promoter and the *F2RL3* exon 2 fragment in the pCpGL reporter vector (pCpGL_*F2RL3*pro_exon2) resulted in increased luciferase activity relative to the promotor only construct (Fig. 3) suggesting that the exon 2 region has enhancer activity that acts with the endogenous *F2RL3* promoter. Mutation of the CEBP recognition sequence in exon 2 (pCpGL_*F2RL3*pro_exon2 CCAAT deletion) attenuated luciferase reporter gene activity (Fig. 3) suggesting that whilst this regulatory element was not completely responsible for expression, presence or absence is important and that the CEBP recognition sequence is necessary for full enhancer activity. Culture of HCAEC with the DNA methyltransferase inhibitor, 5-Azacytidine resulted in a 4.9-fold (95% CI: 1.7, 14.0, *p*=0.01) increased occupancy of the *F2RL3* exon 2 CEBP recognition site with CEBP-β (a prototypical isoform abundantly expressed in haemopoietic tissue and endothelium^27^), quantified by ChIP.

**Fig. 3.**
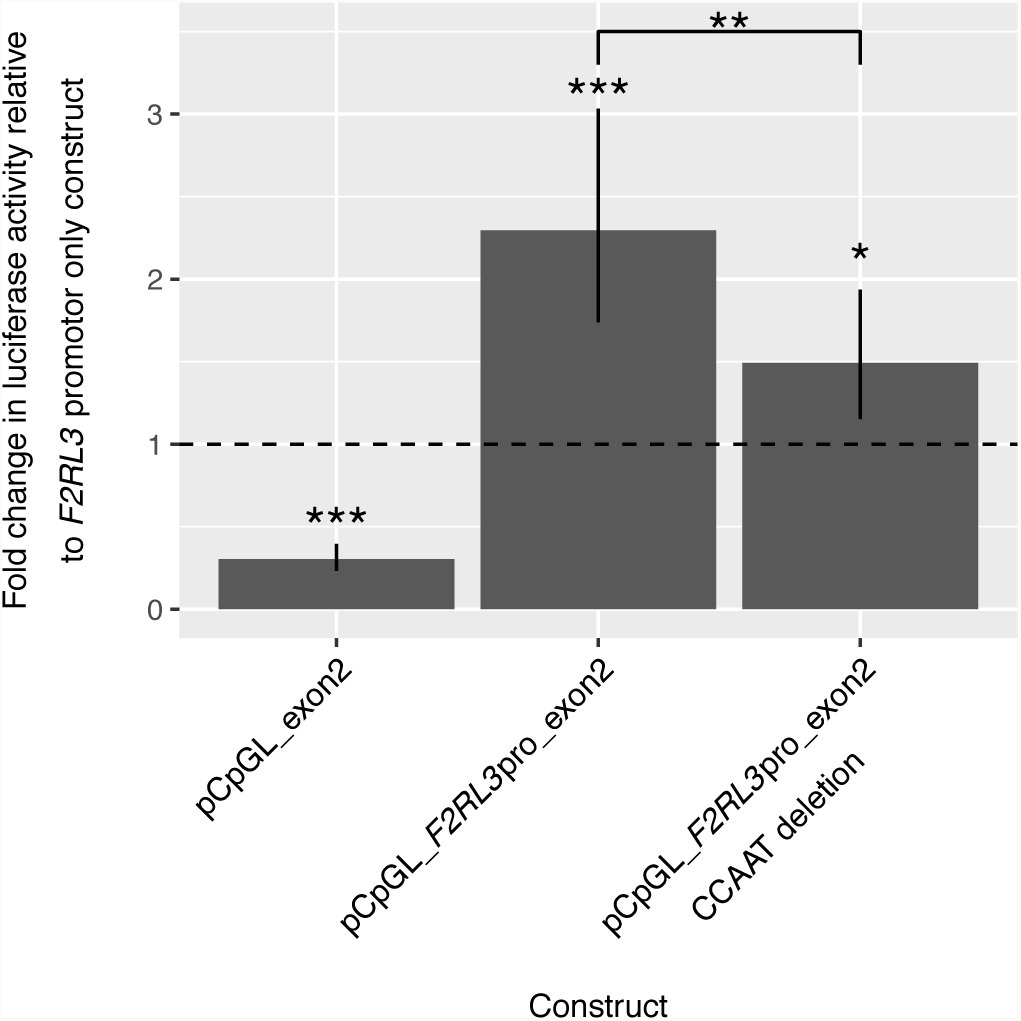
Enhancer activity of the *F2RL3* exon 2 region in HEK-293 cells. Basal activity of luciferase was observed with a pCpGL reporter vector containing the 2kB promoter region of *F2RL3* immediately upstream of the *F2RL3* transcription start site (pCpGL_*F2RL3*pro; represented as the baseline level indicated by the dashed line at 1-fold). Insertion of the *F2RL3* exon 2 fragment alone into the pCpGL vector (pCpGL_exon2) resulted in low level (below baseline), but measurable luciferase activity. Luciferase activity was increased in a pCpGL reporter vector containing both the *F2RL3* promoter region and the exon 2 fragment (pCpGL_*F2RL3*pro_exon2). This enhancer effect of the *F2RL3* exon 2 fragment was abrogated by deletion of the CCAAT recognition site within the exon 2 fragment (pCpGL_*F2RL3*pro_exon2 CCAAT deletion). Data presented are means with 95% confidence intervals and p-values from a *t*-test comparing expression to baseline where * appear directly above the bar and comparing expression with and without the CCAAT deletion where * appears between bars (* = *p*<0.05, ** = *p*<0.01, *** = *p*<0.001).

### (iv) Differential platelet function in a human experiment

ALSPAC participants (see Tables S5 and S6 for descriptive statistics) recalled based on high (retrospective) DNA methylation (n=22) had on average higher contemporary DNA methylation values than the group recalled based on low DNA methylation (n=19) (Table S7; Fig. 4A). The largest difference in DNA methylation (4.8%) was observed at CpG_1 and the correlation across positions ranged from 0.41 to 0.80 (Table S8). The difference in methylation between the groups is small relative to the difference observed between smokers and non-smokers (Fig. S6). The allele frequency at rs773902 in those invited and recruited to this study is shown in Supplementary Tables S5, S6 and S11.

**Fig. 4.**
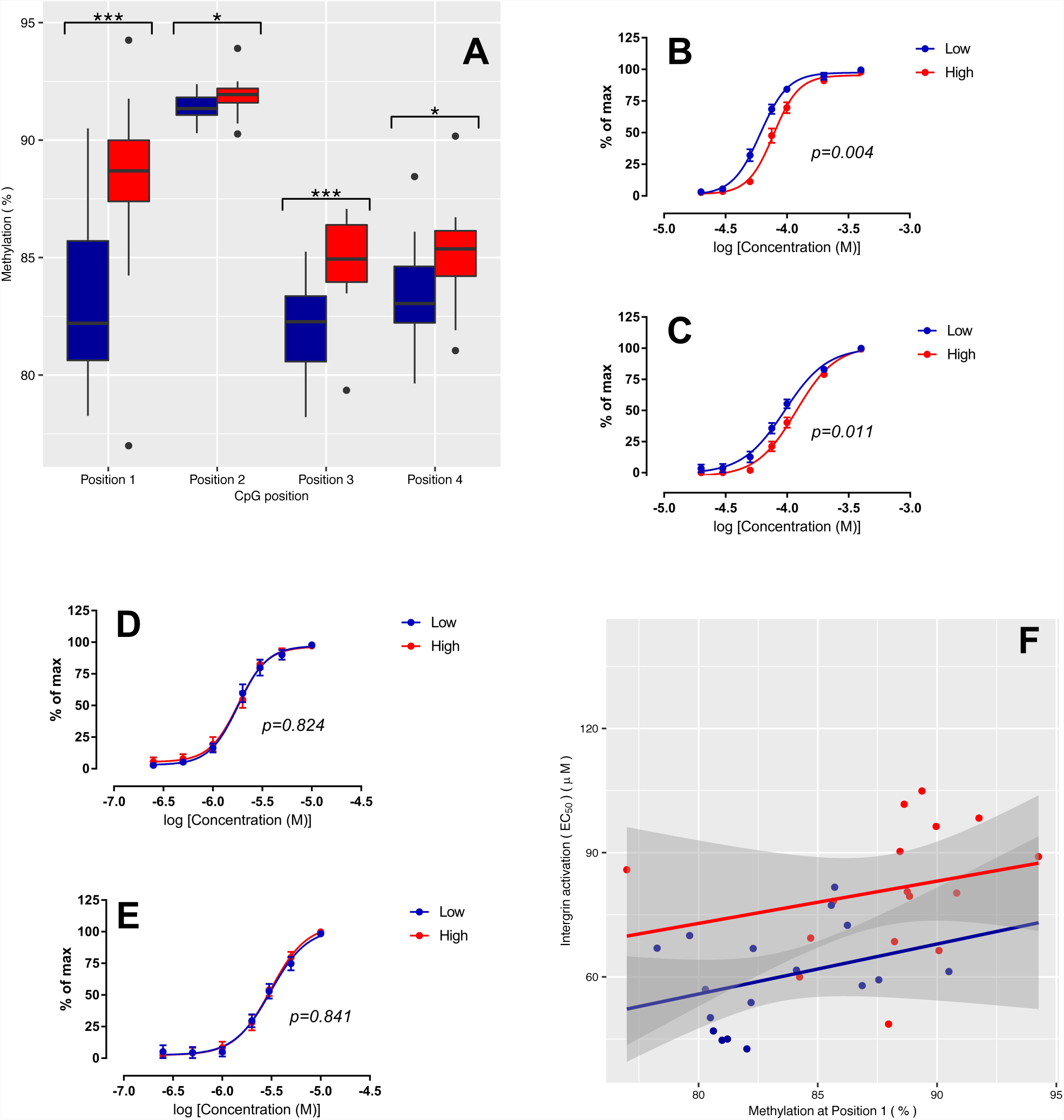
Differential platelet reactivity between groups of ALSPAC participants selected on the basis of low (blue) or high (red) DNA methylation during childhood and adolescence. **(A)** Boxplot showing the between group difference in DNA methylation at the time of platelet reactivity assessment at four CpG positions in *F2RL3* as assessed by two-sample Wilcoxon rank-sum (Mann-Whitney) test (* = *p*<0.05, ** = *p*<0.01, *** = *p*<0.001); **(B & C)** Levels of *α*_IIb_*β*_3_ integrin **(B)** and *α*-granule P-selectin exposure **(C)** following platelet stimulation with the PAR4-specific agonist peptide, AYPGKF. *P*-values derived from a by-group comparison of dose response curves carried out by two-way ANOVA; **(D & E)** Levels of *α*_IIb_*β*_3_ integrin **(D)** and *α*-granule P-selectin exposure **(E)** following platelet stimulation with the PAR-1-specific agonist peptide, SFLLRN. *P*-values derived from a by-group comparison of dose response curves carried out by two-way ANOVA; **(F)** Linear relationship between DNA methylation at CpG_1 and *α*_IIb_*β*_3_ integrin EC_50_ after stimulation (after removal of outliers identified by robust regression of *α*_IIb_*β*_3_ integrin EC_50_ on DNA methylation group (high/low)).

Comparison of dose response curves revealed lower mean half maximum effective concentration (EC_50_) values for both integrin activation (EC_50_, *p*=0.001) and P-selectin exposure (EC_50,_ *p*=0.046) in the low DNA methylation group (Table S9; Fig. 4B & C). These results correspond to an increase in responsiveness with lower DNA methylation. For example, at 75µM AYPGKF, the response in high methylation status individuals was 47.6% compared to 68.6% in the low methylation status individuals, for integrin activation. For P-selectin exposure, the equivalent responses were 21.1% versus 35.6%. These differences could not be explained either by differences in hematological measures (Table S10) or by other measured confounders (Table S11). No between-group differences were observed after the stimulation of platelets via a PAR1-specific agonist (Table S9; Fig. 4D & E). Furthermore, we found no evidence of a between-group difference in the expression of the individual components of integrin (CD41 and CD61) in basal (non-stimulated) samples (Table S9). Accounting for DNA methylation group (high/low), linear regression of integrin activation EC_50_ on DNA methylation at CpG_1 gave evidence of a 1.10 μM (95% CI: −0.26,2.47) decrease in EC_50_ per unit (%) decrease in DNA methylation (Fig. 4F).

### (v) Mendelian randomization

In the Copenhagen City Heart Study, each copy of the minor allele of rs1051730, which is associated with an 0.83 increase in the number of cigarettes smoked per day of (95% CI: 0.24, 1.43) amongst current smokers (Table S12), was associated with decreases in *F2RL3* DNA methylation at CpG_3 of 1.10% (95% CI: −1.93, −0.28) in current smokers and 0.52% (95% CI −1.23, 0.19) in former smokers (Table S13) (we present results from CpG_3, which is the only *F2RL3* site available on the Illumina 450k array (corresponding to cg03636183)). In never smokers, there was no clear evidence that rs1051730 was associated with *F2RL3* DNA methylation (beta per minor allele: −0.08%, 95% CI: −0.78, 0.63). Whilst this apparent lack of association of rs1051730 with methylation in never smokers supports the use of this SNP as an instrument for smoking intensity, power was limited to test for differences between smoking groups (p-value for heterogeneity=0.18). A causal effect estimate was not calculated because the instrument used is not considered to be a good proxy for lifetime exposure to tobacco and therefore, such estimates would likely be biased^20^.

In UK Biobank, causal estimates for the effect of *F2RL3* DNA methylation on CHD/IHD disease risk were generated, with the odds of disease being 1.04 (95% CI: 1.00, 1.08) given a one SD decrease in methylation (Fig. 5). The same estimate generated using data from CARDIoGRAMplusC4D for the equivalent phenotype of coronary heart disease yielded an OR of 1.02 (95% CI: 0.95, 1.09) and the meta-analysed estimate from the two datasets was 1.03 (95% CI: 1.00, 1.07). An analysis in UK Biobank of incident fatal events generated imprecise estimates (Fig. S9). When repeated in a subset of UK Biobank participants designated as ‘white British’, effect estimates are consistent (IHD: OR 1.05 (95% CI: 1.00,1.10)) (Fig. S10). Additional analysis performed in UK Biobank provided no clear evidence of a difference in estimates for ever smokers versus never smokers (Fig. S11). There appeared to be no strong evidence for a causal effect of *F2RL3* methylation on the broadest categorisation of CVD in UK Biobank which includes non-thrombotic events (OR 0.99 (95% CI: 0.97, 1.02) (Fig. 5). Results for analyses using all SNPs separately (including rs773902) and for sensitivity analyses can be found in Supplementary Appendix (Fig. S8 – S13) for both UK Biobank and CARDIoGRAMplusC4D.

**Fig. 5.**
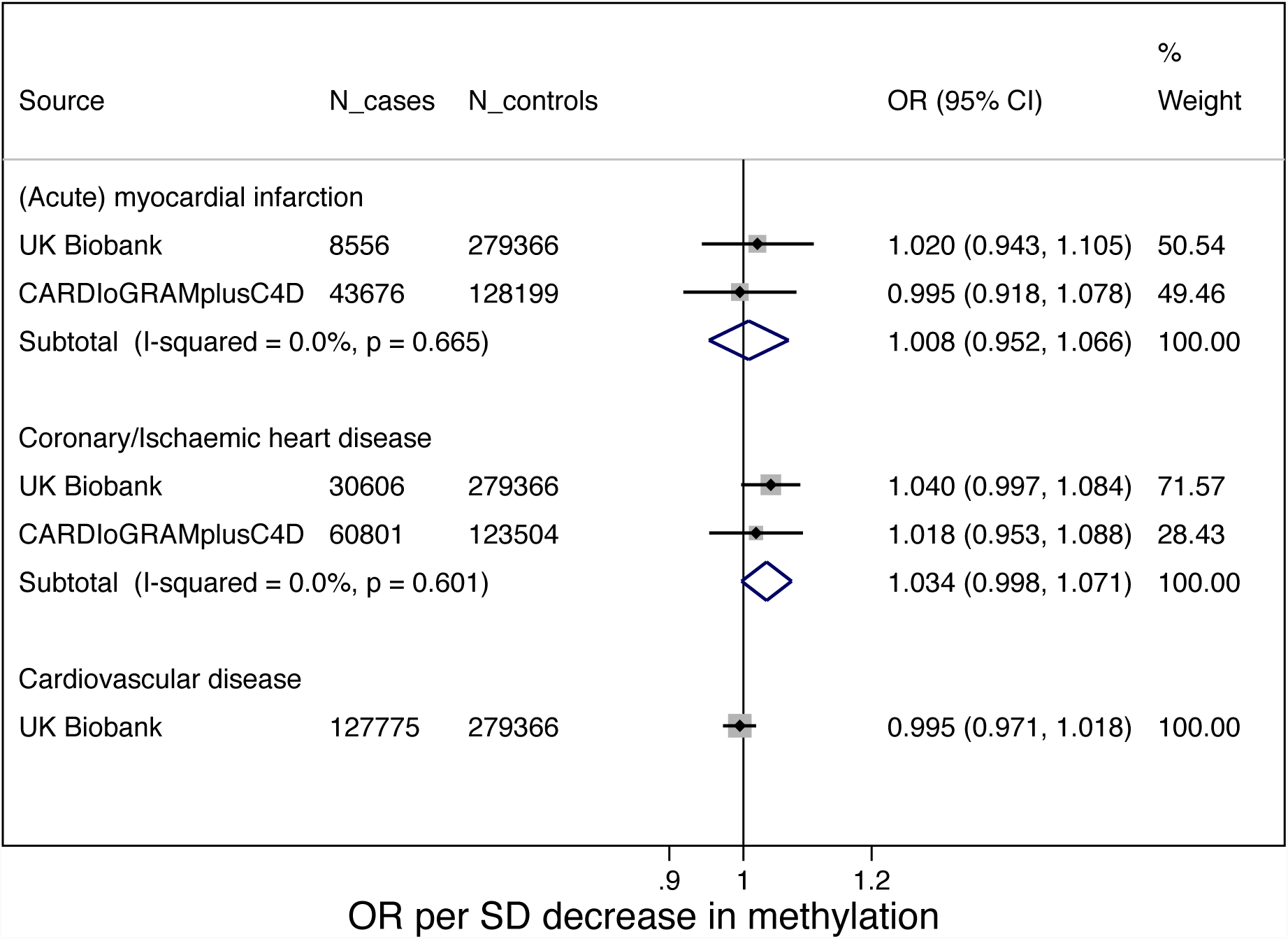
Forest plot of odds ratios for CVD outcomes for each standard deviation decrease in *F2RL3* DNA methylation. Results shown for UK Biobank and CARDIoGRAMplusC4D using a multi-SNP instrument. Where equivalent outcomes were available in both datasets, results were meta-analysed (acute myocardial infarction in UK Biobank matched with myocardial infarction (MI) in CARDIoGRAMplusC4D; ischaemic heart disease matched with coronary heart disease (CHD) in CARDIoGRAMplusC4D). Effect estimates represent the OR (95% CI) for each outcome per one standard deviation unit decrease in DNA methylation at *F2RL3* (CpG_3 / cg03636183). Heterogeneity (Cochran’s Q Statistic) across SNPs: *p*>0.35 for all outcomes in UK Biobank; *p*=0.045 for MI in CARDIoGRAMplusC4D; *p*=0.094 for CHD in CARDIoGRAMplusC4D.

## Discussion

Until now, little direct and causal evidence has been available linking environmental exposure, epigenetic regulation and health outcomes. We have been able to unify evidence from multiple independent sources to address this and to specifically target platelet function as a modifiable aetiological component of smoking-related cardiovascular risk. Observationally, we were able to show that one SD decrease in DNA methylation at *F2RL3* was associated with a 25% increase in the odds of AMI. *In vitro*, short-term exposure of cells to cigarette smoke yielded reductions in *F2RL3* DNA methylation and increased gene expression. With this, transcriptional assays flagged a role for a CEBP recognition sequence in modulating the enhancer activity of *F2RL3* exon 2 and in human, lower DNA methylation at *F2RL3* was associated with increased platelet reactivity.

Cell-based modelling in two CVD-relevant cell types gave evidence of both differential DNA methylation and expression at *F2RL3* because of exposure to aqueous CSE. Functional analysis of the expression of PAR4 identified a CEBP binding site in exon 2 of *F2RL3* as important and occupancy of this site by CEBP-β was shown by ChIP to increase in response to global demethylation. CEBP binding to an identical CCAAT recognition site at a different locus (*MLH1*), is known to be reduced by DNA methylation of a CpG residue in an identical relative position to the CCCAT recognition sequence to that observed with CpG_1^28^. Together, these findings suggest that *F2RL3* expression could be in part constrained by constitutive DNA methylation of CpG sites within an exon 2 enhancer, with methylation reducing CEBP-β occupancy and enhancer activity – smoking disturbs this regulation, reducing methylation and increasing CEBP-β occupancy and *F2RL3* expression. Evidence in support of this hypothesis comes from a recent transcriptome-wide association study that revealed an association between *F2RL3* expression assessed in lymphoblastoid cell lines (LCLs) and smoking (n=92 current versus n=364 never smokers)^29^. However, no relationship between *F2RL3* expression and mortality was observed^29^ and the same association was not seen in a similar study based on whole blood gene expression (n=1,421 current versus n=4,860 never smokers)^30^.

Recruitment and fresh blood sample collection in participants selected based on DNA methylation at *F2RL3* was able not only to recapitulate a DNA methylation gradient independent of smoking, but to also show that this was associated with platelet activation. Although the increase in platelet responsiveness may be directly related to increased PAR4 expression, leading to a leftward shift in a concentration response curve, mechanistically it is possible that this is a result of a change in heterodimer arrangement for PAR4 with other associated GPCRs. It is known, for example, that PAR4 heterodimerises with PAR1 and with P2Y12, and an alteration in the stoichiometry of the association may lead to an altered responsiveness to PAR4 agonism^31,32^. It will therefore be important to determine whether this underlies part of the increased responsiveness when changes in methylation of *F2RL3* gene occur, either naturally or induced by smoking.

Observational evidence was complemented by an MR analysis designed to interrogate the extent to which this relationship is likely to be causal. We were able to estimate the likely causal effect of differential DNA methylation at *F2RL3* on disease, at least in the subcategories of CVD involving coronary thrombosis. It is important to consider that estimates of this nature are subject to the potential complicating factors specific to both the underlying aetiology of these effects and also the nature of the sampling frames used to examine them. For example, we anticipate survivorship bias to be a complicating factor in this analysis. Cases represent a mixture of incident and prevalent disease instances and since the likelihood of death at first presentation is different across the subcategories of disease (AMI, IHD, CVD) the extent of the bias is likely to be different across the different outcomes and may be expected to interact with environmental factors such as smoking. Despite this, causal effect estimates from an analysis in UK Biobank of incident fatalities fell within the 95% confidence interval of estimates from observational analyses and the signals of association in causal analysis were persistent – an important observation when set into the context of the other evidence sources in this investigation.

Differences between the estimated effect of *F2RL3* DNA methylation on risk of AMI from the observational work conducted in CCHS (OR 1.25, 95% CI: 1.11,1.42) and the two sample MR analysis in UK Biobank (1.02, 95% CI: 0.94,1.10) are difficult to assess given precision and the non-specificity of observational estimates, but are not unexpected. In the observational setting, methylation at *F2RL3* is essentially acting as a proxy for exposure to cigarette smoke and therefore, the effect estimate in this case incorporates not only all pathways from smoking to AMI (with PAR4 expression being potentially only one of many), but also all pathways between confounders of smoking and AMI (e.g. alcohol intake, socioeconomic position, educational attainment). In contrast, by using genetic determinants of *F2RL3* DNA methylation we are able to restrict our attention to that pathway alone and therefore the effect estimate from the MR should represent only the effect of methylation at *F2RL3* on the CVD outcome (provided that the standard assumptions of MR are not violated).

Combined evidence here not only implicates *F2RL3* DNA methylation as a likely contributory pathway from smoking to disease risk, but from any feature potentially influencing *F2RL3* regulation in a similar manner. It has been demonstrated that a relatively large proportion of the variation observed in DNA methylation across the genome arises from genetic perturbation^33^, in particular *cis-*acting loci located close to the DNA methylation site they control^34^. Results from the recent GoDMC (unpublished) used here suggest a number of genetic variants in *F2RL3* are associated with methylation at the locus. The difference in minor allele frequency at rs773902 observed in the two arms of our recall experiment suggest part of the non-tobacco smoking related natural variation in DNA methylation at *F2RL3* may be due to genotype at this SNP; similar dual (genetic and environmental) control of DNA methylation has been observed at other loci^35^. When combined with the fact that residue 120 in PAR4 is not within a protein region that is essential for receptor function in the highly homologous PAR1 or in a consensus model structure for all class A GPCRs^36^, it is possible that the associations that have previously been observed between rs773902 and platelet aggregation and reactivity are driven by methylation rather than direct functional effects on the expressed protein.

However, the possibility of this SNP exerting a pleiotropic effect (acting both on DNA methylation *and* PAR4 function directly) cannot be ruled out. There is currently incomplete evidence concerning whether or not the difference in platelet reactivity seen with rs773902 genotype can be attributed to changes in PAR4 expression levels. Despite rs773902 being found to explain 48% of variability in PAR4 reactivity, Edelstein *et al.* (2013)^9^ observed no association between rs773902 and PAR4 protein level and no correlation between expression levels and PAR4 reactivity. Similarly, Morikawa *et al.* (2018)^11^ found surface expression of PAR4 to be comparable across homozygote genotype groups. Results from a search for *F2RL3* expression quantitative trait loci (eQTLs) in data from the Genotype-Tissue Expression Project (GTEx) showed rs773902 as being associated with expression in both adipose-subcutaneous (n=385, *p*=1.8 x 10^−05^) and esophagus-mucosa tissue (n=358, *p*=1.1 x 10^−05^)^37^ whilst other nearby variants acted as eQTLs in other tissues (Table S17A) and in blood^38^ (Table S17B).

We have begun to characterise the pathway from reduced *F2RL3* DNA methylation to increased risk of cardiovascular events but further work is needed to fully understand the mechanisms behind DNA methylation related risk at this locus. It has been proposed that activated platelets have key thromboinflammatory activities with platelets being able to both respond and contribute to inflammatory signals^39,40^. Therefore, the increased activation of platelets that we see in response to decreased DNA methylation may be influenced by or have downstream effects on inflammation^41^. Given that an important part of the development of risk may be associated with increased responsiveness of PAR4 to thrombin, development of PAR4 antagonists as novel antithrombotics is likely to be an attractive intervention^32^. Indeed this is now being realised through the development of several new drug compounds that enable highly specific inhibition of the platelet PAR4 receptor and which potentially have the additional advantage of lower bleeding risk compared with anti-platelet drugs that target other receptors^42–45^. Most recently, French *et al.* (2018)^46^ have added a function-blocking PAR4 antibody to the list of potential PAR4-targeting antithrombotic therapies; using this candidate, they demonstrate inhibition of PAR4 cleavage and activation irrespective of genotype at rs773902. In addition, evidence supporting the combinatorial use of traditional anti-coagulants and new agents along with aspirin, anti-platelet drugs and other treatments suggests that a greater understanding of any individual’s coagulation profile may be advantageous in tailoring intervention^47–52^.

The translational implications of this research are heightened by our growing understanding of the temporal nature of DNA methylation marks. It has been suggested that it can take many years from smoking cessation for DNA methylation levels to return to the level of never smokers, and that the rate of change is CpG site-specific^2,53,54^. In our own collections, it is evident that smoking-induced DNA hypomethylation of the *F2RL3* CpG sites persists for decades after tobacco smoking cessation (Fig. 6). This observation is consistent with the elevated mortality risk that may be seen for ex-cigarette smokers, even after they have given up smoking for many years^55,56^ and has obvious implications for the design of therapy.

**Fig. 6.**
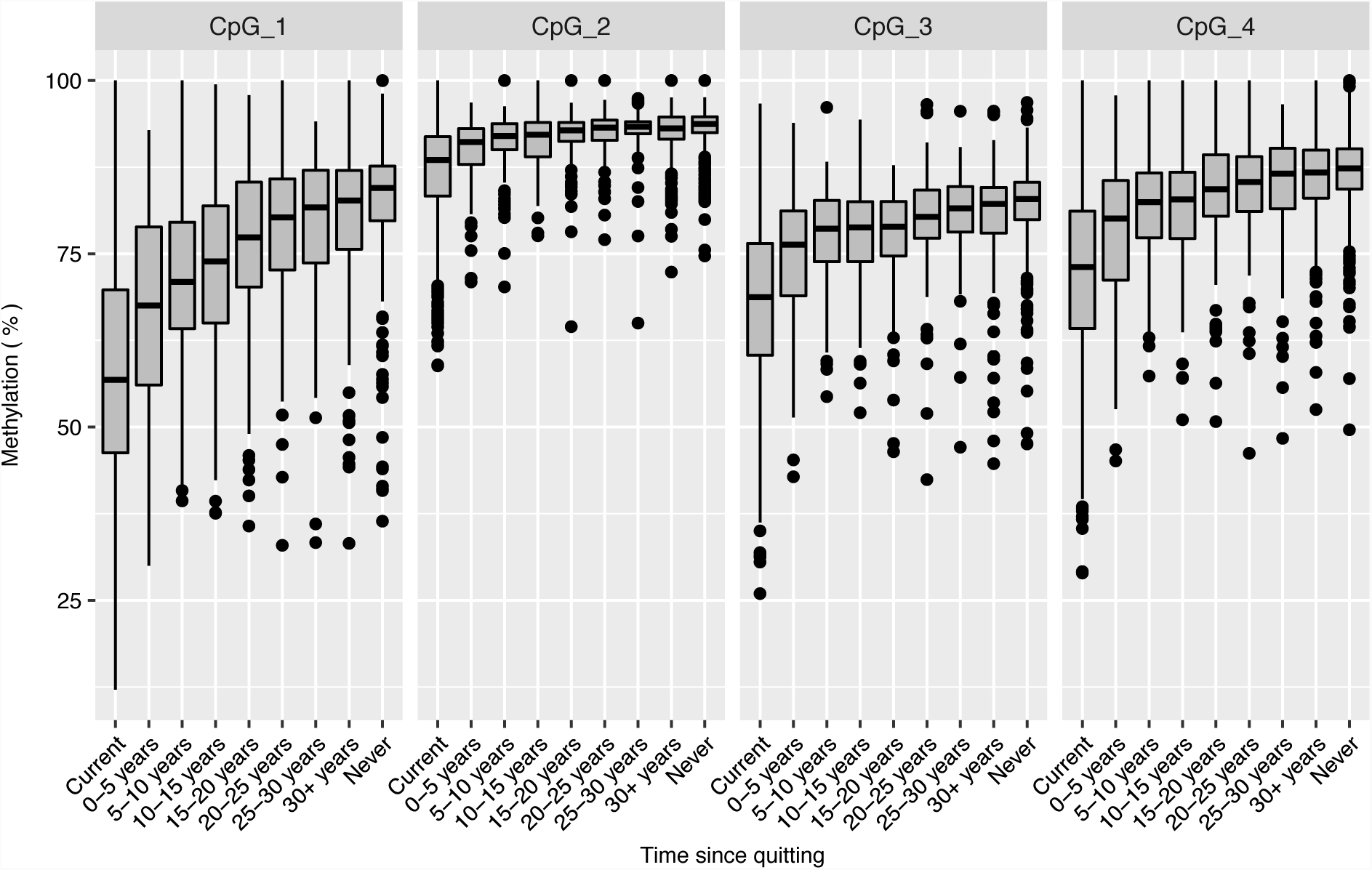
DNA methylation (%) at *F2RL3* by time since quitting amongst participants in the Copenhagen City Heart Study. Box-and-whisker plot of DNA methylation percentage at each CpG of *F2RL3* according to time since quitting tobacco smoking in former smokers. DNA methylation is also presented for current smokers and never smokers. N (with at least one DNA methylation value) for each group is as follows: Current: 1,616, 0-5 years: 170, 5-10 years: 138, 10-15 years: 131: 15-20 years: 110, 20-25 years: 100, 25-30 years: 75, 30+ years: 208, Never: 564.

In summary, the observation that variation in DNA methylation at *F2RL3* appears to have an impact on platelet reactivity suggests this as a pathway through which smoking and potentially other factors affect CVD risk. There are clear therapeutic implications for this work and there may be behavioural and policy implications for this work in a broader context following further investigation.

## Acknowledgments

The authors would like to acknowledge Elizabeth Aitken at the University of Bristol for her contribution to the laboratory work which is presented in the ‘*F2RL3* DNA methylation in a cell model’ section of the methods.

### Funding

This work was specifically supported by the Medical Research Council Integrative Epidemiology Unit (MC_UU_12013/3). NJT is a Wellcome Trust Investigator (202802/Z/16/Z) and works within the University of Bristol NIHR Biomedical Research Centre (BRC). LJC is supported by NJT’s Wellcome Trust Investigator grant (202802/Z/16/Z). NJT, LJC, AT, CR and GDS work in the Medical Research Council Integrative Epidemiology Unit (IEU) at the University of Bristol which is supported by the Medical Research Council (MC_UU_00011/1, MC_UU_00011/5) and the University of Bristol. NJT and CR are supported by the CRUK Integrative Cancer Epidemiology Programme (C18281/A19169). GDS and AWP are funded by the British Heart Foundation (AA/18/1/34219). This study was supported by the NIHR Biomedical Research Centre at the University Hospitals Bristol NHS Foundation Trust and the University of Bristol. The views expressed in this publication are those of the author(s) and not necessarily those of the NHS, the National Institute for Health Research or the Department of Health. The study was supported by the NIHR Bristol Biomedical Research Unit in Cardiovascular Medicine and by British Heart Foundation, grants PG/11/44/28972, FS/12/77/29887 and CH95/001. Funding was also provided by programme and project support from the British Heart Foundation to AWP and KT (RG/15/16/31758, PG/15/96/31854, PG/13/14/30023).

### Avon Longitudinal Study of Parents and Children (ALSPAC)

We are extremely grateful to all the families who took part in this study, the midwives for their help in recruiting them and the whole ALSPAC team, which includes interviewers, computer and laboratory technicians, clerical workers, research scientists, volunteers, managers, receptionists and nurses. The UK Medical Research Council and the Wellcome Trust (Grant ref: 102215/2/13/2) and the University of Bristol provide core support for ALSPAC.

ALSPAC children were genotyped using the Illumina HumanHap550 quad chip genotyping platforms. GWAS data was generated by Sample Logistics and Genotyping Facilities at the Wellcome Sanger Institute and LabCorp (Laboratory Corporation of America) using support from 23andMe. ALSPAC mothers were genotyped using the Illumina human660W-quad array at Centre National de Génotypage (CNG) and genotypes were called with Illumina GenomeStudio.

ARIES was funded by the BBSRC (BBI025751/1 and BB/I025263/1). ARIES is maintained under the auspices of the MRC Integrative Epidemiology Unit at the University of Bristol (MC_UU_12013/2 and MC_UU_12013/8).

### Copenhagen City Heart Study (CCHS)

We acknowledge participants and team of the Copenhagen City Heart Study. The Danish Heart Foundation and the Capital Region of Denmark supported the CCHS.

### UK Biobank

This research has been conducted using the UK Biobank Resource (application 15825).

Data from ALSPAC is made available to researchers through a standard application process (see <u>website</u>). Data from CCHS is made available to researchers upon application to and approval by the Steering Committee.

The Genotype-Tissue Expression (GTEx) Project was supported by the <u>Common Fund</u> of the Office of the Director of the National Institutes of Health, and by NCI, NHGRI, NHLBI, NIDA, NIMH, and NINDS. The data used for the analyses described in this manuscript were obtained from the <u>GTEx Portal</u> on 04/07/18.

## Author contributions

The authors contributed as follows (roles as defined at: https://wellcomeopenresearch.org/for-authors/article-guidelines/research-articles):

Conceptualization: N.J.T., G.D.S, C.R., A.D.M., A.W.P. & S.J.W. Data curation: B.G.N., A.T-H. & S.E.B. Formal analysis: L.J.C., A.E.T., C.M.W., K.T & S.J.W. Funding acquisition: N.J.T., A.W.P & S.J.W. Investigation: L.J.C., A.E.T., S.J.W., C.M.W., K.T., M.T.v.d.B., M.J., M.B., M.T.H., L.F., A.G., G.G.J.H., J.L.M. & J.E.T. Methodology: J.E.T., M.J., M.B., M.T.H & L.P. Supervision: N.J.T., A.D.M., A.W.P & S.J.W. Writing – original draft preparation: L.J.C., A.E.T & S.J.W. Writing – review & editing: All authors.

## Notes

#### Summary of Updates

Minor modification to Supplementary Tables S16A-C. Statistical test for categorical variables revised to be a multinomial logistic regression. This led to very minor changes to p-values in some instances. No change to main manuscript.

