## Supplementary materials for "Epigenetic regulation of PAR4-related platelet activation: mechanistic links between environmental exposure and cardiovascular disease"

### SUPPLEMENTARY FIGURES

**Fig. S1. A two-step epigenetic Mendelian randomization approach applied to smoking and cardiovascular disease (based on Fig. 5 in Relton & Davey Smith (2012)[18]).**

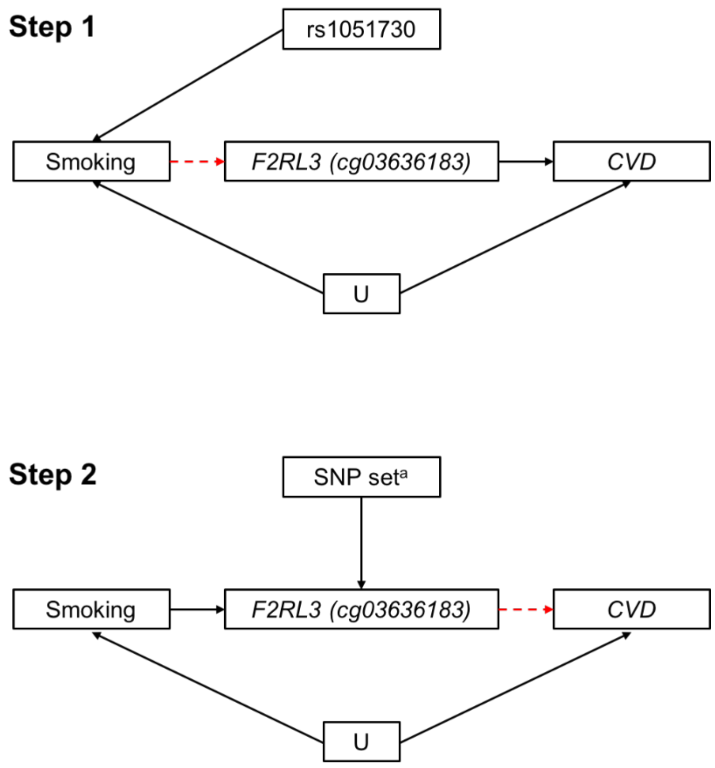

CVD = cardiovascular disease; U = unmeasured confounder; [a] SNP set comprising: rs3848657, rs2227349, rs55961639, rs112791293, rs812847 and rs189158451.

**Fig. S2. The Pearson correlation between the extent of DNA methylation at CpG_1 to CpG_4. Lower triangle shows correlations calculated in 548 never smokers; upper triangle shows correlations calculated in 1,589 current smokers.**

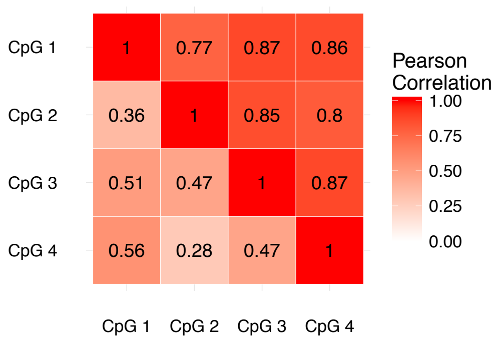

**Fig. S3**. **Comparison of associations of four DNA methylation sites with AMI amongst participants in the Copenhagen City Heart Study.**

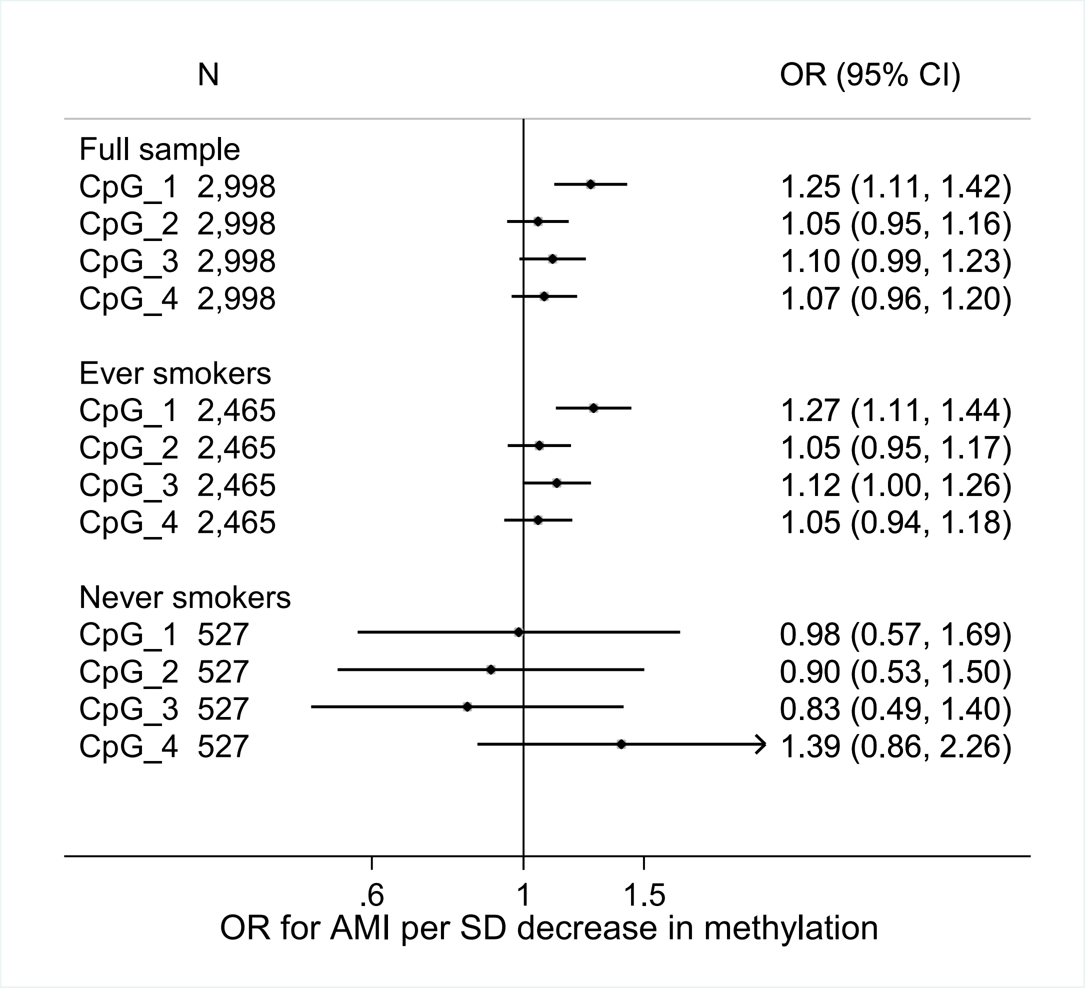

Odds ratios (OR) from logistic regression. Models in full sample are adjusted for: age, sex, smoking status, passive smoking, *AHRR* DNA methylation. Models in ever smokers adjusted for: age, sex, pack years of smoking, passive smoking and *AHRR* DNA methylation. Models in never smokers adjusted for: age, sex, passive smoking and *AHRR* DNA methylation. Individuals with missing information on pack years of smoking were excluded from analyses of ever smokers (N=6). P-values for heterogeneity between CpG sites were as follows: full sample=0.15, ever smokers=0.12, never smokers=0.49.

**Fig. S4. Comparison of associations of four DNA methylation sites with mortality amongst AMI cases in the Copenhagen City Heart Study.**

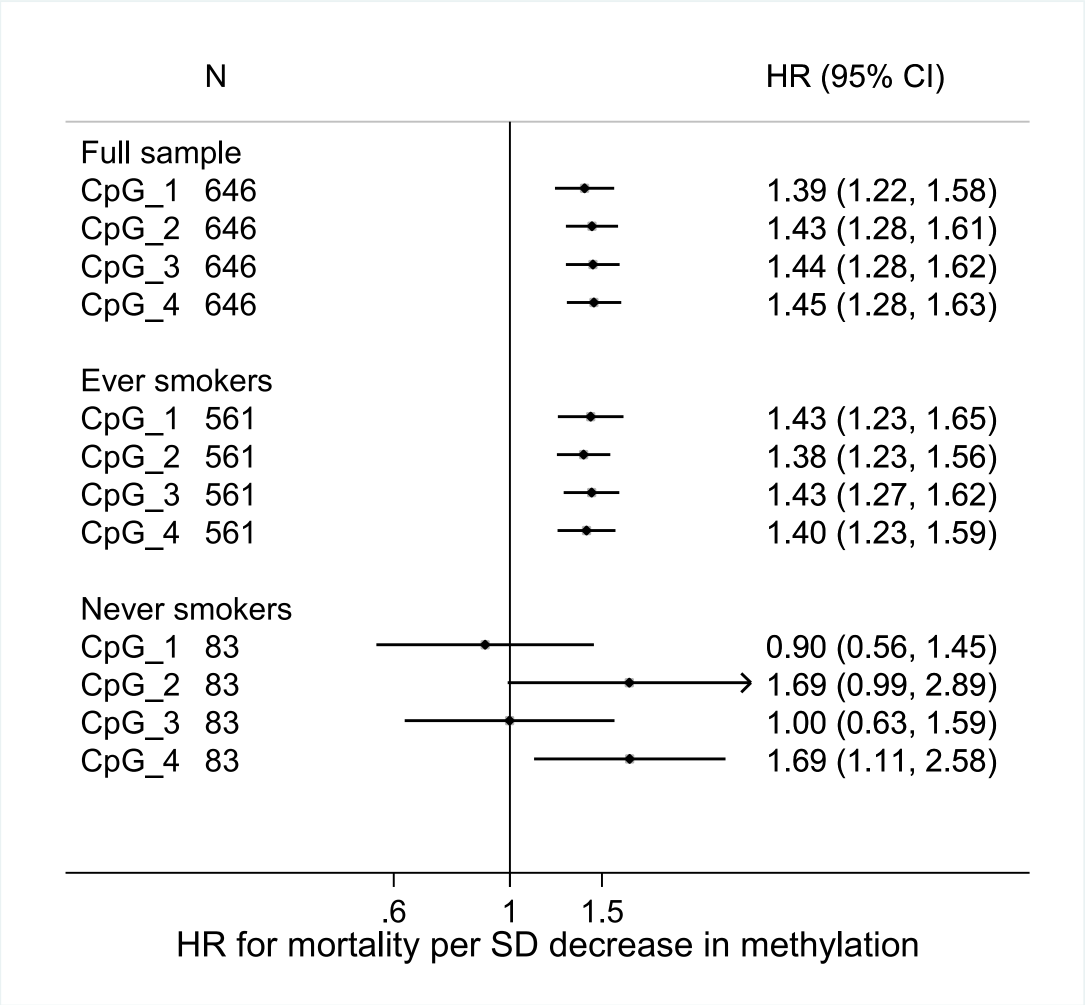

Hazard ratios (HR) from Cox regression. Models in full sample are adjusted for: age, sex, smoking status, passive smoking, *AHRR* DNA methylation. Models in ever smokers adjusted for: age, sex, pack years of smoking, passive smoking and *AHRR* DNA methylation. Models in never smokers adjusted for: age, sex, passive smoking and *AHRR* DNA methylation. Individuals with missing information on pack years of smoking were excluded from analyses of ever smokers (N=2). P-values for heterogeneity between CpG sites were as follows: full sample=0.97, ever smokers=0.98, never smokers=0.11.

**Fig. S5. Results from an assessment of collider bias and its potential impact on the case-only analysis**

**(A) Associations between triglycerides, LDL cholesterol and BMI and AMI in the Copenhagen City Heart Study**

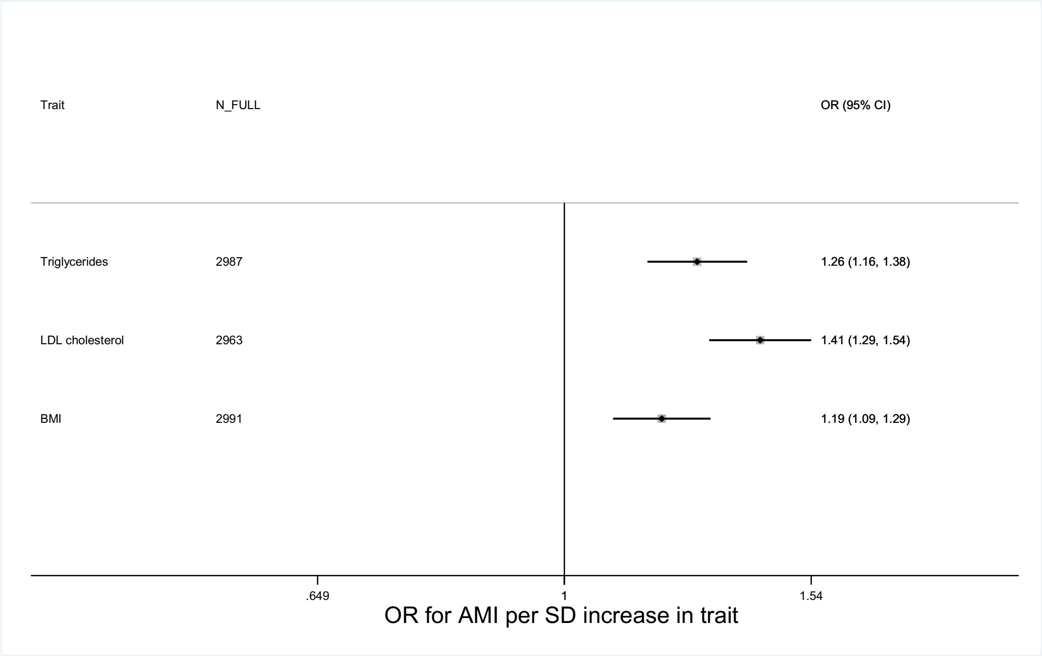

**(B) Associations between between triglyceride, LDL cholesterol and BMI and *F2RL3* DNA methylation (CpG_1) in the Copenhagen City Heart Study.**

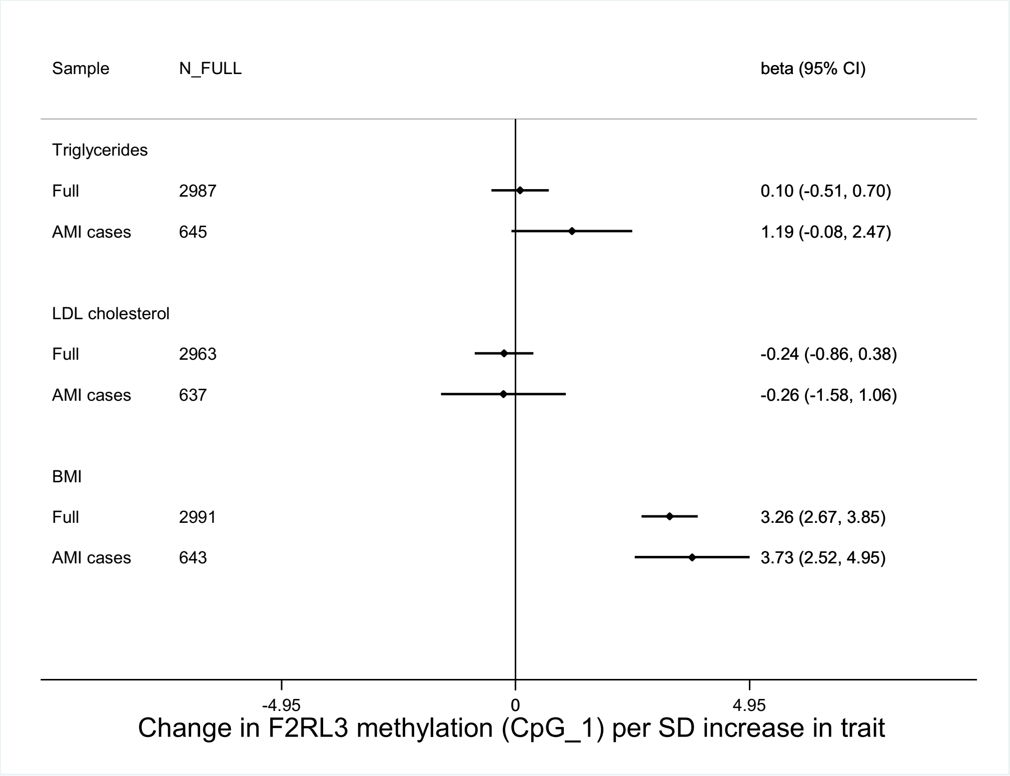

We undertook an analysis of risk factors for AMI incidence to evaluate the possibility of the mortality analysis (where only cases are selected) being affected by collider bias. By comparing the association between AMI incidence risk factors in unselected (Full) and selected (AMI cases) samples, we can identify potential issues. AMI incidence risk factors which are not associated with each other in a random population sample may become associated in a selected sample and act as a source of confounding in the association with an outcome such as mortality (NB. If present such collider bias and confounding would also affect associations between genetic variants and mortality (including in Mendelian randomization analysis), which in a random sample are reasonably assumed to be unaffected by confounding).

Part (a) shows the association of three risk factors with AMI incidence, indicating that there is potential for these factors to be influencing collider bias in the analysis of methylation and mortality in AMI cases. Part (b) shows that these associations between these risk factors and *F2RL3* methylation did not vary between the two groups (Full and AMI cases) and the association between CpG_1 and LDL cholesterol (the strongest AMI incidence risk factor of those tested) remains unassociated in the selected sample, suggesting there is unlikely to be collider bias of any appreciable magnitude affecting our results.

**Fig. S6 Difference in mean % methylation between low (blue) and high (red) methylation groups in the recall study (ALSPAC) relative to the distribution of methylation in smokers, former smokers and never smokers (data from CCHS)**

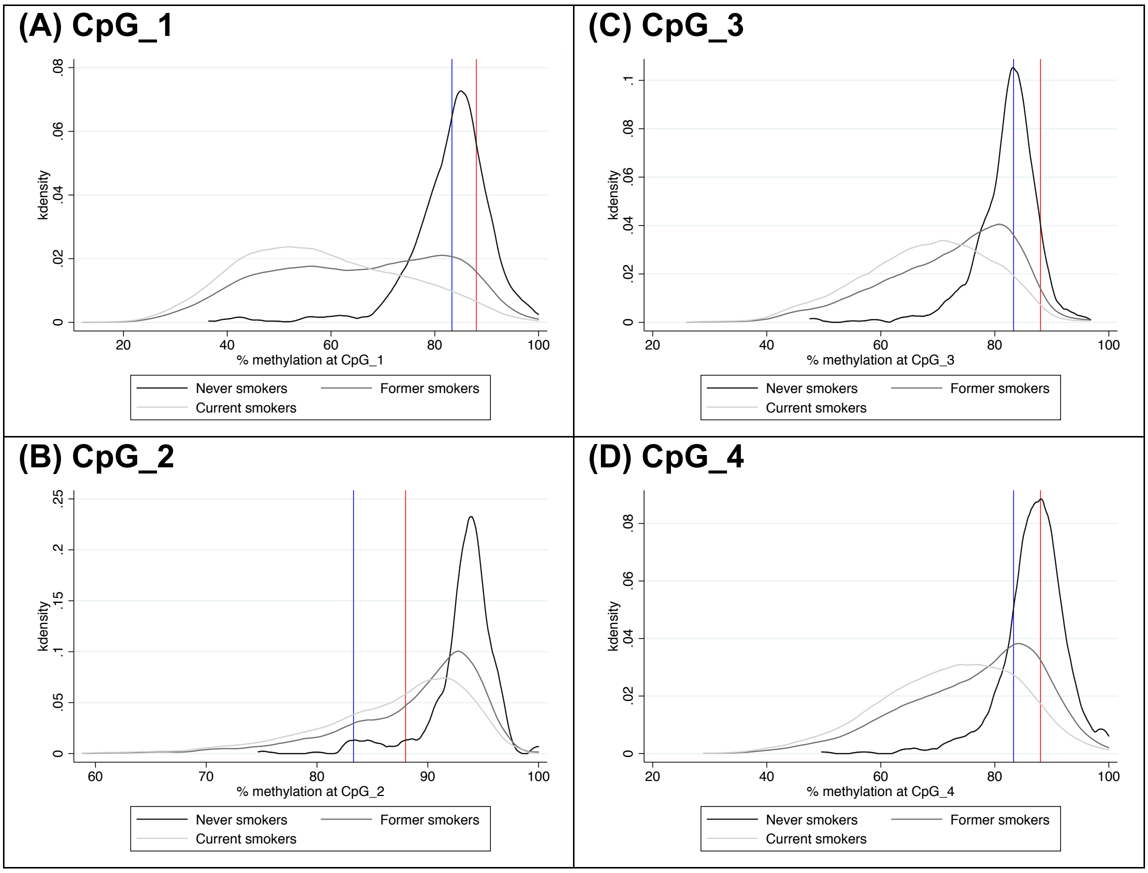

**Fig. S7. Forest plot of odds ratios for CVD outcomes for each additional *F2RL3* DNA methylation decreasing allele.**

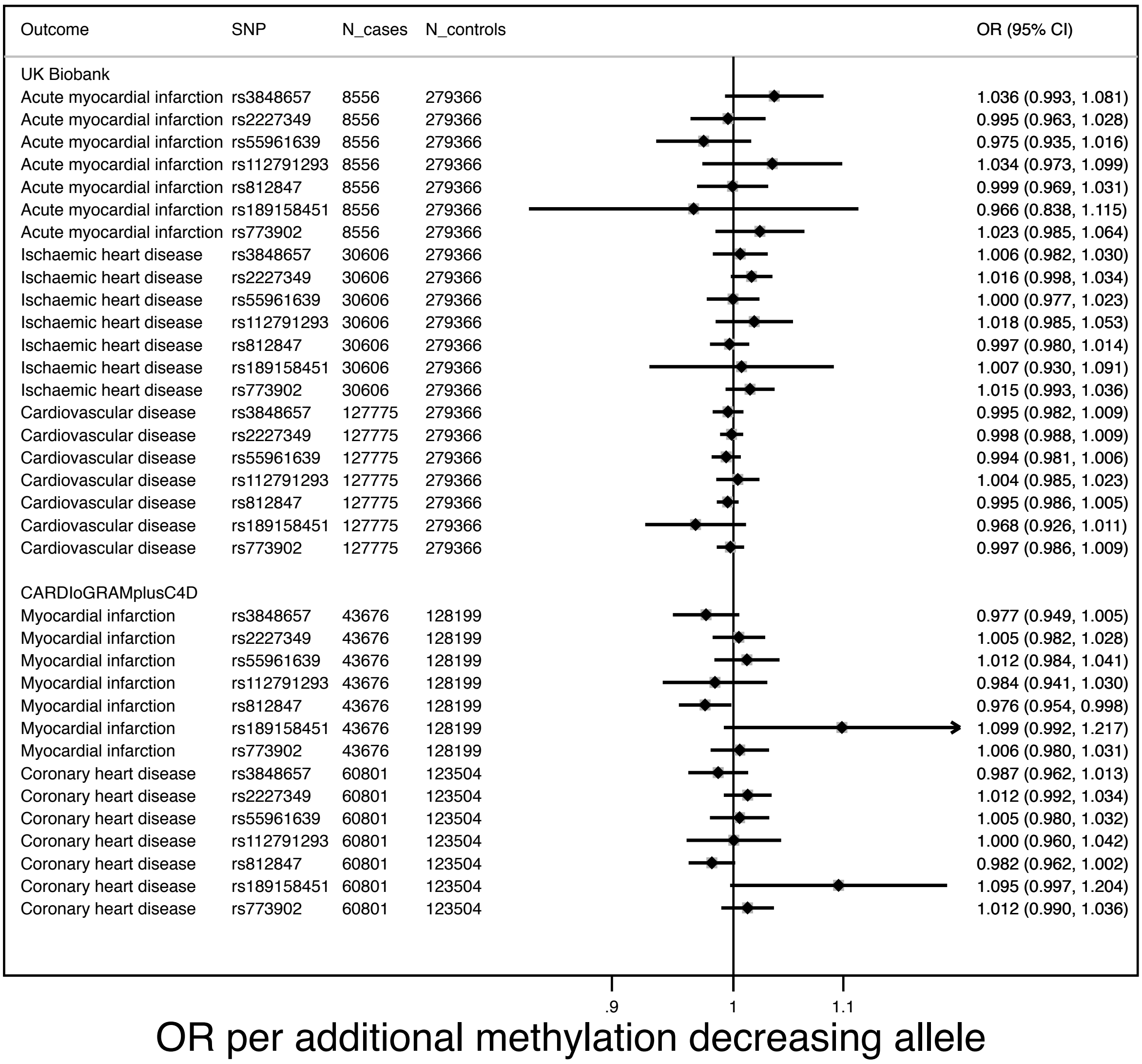

Results in UK Biobank and CARDIoGRAMplusC4D shown for the SNPs included in the multi-SNP instrument and for rs773902. Effect estimates represent the odds ratio (OR) (95% CI) for each outcome per DNA methylation decreasing allele.

**Fig. S8A. MR Result: Forest plot of odds ratios for CVD outcomes for each standard deviation decrease in *F2RL3* DNA methylation by SNP.**

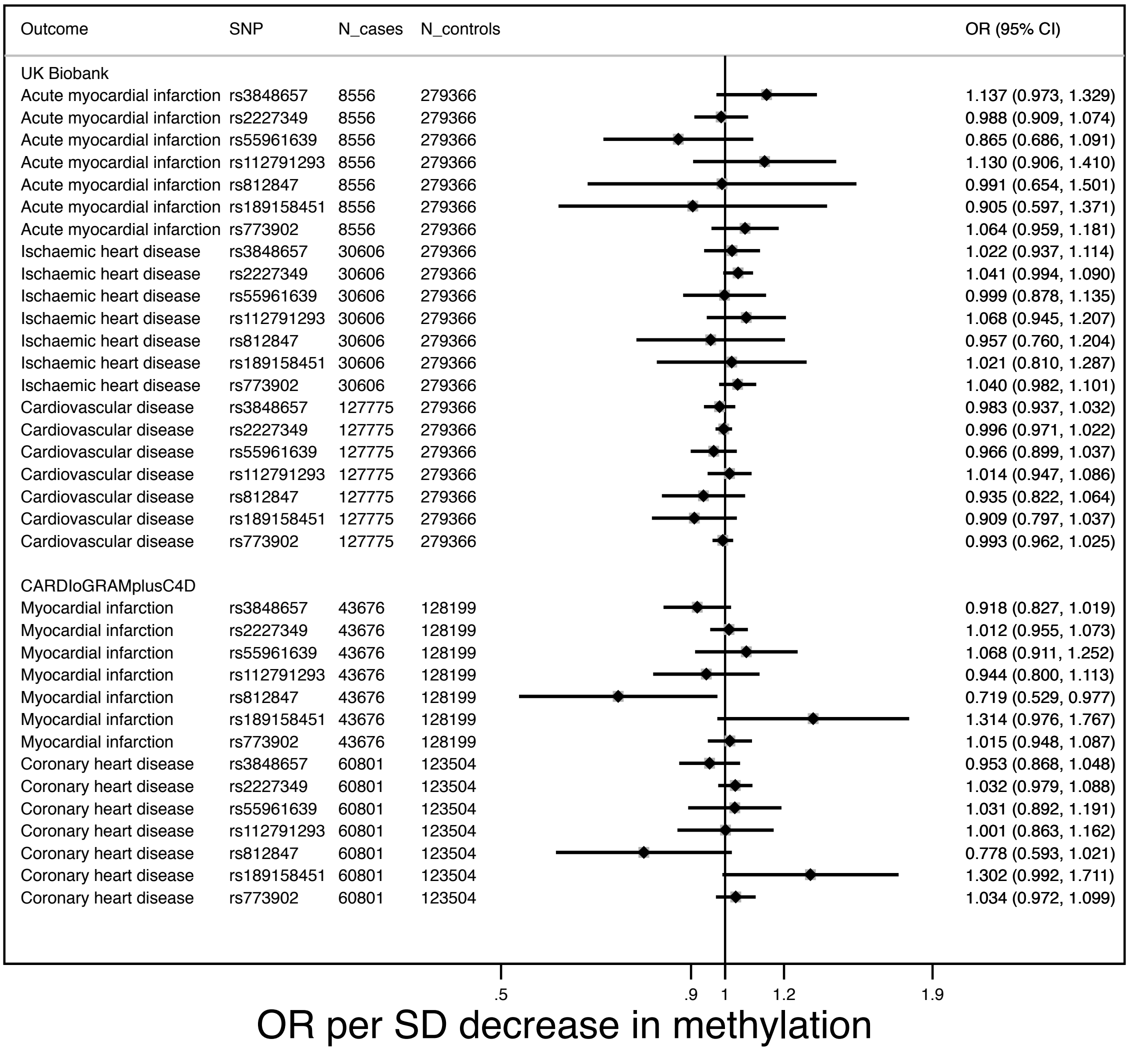

Results in UK Biobank and CARDIoGRAMplusC4D shown for the SNPs included in the multi-SNP instrument and for rs773902. Effect estimates represent the OR (95% CI) for each outcome per one standard deviation unit decrease in DNA methylation at *F2RL3* (CpG_3 / cg03636183). Heterogeneity (Cochran's Q Statistic) across SNPs: $p>0.35$ for all outcomes in UK Biobank; $p=0.045$ for MI in CARDIoGRAMplusC4D; $p=0.094$ for CHD in CARDIoGRAMplusC4D.

**Fig. S8B. MR Result: Forest plot of odds ratios for CVD outcomes for each standard deviation decrease in *F2RL3* DNA methylation in a leave one (SNP) out analysis.**

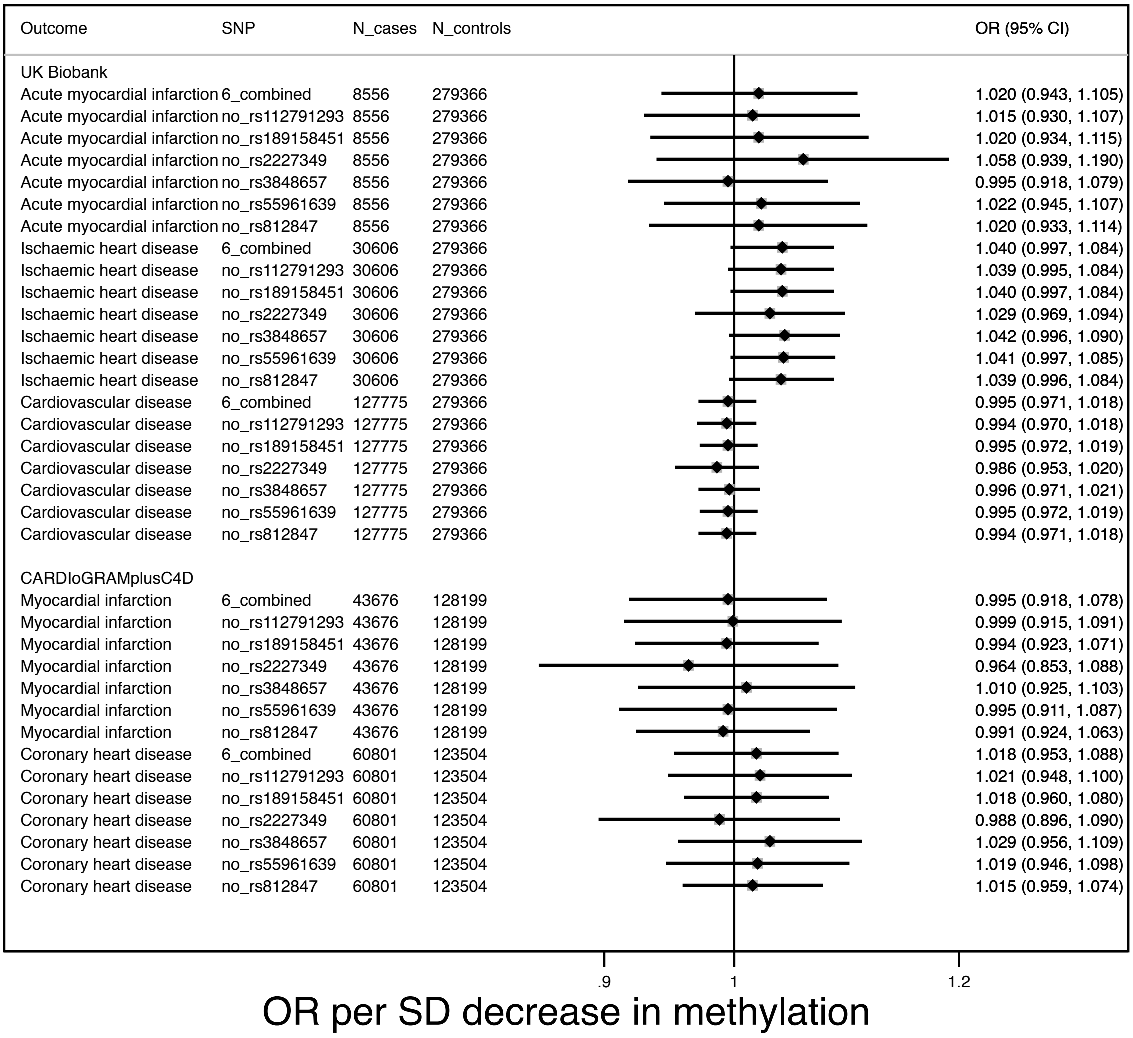

Results in UK Biobank and CARDIoGRAMplusC4D shown for the SNP set (6_combined) and the SNP set without each SNP in turn. Effect estimates represent the OR (95% CI) for each outcome per one standard deviation unit decrease in DNA methylation at *F2RL3* (CpG_3 / cg03636183). Heterogeneity (Cochran's Q Statistic) across SNPs: $p$>0.35 for all outcomes in UK Biobank; $p$=0.045 for MI in CARDIoGRAMplusC4D; $p$=0.094 for CHD in CARDIoGRAMplusC4D.

**Fig. S9. MR Result: Forest plot of odds ratios for CVD outcomes for each standard deviation decrease in *F2RL3* DNA methylation - UK Biobank mortality analysis**

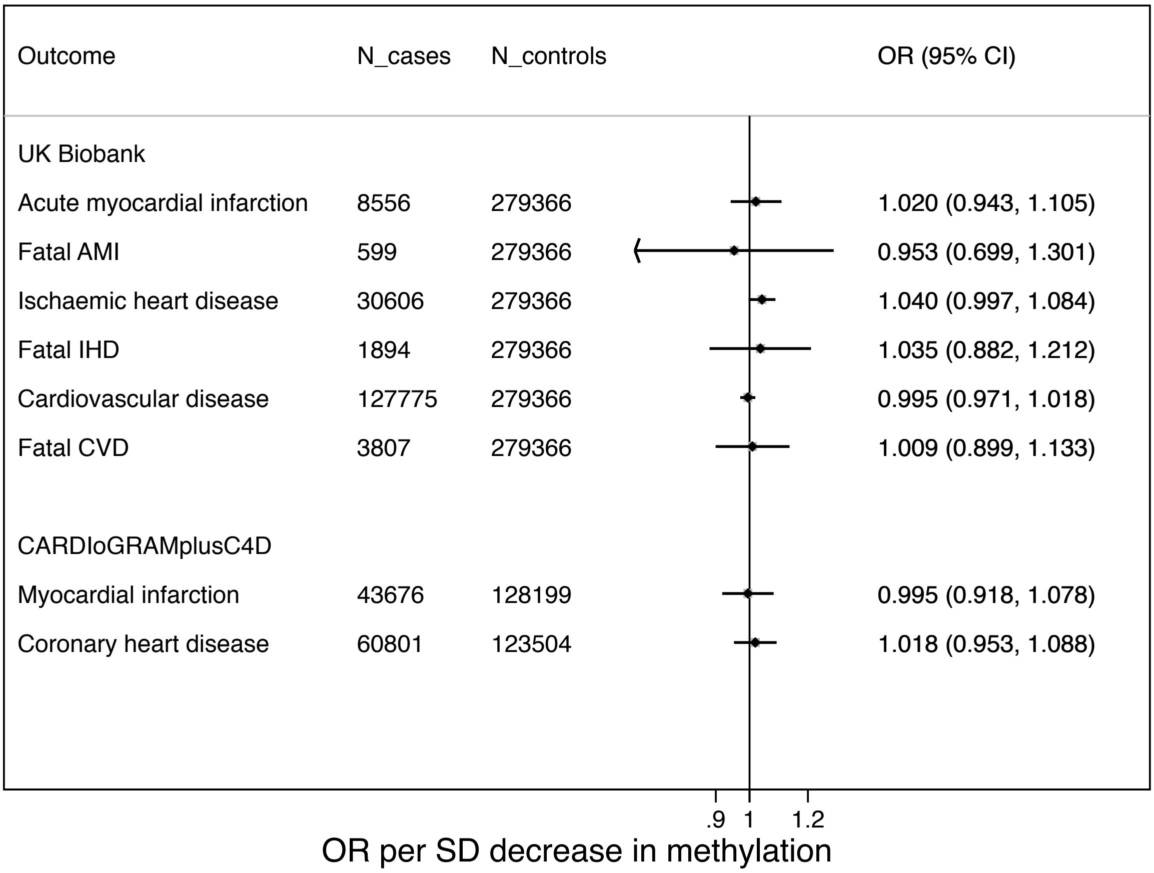

Results shown for the multi-SNP instrument. Effect estimates represent the OR (95% CI) for each outcome per one standard deviation unit decrease in DNA methylation at *F2RL3* (CpG_3 / cg03636183). ‘Fatal AMI/IHD/CVD’ – cases include only those with AMI/IHD/CVD recorded as a primary or secondary cause of death (those with AMI/IHD/CVD recorded as a main or secondary diagnosis from a hospital episode are excluded).

**Fig. S10. MR Result: Forest plot of odds ratios for CVD outcomes for each standard deviation decrease in *F2RL3* DNA methylation – UK Biobank sensitivity analyses.**

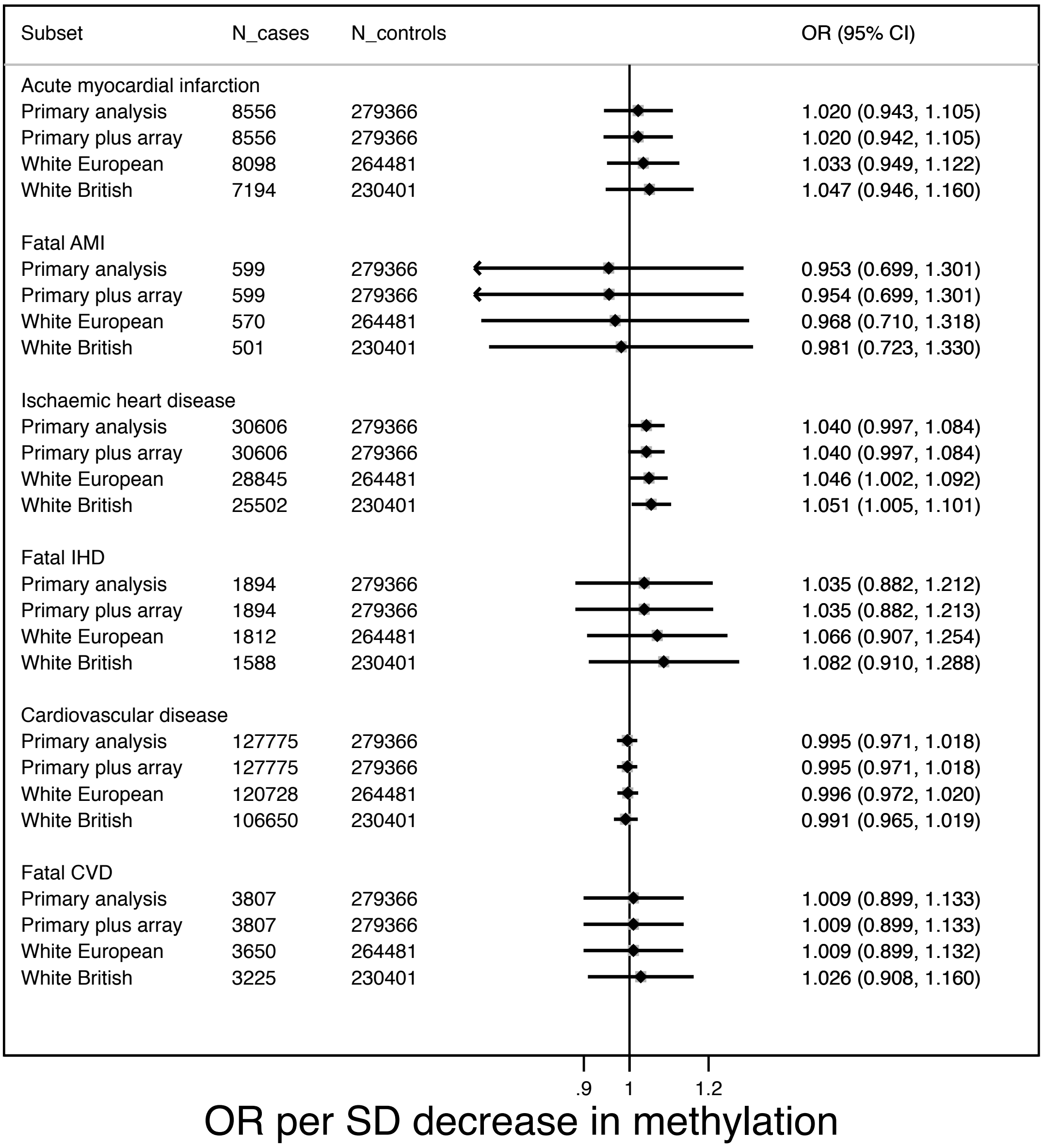

Results in UK Biobank shown for the multi-SNP instrument. *Primary analysis:* results from the principal analysis, as presented in the main manuscript; *Primary plus array:* results from an analysis in which genotyping array (UK Biobank or UK BiLEVE) was fitted alongside sex and principal components in the model for SNP-outcome association; *White European:* results from an analysis conducted in the White European subset of UK Biobank; *White British:* results from an analysis conducted in the White British subset of UK Biobank. Effect estimates represent the OR (95% CI) for each outcome per one standard deviation unit decrease in DNA methylation at *F2RL3* (CpG_3 / cg03636183).

**Fig. S11. MR Result: Forest plot of odds ratios for CVD outcomes for each standard deviation decrease in *F2RL3* DNA methylation by smoking status in UK Biobank using a multi-SNP instrument**

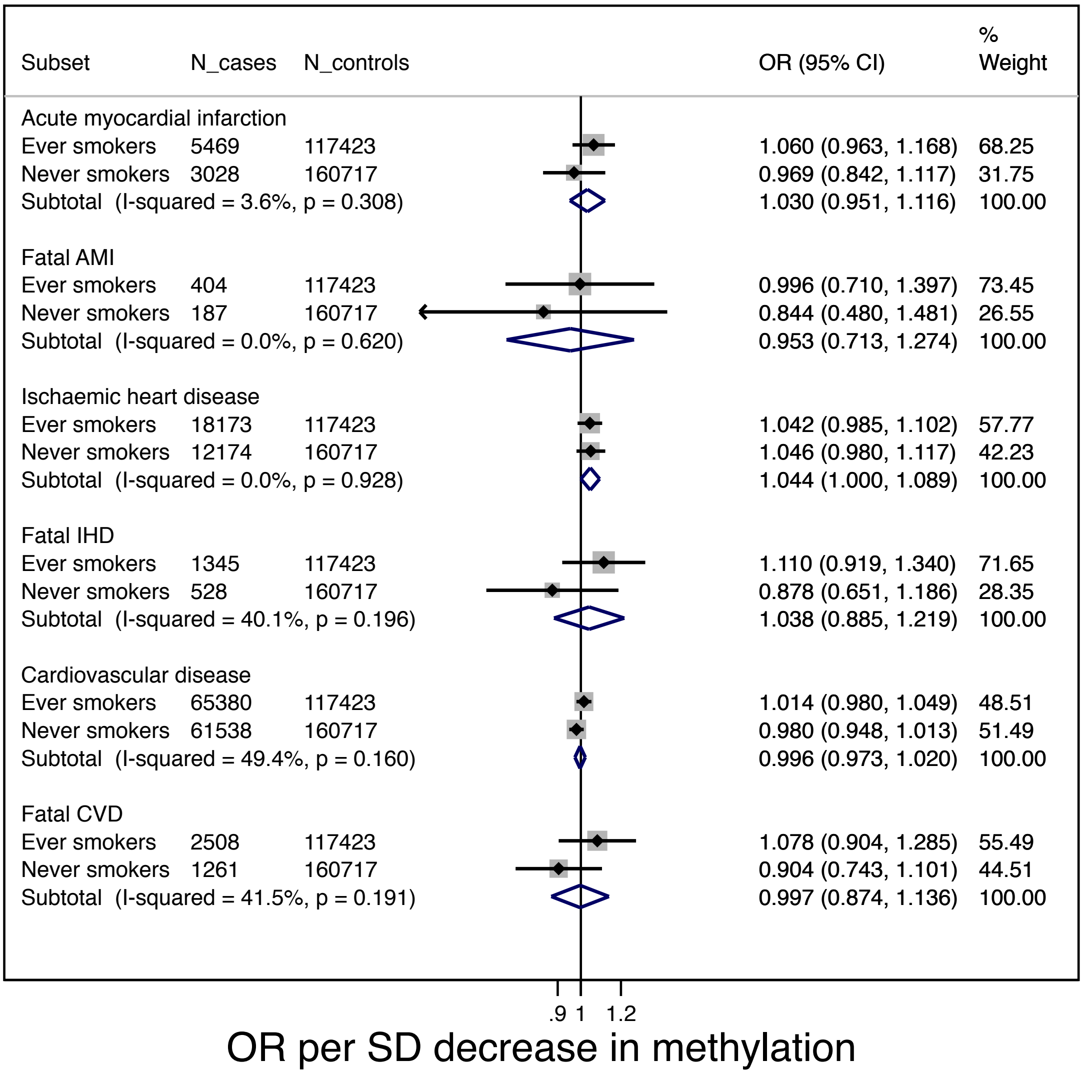

Results shown for the multi-SNP instrument. Effect estimates represent the OR (95% CI) for each outcome per one standard deviation unit decrease in DNA methylation at *F2RL3* (CpG_3 / cg03636183).

**Fig. S12. MR Result: Forest plot of odds ratios for CVD outcomes for each standard deviation decrease in *F2RL3* DNA methylation by smoking status in UK Biobank using rs773902 as an instrument**

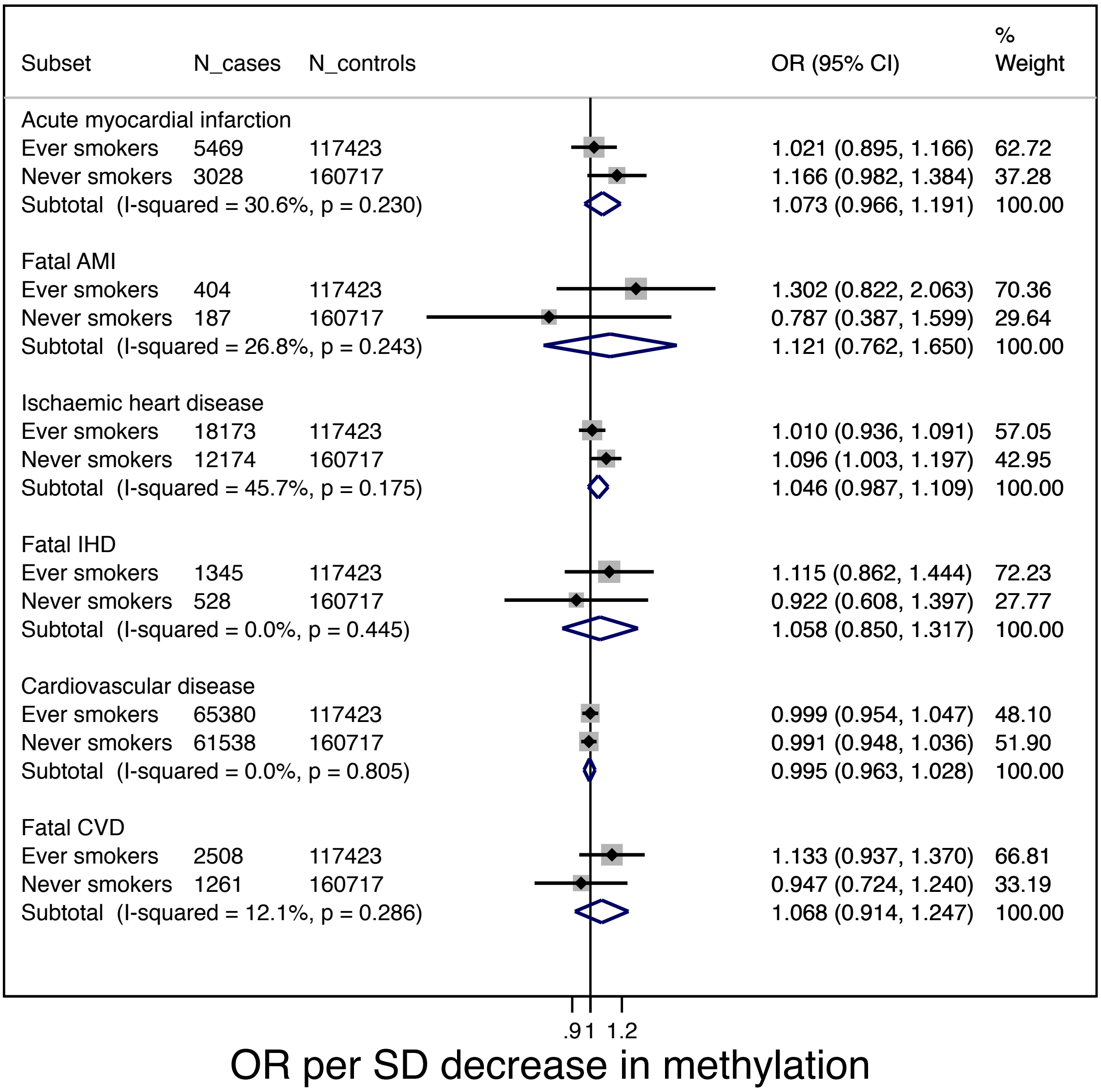

Results shown for rs773902 as the instrument. Effect estimates represent the OR (95% CI) for each outcome per one standard deviation unit decrease in DNA methylation at *F2RL3* (CpG_3 / cg03636183).

**Fig. S13. MR Result: Forest plot of odds ratios for CVD outcomes for each standard deviation decrease in *F2RL3* DNA methylation – CARDIoGRAMplusC4D sensitivity analysis.**

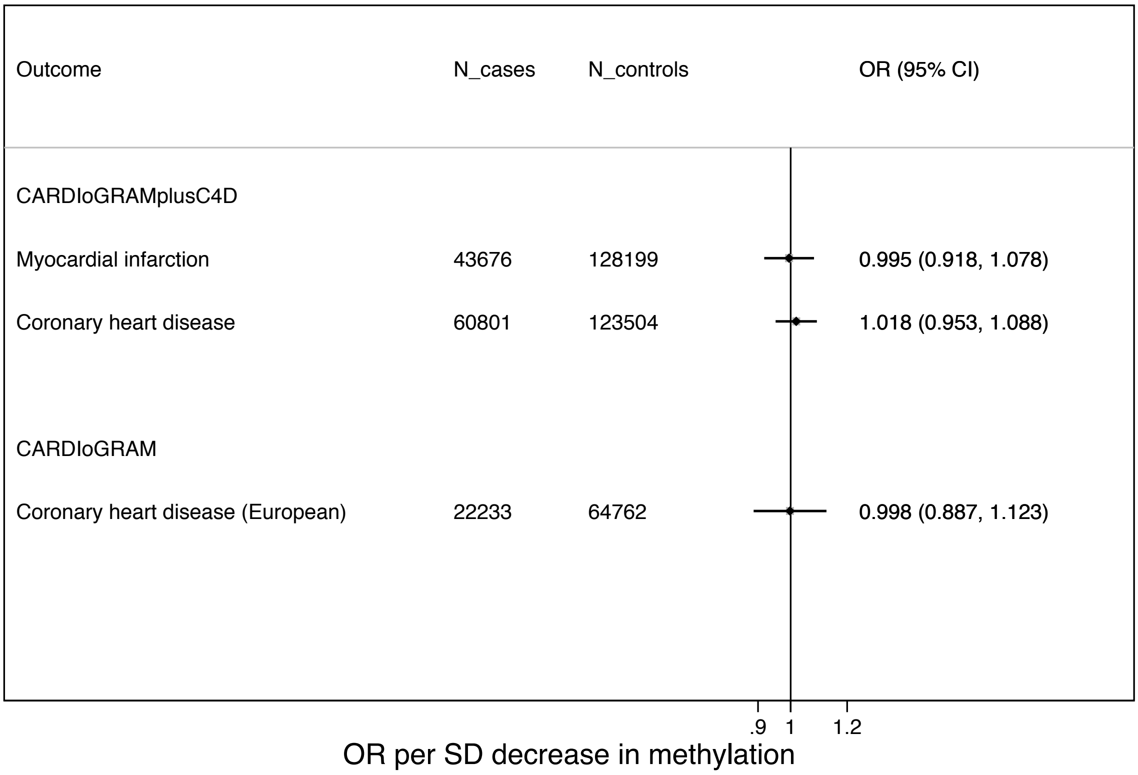

Results in CARDIoGRAM shown for the multi-SNP instrument, as compared to the results in CARDIoGRAMplusC4D. Effect estimates represent the OR (95% CI) for each outcome per one standard deviation unit decrease in DNA

**TABLES**

**Table S1. Characteristics of study population in the Copenhagen City Heart Study**

| | **With *F2RL3* DNA methylation data** | | | **Without *F2RL3* DNA methylation data** | | | **P-value** |
|---|---|---|---|---|---|---|---|
| | **N** | | | **N** | | | |
| Age (years) | 3,205 | **Mean (SD)** | 65.5 (11.0) | 1,087 | **Mean (SD)** | 65.3 (11.3) | 0.58 |
| Males | 3,205 | **N (%)** | 1,447 (45.2) | 1,087 | **N (%)** | 521 (47.9) | 0.11 |
| Acute myocardial infarction | 3,205 | **N (%)** | 853 (26.6) | 1,087 | **N (%)** | 272 (25.0) | 0.30 |
| Ischaemic heart disease | 3,205 | **N (%)** | 1,321 (41.2) | 1,087 | **N (%)** | 434 (39.9) | 0.46 |
| **Smoking status** | 3,205 | | | 1,087 | | | |
| Never | | **N (%)** | 548 (17.1) | | **N (%)** | 241 (22.2) | <0.001 |
| Former | | **N (%)** | 1,068 (33.3) | | **N (%)** | 366 (33.7) | |
| Current | | **N (%)** | 1,589 (49.6) | | **N (%)** | 480 (44.2) | |
| Pack years (ever smokers) | 2,649 | **Median (IQR)** | 31.5 (20,45) | 844 | **Median (IQR)** | 31.3 (18.3, 45) | 0.69 |
| Cigarettes per day (current smokers) | 1,589 | **Median (IQR)** | 15 (10,20) | 480 | **Median (IQR)** | 15 (11,20) | 0.28 |
| ***AHRR* DNA methylation intensity (%)** | 3,205 | **Mean (SD)** | 56.0 (8.9) | 1,087 | **Mean (SD)** | 57.1 (9.6) | <0.001 |
| ***F2RL3* DNA methylation intensity (%)** | 3,205 | | | | | | |
| CpG_1 | | **Mean (SD)** | 68.1 (17.1) | | | | |
| CpG_2 | | **Mean (SD)** | 89.6 (6.0) | | | | |
| CpG_3 | | **Mean (SD)** | 73.9 (11.1) | | | | |
| CpG_4 | | **Mean (SD)** | 78.3 (11.8) | | | | |

**Table S2. Associations between smoking and *F2RL3* DNA methylation in the Copenhagen City Heart Study.**

| | N | % DNA methylation *F2RL3* CpG_1 (95% CI) | $R^2$ | % DNA methylation *F2RL3* CpG_2 (95% CI) | $R^2$ | % DNA methylation *F2RL3* CpG_3 (95% CI) | $R^2$ | % DNA methylation *F2RL3* CpG_4 (95% CI) | $R^2$ |
|---|---|---|---|---|---|---|---|---|---|
| **Never** | 548 | - | 0.36 | | 0.20 | | 0.29 | | 0.28 |
| **Former** | 1,068 | -7.15 (-8.58, -5.71) | | -1.18 (-1.75, -0.61) | | -3.23 (-4.21, -2.25) | | -3.50 (-4.56, -2.45) | |
| **Current** | 1,589 | -23.98 (-25.33, -22.62) | | -5.92 (-6.46, -5.38) | | -13.4 (-14.40, -12.60) | | -14.12 (-15.12, -13.13) | |
| **Pack years in ever smokers** | 2,649 | -0.23 (-0.25, -0.20) | 0.12 | -0.06 (-0.07, -0.05) | 0.06 | -0.13 (-0.14, -0.11) | 0.09 | -0.14 (-0.16, -0.12) | 0.09 |
| **Cigarettes per day in current smokers** | 1,589 | -0.28 (-0.36, -0.20) | 0.04 | -0.10 (-0.13, -0.06) | 0.03 | -0.17 (-0.23, -0.12) | 0.03 | -0.20 (-0.26, -0.14) | 0.03 |
| **DNA methylation at AHRR** | 3,205 | 1.24 (1.19, 1.29) | 0.44 | 0.33 (0.31, 0.35) | 0.25 | 0.72 (0.69, 0.76) | 0.35 | 0.76 (0.72, 0.80) | 0.34 |

Analyses are adjusted for age and sex.

**Table S3A. Associations between *F2RL3* DNA methylation at CpG_1 and acute myocardial infarction in the Copenhagen City Heart Study.**

| Exposure (DNA methylation /Smoking) | N | N events | OR (95% CI) | | | |
|---|---|---|---|---|---|---|
| | | | **Age, sex adjusted** | **Age, sex, smoking status adjusted** | **Age, sex, smoking status, passive smoking adjusted** | **Age, sex, smoking status, passive smoking, *AHRR* DNA methylation adjusted** |
| ***F2RL3* (CpG_1) (in full sample) (per SD decrease)** | 2,998 | 646 | 1.33 (1.21, 1.45) | 1.23 (0.85, 1.53) | 1.22 (1.10, 1.36) | 1.25 (1.11, 1.42) |
| | | | **Age, sex adjusted** | **Age, sex, pack years adjusted** | **Age, sex, pack years, passive smoking adjusted** | **Age, sex, smoking status, pack years, *AHRR* DNA methylation adjusted[b]** |
| ***F2RL3* (CpG_1) (in ever smokers) (per SD decrease)[a]** | 2,465 | 561 | 1.29 (1.17, 1.42) | 1.29 (1.16, 1.43) | 1.27 (1.14, 1.41) | 1.27 (1.11, 1.44) |
| | | | **Age, sex adjusted** | **Age, sex, passive smoking adjusted** | - | **Age, sex, passive smoking, AHRR DNA methylation adjusted[b]** |
| ***F2RL3* (CpG_1) (in never smokers) (per SD decrease)** | 527 | 83 | 0.90 (0.53,1.53) | 0.92 (0.54, 1.56) | - | 0.98 (0.57, 1.69) |

Odds ratios are per standard deviation (SD) decrease in DNA methylation. Individuals with AMI diagnosed prior to date of clinic at which blood for DNA methylation assay was taken were excluded from analysis (N=207)

[a] Smokers with missing data on pack years were excluded from the ever smoker analysis (N=6)

[b] P-value for heterogeneity between fully adjusted models in ever and never smokers = 0.36

**Table S3B. Associations between smoking and acute myocardial infarction in the Copenhagen City Heart Study.**

| Smoking status | N | N events | OR (95% CI) | | |
|---|---|---|---|---|---|
| | | | Age, sex, adjusted | Age, sex, *F2RL3* DNA methylation (CpG_1), adjusted | Age, sex, *AHRR* DNA methylation, adjusted |
| **Never** | 527 | 83 | - | - | - |
| **Former** | 981 | 184 | 1.24 (0.93, 1.65) | 1.14 (0.85, 1.53) | 1.21 (0.90, 1.62) |
| **Current** | 1,490 | 379 | 1.89 (1.44, 2.46) | 1.42 (1.04, 1.93) | 1.74 (1.27, 2.37) |
| **Pack years in smokers[a]** | 2,465 | 561 | 1.004 (1.000, 1.008) | 1.000 (0.996, 1.005) | 1.002 (0.997, 1.006) |
| **Cigarettes per day (current smokers)** | 1,490 | 379 | 1.000 (0.988, 1.013) | 0.997 (0.984, 1.010) | 0.999 (0.986, 1.008) |
| **DNA methylation at *AHRR* (per 10% decrease)** | 2,998 | 646 | 1.24 (1.18, 1.38) | 1.00 (0.88, 1.15) | - |

Odds ratios are per standard deviation (SD) decrease in DNA methylation. Individuals with AMI diagnosed prior to date of clinic at which blood for DNA methylation assay was taken were excluded from analysis (N=207)

[a]Smokers with missing data on pack years were excluded from the pack years analysis (N=6)

**Table S4A. Associations between *F2RL3* DNA methylation at CpG_1 and mortality in individuals experiencing an AMI in the Copenhagen City Heart Study.**

| Exposure (DNA S5methylation /Smoking) | N | N events | Person years | HR (95% CI) | | | |
|---|---|---|---|---|---|---|---|
| | | | | Age, sex adjusted | Age, sex, smoking status adjusted | Age, sex, smoking status, passive smoking adjusted | Age, sex, smoking status, passive smoking, *AHRR* DNA methylation adjusted |
| ***F2RL3*** **(CpG_1) (in full sample) (per SD decrease)** | 646 | 506 | 2,934 | 1.50 (1.36, 1.66) | 1.49 (1.34, 1.67) | 1.49 (1.33, 1.66) | 1.39 (1.22, 1.58) |
| | | | | **Age, sex adjusted** | **Age, sex, pack years adjusted** | **Age, sex, pack years, passive smoking adjusted** | **Age, sex, smoking status, pack years, *AHRR* DNA methylation adjusted[b]** |
| ***F2RL3*** **(CpG_1) (in ever smokers) (per SD decrease)[a]** | 561 | 441 | 2,522 | 1.55 (1.39, 1.73) | 1.52 (1.36, 1.70) | 1.51 (1.34, 1.69) | 1.43 (1.23, 1.65) |
| | | | | **Age, sex adjusted** | **Age, sex, passive smoking adjusted** | - | **Age, sex, passive smoking, AHRR DNA methylation adjusted[b]** |
| ***F2RL3*** **(CpG_1) (in never smokers) (per SD decrease)** | 83 | 64 | 393 | 0.91 (0.56, 1.47) | 0.90 (0.56, 1.45) | - | 0.90 (0.56, 1.45) |

HR: Hazard ratios from Cox regression are per standard deviation (SD) decrease in DNA methylation, adjusted for age and sex, with date of clinic as the entry point. No clear evidence was found for violation of the proportional hazards assumption (P all > 0.21) Individuals with AMI diagnosed prior to date of clinic at which blood for DNA methylation assay was taken were excluded from analysis (N=207).

[a]Smokers with missing data on pack years excluded from the pack years analysis (N=2)

[b]P-value for heterogeneity between fully adjusted models in ever and never smokers = 0.07

**Table S4B. Associations between smoking and mortality in individuals experiencing an AMI in the Copenhagen City Heart Study.**

| Smoking status | N | N events | Person years | HR (95% CI) | | |
|---|---|---|---|---|---|---|
| | | | | Age, sex, adjusted | Age, sex, *F2RL3* DNA methylation (CpG_1), adjusted | Age, sex, *AHRR* DNA methylation, adjusted |
| **Never** | 83 | 64 | 393 | - | - | - |
| **Former** | 184 | 143 | 779 | 1.22 (0.90, 1.65) | 1.01 (0.74, 1.38) | 0.99 (0.73, 1.36) |
| **Current** | 379 | 299 | 1,762 | 1.65 (1.25, 2.19) | 1.03 (0.75, 1.42) | 1.10 (0.80, 1.52) |
| **Pack years in smokers[a]** | 561 | 441 | 2,522 | 1.007 (1.003, 1.012) | 1.003 (0.999, 1.008) | 1.006 (1.001, 1.010) |
| **Cigarettes per day (current smokers)** | 379 | 299 | 1,762 | 1.015 (1.001, 1.023) | 1.005 (0.990, 1.021) | 1.010 (0.996, 1.025) |
| **DNA methylation at *AHRR* (per 10% decrease)** | 646 | 506 | 2,934 | 1.45 (1.30, 1.63) | 1.15 (1.00, 1.33) | - |

HR: Hazard ratios from Cox regression, adjusted for age and sex, with date of clinic as the entry point. No clear evidence was found for violation of the proportional hazards assumption (P all > 0.11). Individuals with AMI diagnosed prior to date of clinic at which blood for DNA methylation assay was taken were excluded from analysis (N=207)

[a]Smokers with missing data on pack years were excluded from the pack years analysis (N=2)

**Table S5. Characteristics of ALSPAC participants selected for invite (n=200):** Values presented are mean (SD) unless specified otherwise.

| | N | Low DNA methylation | N | High DNA methylation | Age range of measure (years) | P-value* |
|---|---|---|---|---|---|---|
| **Selection Criteria** | | | | | | |
| ***F2RL3* DNA methylation [b] (%) during childhood** | 100 | 65.3 (0.04) | 100 | 74.2 (0.04) | 7.1-8.8 | <0.0001 |
| ***F2RL3* DNA methylation [b] (%) during adolescence** | 100 | 62.9 (0.04) | 100 | 71.5 (0.04) | 15.3-19.3 | <0.0001 |
| **Potential confounders** | | | | | | |
| **Sex (% male)** | 100 | 54% | 100 | 49% | n/a | 0.48[a] |
| **Age (years)** | 100 | 23.0 (0.5) | 100 | 23.2 (0.5) | 22.4-24.0 | 0.04 |
| **Maternal education (% with A-level/Degree)** | 99 | 46.5% | 99 | 52.5% | n/a | 0.39[a] |
| **BMI (at age 18 clinic)** | 89 | 22.3 (3.8) | 87 | 22.9 (3.4) | 16.5-19.3 | 0.76 |
| **Systolic blood pressure (at age 18 clinic)** | 86 | 119.6 (10.5) | 86 | 119.2 (10.2) | 16.5-19.3 | 0.81 |
| **Diastolic blood pressure (at age 18 clinic)** | 86 | 63.4 (5.8) | 86 | 63.2 (5.9) | 16.5-19.3 | 0.76 |
| **C-reactive protein (age 9)[e]** | 77 | 0.23(0.18,0.29) | 79 | 0.24 (0.18, 0.30) | 9.2-10.7 | 0.95 |
| **Interleukin 6 (age 9)[e]** | 77 | 0.74 (0.61, 0.89) | 79 | 0.76(0.63, 0.93) | 9.2-10.7 | 0.82 |
| **Cholesterol (age 9)** | 77 | 4.3 (0.5) | 79 | 4.3 (0.7) | 9.2-10.7 | 0.88 |
| **C-reactive protein (age 15)[e]** | 75 | 0.48 (0.38, 0.60) | 84 | 0.46 (0.37, 0.58) | 15.0-16.9 | 0.86 |
| **Cotinine (age 15)[c]** | 74 | 0.87 (0.31,1.27) | 84 | 0.76 (0.16, 1.12) | 15.0-16.9 | 0.68[d] |
| **Cholesterol (age 15)** | 75 | 3.7 (0.6) | 84 | 3.7 (0.6) | 15.0-16.9 | 0.50 |
| ***AHRR* DNA methylation [f] (%) during childhood** | 100 | 82.8 (0.03) | 100 | 83.3 (0.04) | 7.1-8.8 | 0.37 |
| ***AHRR* DNA methylation [f] (%) during adolescence** | 100 | 81.7 (0.04) | 100 | 81.9 (0.04) | 15.3-19.3 | 0.70 |
| **Genetic data** | | | | | | |
| **Minor allele (A) frequency at rs773902** | 100 | 0.34 | 100 | 0.08 | n/a | <0.0001[a] |

*P-value from two-sample t test unless specified otherwise

[a] Chi-square p-value

[b]DNA methylation at cg03636183 assessed by Illumina Infinium Human Methylation 450K BeadChip assay and presented as raw betas (proportion methylated) x 100 to give % DNA methylation.

[c]Median (interquartile range)

[d]Kruskal-Wallis equality-of-population rank test

[e]Log transformation performed prior to test. Geometric means presented.

[f]DNA methylation at cg05575921 assessed by Illumina Infinium Human Methylation 450K BeadChip assay and presented as raw betas (proportion methylated) x 100 to give % DNA methylation.

**Table S6. Comparison of recruited sample of ALSPAC participants (n=41) and all those selected for invite (n=200):** Values presented are mean (SD) unless specified otherwise

| | N | Total eligible | N | Recruited | Age range of measure (years) | P-value* |
|---|---|---|---|---|---|---|
| **Selection criteria** | | | | | | |
| ***F2RL3* DNA methylation [b] (%) during childhood** | 200 | 69.8 (0.06) | 41 | 69.7 (0.06) | 7.1-8.8 | 0.95 |
| ***F2RL3* DNA methylation [b] (%) during adolescence** | 200 | 67.2 (0.06) | 41 | 68.8 (0.07) | 15.3-19.3 | 0.06 |
| **Potential confounders** | | | | | | |
| **Sex (% male)** | 200 | 51.5% | 41 | 41.5% | n/a | 0.15[a] |
| **Age (years)** | 200 | 23.1 (0.5) | 41 | 23.1 (0.5) | 22.4-24.0 | 0.42 |
| **Maternal education (% with A-level/Degree)** | 198 | 49.5% | 41 | 53.7% | n/a | 0.55[a] |
| **BMI (at age 18 clinic)** | 176 | 22.6 (3.6) | 38 | 22.3 (3.8) | 16.5-18.3 | 0.58 |
| **Systolic blood pressure (at age 18 clinic)** | 172 | 119.4 (10.3) | 38 | 119.1 (11.5) | 16.5-18.3 | 0.86 |
| **Diastolic blood pressure (at age 18 clinic)** | 172 | 63.3 (5.8) | 38 | 64.3 (6.7) | 16.5-18.3 | 0.21 |
| **C-reactive protein (age 9)[e]** | 156 | 0.23 (0.20, 0.28) | 29 | 0.27 (0.19, 0.38) | 9.4- 10.0 | 0.47 |
| **Interleukin 6 (age 9)[e]** | 156 | 0.75 (0.66, 0.86) | 29 | 0.68 (0.46, 1.02) | 9.4- 10.0 | 0.50 |
| **Cholesterol (age 9)** | 156 | 4.3 (0.6) | 29 | 4.4 (0.6) | 9.4- 10.0 | 0.22 |
| **C-reactive protein (age 15)[e]** | 159 | 0.47 (0.40, 0.55) | 33 | 0.47 (0.33, 0.67) | 15.1- 15.6 | 0.98 |
| **Cotinine (age 15)[c]** | 158 | 0.80 (0.31, 1.14) | 33 | 0.70 (0.35, 0.93) | 15.1- 15.6 | 0.20[d] |
| **Cholesterol (age 15)** | 159 | 3.7 (0.6) | 33 | 3.8 (0.5) | 15.1- 15.6 | 0.28 |
| ***AHRR* DNA methylation [f] (%) during childhood** | 200 | 83.0 (0.03) | 41 | 82.7 (0.03) | 7.1-8.8 | 0.42 |
| ***AHRR* DNA methylation [f] (%) during adolescence** | 200 | 81.8 (0.04) | 41 | 82.3 (0.04) | 15.3-19.3 | 0.42 |
| **Genetic data** | | | | | | |
| **Minor allele (A) frequency at rs773902** | 200 | 0.21 | 41 | 0.26 | n/a | 0.22[a] |

*P-value from two-sample t test comparing recruited to non-recruited participants unless specified otherwise

[a] Chi-square p-value

[b]DNA methylation at cg03636183 assessed by Illumina Infinium Human Methylation 450K BeadChip assay and presented as raw betas (proportion methylated) x 100 to give % DNA methylation.

[c] Median (interquartile range)

[d]Kruskal-Wallis equality-of-population rank test

[e]Log transformation performed prior to test. Geometric means presented.

[f]DNA methylation at cg05575921 assessed by Illumina Infinium Human Methylation 450K BeadChip assay and presented as raw betas (proportion methylated) x 100 to give % DNA methylation.

**Table S7. Contemporary DNA methylation assessed by targeted pyrosequencing in the recruited sample of ALSPAC participants (n=41*):** Values presented are mean (SD)

| | N | Low DNA methylation | N | High DNA methylation | Age range of measure | P-value[a] |
|---|---|---|---|---|---|---|
| **CpG_1** | 17 | 83.2 (3.3) | 16 | 88.0 (3.9) | 22.5-24.4 | **$8.0 \times 10^{-4}$** |
| **CpG_2** | 18 | 91.4 (0.6) | 17 | 91.9 (0.8) | 22.5-24.4 | **$4.4 \times 10^{-2}$** |
| **CpG_3** | 18 | 82.1 (1.8) | 17 | 84.9 (1.9) | 22.5-24.4 | **$2.0 \times 10^{-4}$** |
| **CpG_4** | 17 | 83.4 (2.1) | 13 | 85.1 (2.3) | 22.5-24.4 | **$4.2 \times 10^{-2}$** |

* Total N in table does not sum to 41 because of assay failures.

[a]P-value from two-sample Wilcoxon rank-sum (Mann-Whitney) test

**Table S8. Correlation across CpG sites in *F2RL3* of contemporary DNA methylation assessed by targeted pyrosequencing**

| | CpG_1 | CpG_2 | CpG_3 |
|---|---|---|---|
| **CpG_2** | 0.50 | | |
| **CpG_3** | 0.80 | 0.41 | |
| **CpG_4** | 0.69 | 0.61 | 0.72 |

**Table S9. By-group comparison of platelet reactivity measures amongst ALSPAC participants:** Values presented are mean (SD). Comparisons made after the removal of outliers identified by robust regression of outcome measure on DNA methylation group (high/low).

| Assay | Measure | N | Low DNA methylation | N | High DNA methylation | P-value[a] |
|---|---|---|---|---|---|---|
| **Stimulation by PAR4 agonist (AYPGKF) (assessed by flow cytometry)** | **Integrin activation ($EC_{50}$) (μM)** | 19 | 60.8 (12.0) | 22 | 80.1 (20.5) | **$9.9 \times 10^{-4}$** |
| | **Integrin activation (Max. response) (MFI)** | 19 | 4,480 (1,005) | 22 | 4,634 (1,128) | 0.68 |
| | **P-selectin exposure ($EC_{50}$) (μM)** | 18 | 96.7 (25.1) | 21 | 119.4 (38.6) | **$4.6 \times 10^{-2}$** |
| | **P-selectin exposure (Max. response) (MFI)** | 19 | 3,711 (1390) | 22 | 3,847 (1,191) | 0.83 |
| **Surface receptor expression (assessed by flow cytometry)** | **CD41 (MFI)** | 19 | 23,720 (8,195) | 22 | 23,216 (9,482) | 0.88 |
| | **CD61 (MFI)** | 19 | 14,511 (2,662) | 22 | 14,311 (3,793) | 0.90 |
| **Stimulation by PAR1 agonist (SFLLRN)** | **Integrin activation ($EC_{50}$) (μM)** | 17 | 1.79 (0.73) | 21 | 1.95 (0.74) | 0.36 |
| | **P-selectin exposure ($EC_{50}$) (μM)** | 17 | 2.73 (1.14) | 22 | 2.84 (1.11) | 0.48 |

$EC_{50}$ = half maximum concentration values (the concentration of agonist required to give half the maximal response); MFI = (geometric) mean fluorescence intensity

[a]P-value from two-sample Wilcoxon rank-sum (Mann-Whitney) test

**Table S10. By-group comparison of haematological measures amongst ALSPAC participants:** Values presented are mean (SD)

| Measure | N | Low DNA methylation | N | High DNA methylation | P-value* |
|---|---|---|---|---|---|
| **WBC ($10^3$/mm$^3$)** | 18 | 5.06 (1.09) | 22 | 5.04 (1.23) | 0.95 |
| **RBC ($10^3$/mm$^3$)** | 19 | 4.18 (0.29) | 22 | 4.10 (0.29) | 0.47 |
| **PLT ($10^3$/mm$^3$)** | 19 | 194.3 (38.6) | 22 | 210.8 (47.0) | 0.32 |
| **MPV (μm$^3$)** | 19 | 7.75 (0.74) | 22 | 7.62 (0.64) | 0.50 |

WBC= white blood cells, RBC= red blood cells, PLT= platelets, MPV= mean platelet volume. *P-value from two-sample Wilcoxon rank-sum (Mann-Whitney) test

**Table S11. Characteristics of recruited sample amongst ALSPAC participants (n=41):** Values presented are mean (SD) unless specified otherwise.

| | N | Low DNA methylation | N | High DNA methylation | Age range of measure (years) | P-value* |
|---|---|---|---|---|---|---|
| **Selection Criteria** | | | | | | |
| ***F2RL3* DNA methylation [b] (%) during childhood** | 19 | 64.3 (0.04) | 22 | 74.3 (0.03) | 7.4-7.7 | <0.0001 |
| ***F2RL3* DNA methylation [b] (%) during adolescence** | 19 | 63.3 (0.04) | 22 | 73.5 (0.04) | 15.4-18.3 | <0.0001 |
| **Potential confounders** | | | | | | |
| **Sex (% male)** | 19 | 36.8 % | 22 | 45.5% | n/a | 0.58/0.75[a] |
| **Age (years)** | 19 | 23.4 (0.6) | 22 | 23.4 (0.5) | 22.5-24.4 | 0.95 |
| **Maternal education (% with A-level/Degree)** | 19 | 36.8% | 22 | 68.2% | n/a | 0.05/0.06 [a] |
| **BMI (age 18)** | 18 | 22.8 (4.2) | 20 | 21.8 (3.5) | 16.5-18.3 | 0.46 |
| **Systolic blood pressure (age 18)** | 19 | 119.0 (8.4) | 19 | 119.2 (14.2) | 16.5-18.3 | 0.97 |
| **Diastolic blood pressure (age 18)** | 19 | 65.7 (7.4) | 19 | 63.0 (5.7) | 16.5-18.3 | 0.21 |
| **C-reactive protein (age 9)[e]** | 14 | 0.28 (0.16, 0.49) | 15 | 0.25 (0.15, 0.42) | 9.4- 10.0 | 0.79 |
| **Interleukin 6 (age 9)[e]** | 14 | 0.61 (0.27, 1.36) | 15 | 0.76 (0.55, 1.06) | 9.4- 10.0 | 0.57 |
| **Cholesterol (age 9)** | 14 | 4.33 (0.48) | 15 | 4.44 (0.72) | 9.4- 10.0 | 0.64 |
| **C-reactive protein (age 15)[e]** | 17 | 0.48 (0.27, 0.85) | 16 | 0.45 (0.27, 0.50) | 15.1- 15.6 | 0.86 |
| **Cotinine (age 15)[c]** | 17 | 0.71 (0.36, 1.03) | 16 | 0.60 (0.36, 0.89) | 15.1- 15.6 | 0.70[d] |
| **Cholesterol (age 15)** | 17 | 3.74 (0.42) | 16 | 3.88 (0.61) | 15.1- 15.6 | 0.47 |
| ***AHRR* DNA methylation [f] (%) during childhood** | 19 | 82.0 (0.03) | 22 | 83.2 (0.03) | 7.4-7.7 | 0.25 |
| ***AHRR* DNA methylation [f] (%) during adolescence** | 19 | 82.2 (0.03) | 22 | 82.3 (0.05) | 15.4-18.3 | 0.90 |
| **Laboratory technician (% technician 1)[g]** | 19 | 57.9 | 22 | 27.3 | n/a | 0.05/0.06[a] |
| **Time to assay (minutes)[h]** | 19 | 41.9 (15.6) | 22 | 31.9 (9.3) | n/a | 0.01 |
| **Genetic data** | | | | | | |
| **Minor allele (A) frequency at rs773902** | 19 | 0.47 | 22 | 0.07 | n/a | <0.0001[a] |

*P-value from two-sample t test unless specified otherwise

[a]Chi-square p-value/Fisher's exact p-value (if different)

[b]DNA methylation at cg03636183 assessed by Illumina Infinium Human Methylation 450K BeadChip assay and presented as raw betas (proportion methylated) x 100 to give % DNA methylation.

[c]Median (interquartile range)

[d]Kruskal-Wallis equality-of-population rank test

[e]Log transformation performed prior to test. Geometric means presented.

[f]DNA methylation at cg05575921 assessed by Illumina Infinium Human Methylation 450K BeadChip assay and presented as raw betas (proportion methylated) x 100 to give % DNA methylation.

[g]Analyses were performed by two laboratory technicians

[h]Represents the time lag between blood draw and the start of the laboratory analysis (sample put into haematology analyser). In cases where the blood draw time was not recorded, draw time was predicted based on the mean time from appointment start time to blood draw (9 minutes).

**Table S12. Associations between rs1051730 and amount of smoking amongst participants in the Copenhagen City Heart Study**

| | N | Beta (95% CI) | P-value |
|---|---|---|---|
| **Cigarettes per day in current smokers** | 2,065 | 0.83 (0.24, 1.43) | 0.006 |
| **Pack years in ever smokers** | 3,487 | 0.79 (-0.34, 1.92) | 0.17 |

Associations are per minor allele of rs1051730. All analyses are age, sex adjusted.

**Table S13. Associations between rs1051730 and *F2RL3* DNA methylation amongst participants in the Copenhagen City Heart Study**

| | N | Beta (95% CI) | P-value | P-value for heterogeneity |
|---|---|---|---|---|
| **CpG_1** | | | | |
| **Never smokers** | 548 | -0.56 (-1.58, 0.46) | 0.29 | |
| **Former smokers** | 1,066 | -1.26 (-2.48, -0.07) | 0.04 | |
| **Current smokers** | 1,585 | -1.57 (-2.71, -0.43) | 0.007 | 0.40[a] |
| **Ever smokers** | 2,651 | -1.55 (-2.50, -0.60) | 0.001 | 0.16[b] |
| **CpG_2** | | | | |
| **Never smokers** | 548 | -0.12 (-0.51, 0.27) | 0.54 | |
| **Former smokers** | 1,066 | -0.08 (-0.46, 0.29) | 0.66 | |
| **Current smokers** | 1,585 | -0.49 (-0.98, 0.005) | 0.05 | 0.40[a] |
| **Ever smokers** | 2,651 | -0.36 (-0.71, -0.001) | 0.05 | 0.38[b] |
| **CpG_3** | | | | |
| **Never smokers** | 548 | -0.08 (-0.78, 0.63) | 0.83 | |
| **Former smokers** | 1,066 | -0.52 (-1.23, 0.19) | 0.15 | |
| **Current smokers** | 1,585 | -1.10 (-1.93, -0.28) | 0.008 | 0.18[a] |
| **Ever smokers** | 2,651 | -0.93 (-1.57, -0.30) | 0.004 | 0.08[b] |
| **CpG_4** | | | | |
| **Never smokers** | 548 | -0.29 (-1.01, 0.44) | 0.34 | |
| **Former smokers** | 1,066 | -0.13 (-0.93, 0.66) | 0.74 | |
| **Current smokers** | 1,585 | -1.08 (-1.95, -0.19) | 0.02 | 0.25[a] |
| **Ever smokers** | 2,651 | -0.76 (-1.45, -0.09) | 0.03 | 0.34[b] |

Associations are per minor allele of rs1051730. All analyses are age, sex adjusted.
[a]P-value for heterogeneity between never, former and current smokers.
[b]P-value for heterogeneity between never and ever smokers.

**Table S14. Meta-analysis results from the GoDMC consortium (N=27,750) for genetic variants on chromosome 19 used as instruments for *F2RL3* DNA methylation. Shaded variants were not included in the multi-SNP instrument.**

| rsID | Base pair position[a] | Effect allele[b] | Other allele | EAF[c] | Beta[d] (SE) | P-value | N | $I^2$ (Q *P*-value[e])[f] |
|---|---|---|---|---|---|---|---|---|
| rs3848657 | 16950821 | G | A | 0.14 | -0.28 (0.01) | 0 | 24,236 | 58.9 (2.5e-05) |
| rs2227349 | 17001551 | G | A | 0.66 | -0.39 (0.01) | 0 | 23,227 | 61 (3.9e-06) |
| rs55961639 | 16975870 | G | C | 0.83 | -0.18 (0.01) | 7.1e-55 | 27,733 | 38.0 (4.2e-22) |
| rs112791293 | 16920006 | T | C | 0.06 | -0.28 (0.03) | 8.6e-27 | 13,897 | 28.2 (0.13) |
| rs773919[g] | 16854486 | T | A | 0.45 | -0.08 (0.01) | 3.4e-17 | 22,488 | 2.1 (0.43) |
| rs189158451 | 16911878 | C | T | 0.98 | -0.35 (0.06) | 9.2e-08 | 6,133 | 37.1 (0.12) |
| rs812847[h] | 16855265 | G | A | 0.45 | -0.07 (0.01) | 8.4e-17 | 25,262 | 1.4 (0.44) |
| rs773902[i] | 17000632 | A | G | 0.19 | -0.37 (0.01) | 1.9e-228 | 23,972 | 87.4 (2.2e-34) |

[a]Based on: Genome Reference Consortium Human Build 37 patch release 13 (GRCh37.p13)

[b]Set to be the DNA methylation decreasing allele

[c]EAF = effect allele frequency

[d]Effect expressed in standard deviation units

[e]P-value from Cochran's Q test

[f]The high $I^2$ values arise because the SNPs exhibit very large effects with very small confidence intervals. Though the majority of heterogeneity is due to between study variance, the absolute differences between those studies is very small.

[g]This SNP was palindromic and was therefore replaced by a proxy (rs812847) in the multi-SNP instrument

[h]This SNP was a proxy for rs773919

[i]This SNP was not part of the multi-SNP instrument

**Table S15. Linkage disequilibrium correlation matrix for SNPs in multi-SNP instrument***

| | rs11279129 | rs18915845 | rs2227349 | rs3848657 | rs55961639 | rs812847 |
|---|---|---|---|---|---|---|
| rs11279129 | 1.00 | | | | | |
| rs18915845 | 0.0352 | 1.00 | | | | |
| rs2227349 | 0.1641 | 0.1527 | 1.0000 | | | |
| rs3848657 | -0.0010 | 0.0450 | 0.1676 | 1.00 | | |
| rs55961639 | -0.2610 | 0.2491 | 0.3281 | 0.1846 | 1.00 | |
| rs812847 | -0.0004 | 0.0951 | 0.1838 | 0.2325 | 0.1249 | 1.00 |

*Calculated in UK Biobank (n=407,141) with respect to the methylation decreasing allele at each SNP (since this allele was used as the reference in the calculation of the SNP-exposure and SNP-outcome associations in the Mendelian randomization analysis).

**Table S16A. Association of genotype with potential confounders in entire UK Biobank cohort (after QC) (N=407,141)**

| | | *p*-value for association with genotype | | | | | | |
|---|---|---|---|---|---|---|---|---|
| | | **rs3848657** | **rs2227349** | **rs55961639** | **rs112791293** | **rs189158451** | **rs812847** | **rs773902** |
| **Genetic principal components** [a] | **1** | < 0.001 | < 0.001 | < 0.001 | < 0.001 | < 0.001 | < 0.001 | < 0.001 |
| | **2** | < 0.001 | < 0.001 | < 0.001 | < 0.001 | < 0.001 | < 0.001 | < 0.001 |
| | **3** | 0.001 | < 0.001 | < 0.001 | < 0.001 | 0.176 | < 0.001 | < 0.001 |
| | **4** | < 0.001 | 0.382 | 0.700 | < 0.001 | 0.636 | < 0.001 | < 0.001 |
| **Age**[a, b] | | 0.147 | < 0.001 | 0.512 | < 0.001 | 0.038 | < 0.001 | < 0.001 |
| **Daily cigarettes**[a, b, c] | | 0.187 | < 0.001 | 0.209 | < 0.001 | 0.704 | 0.003 | 0.022 |
| **Sex**[a] | | 0.165 | 0.730 | 0.670 | 0.562 | 0.490 | 0.106 | 0.797 |
| **Smoking status (ever/never)**[a] | | 0.163 | < 0.001 | 0.319 | < 0.001 | 0.024 | < 0.001 | < 0.001 |
| **Smoking status (never/former/current)** [b, d] | | 0.347 | < 0.0001 | 0.497 | < 0.0001 | 0.087 | < 0.0001 | < 0.0001 |
| **Highest employment status**[b, d] | | 0.412 | < 0.0001 | 0.331 | < 0.0001 | 0.412 | < 0.0001 | < 0.0001 |
| **Highest qualification**[b, d] | | 0.0007 | 0.0001 | 0.0009 | < 0.0001 | 0.592 | < 0.0001 | 0.0001 |

[a]*p*-value from linear regression;

[b]at first assessment;

[c]with those who smoked less than one cigarette per day given a value of -10;

[d]*p*-values from multinomial logistic regression.

**Table S16B. Association of genotype with potential confounders in a subset of White Europeans from the UK Biobank cohort (after QC) (N=385,209)**

| | | ***p*-value for association with genotype** | | | | | | |
|---|---|---|---|---|---|---|---|---|
| | | **rs3848657** | **rs2227349** | **rs55961639** | **rs112791293** | **rs189158451** | **rs812847** | **rs773902** |
| **Genetic principal components** [a] | **1** | < 0.001 | < 0.001 | 0.238 | < 0.001 | 0.007 | < 0.001 | < 0.001 |
| | **2** | < 0.001 | < 0.001 | 0.017 | 0.399 | 0.125 | < 0.001 | < 0.001 |
| | **3** | < 0.001 | < 0.001 | 0.302 | 0.026 | 0.001 | < 0.001 | < 0.001 |
| | **4** | < 0.001 | < 0.001 | 0.925 | 0.105 | 0.013 | < 0.001 | < 0.001 |
| **Age**[a, b] | | 0.505 | 0.661 | 0.788 | 0.282 | 0.594 | 0.009 | 0.876 |
| **Daily cigarettes**[a, b, c] | | 0.674 | 0.155 | 0.252 | 0.458 | 0.986 | 0.599 | 0.408 |
| **Sex**[a] | | 0.317 | 0.141 | 0.756 | 0.752 | 0.345 | 0.262 | 0.517 |
| **Smoking status (ever/never)**[a] | | 0.747 | 0.245 | 0.459 | 0.578 | 0.198 | 0.232 | 0.873 |
| **Smoking status (never/former/current)**[b, d] | | 0.429 | 0.474 | 0.607 | 0.649 | 0.496 | 0.195 | 0.098 |
| **Highest employment status**[b, d] | | 0.164 | 0.039 | 0.169 | 0.198 | 0.895 | 0.728 | 0.021 |
| **Highest qualification**[b, d] | | 0.260 | 0.558 | 0.021 | 0.084 | 0.694 | 0.001 | 0.313 |

[a]*p*-value from linear regression;

[b]at first assessment;

[c]with those who smoked less than one cigarette per day given a value of -10;

[d]*p*-values from multinomial logistic regression.

**Table S16C. Association of genotype with potential confounders in entire UK Biobank cohort (after QC) (N=407,141) after adjustment for ten principal components**

| | *p*-value for association with genotype | | | | | | |
|---|---|---|---|---|---|---|---|
| | **rs3848657** | **rs2227349** | **rs55961639** | **rs112791293** | **rs189158451** | **rs812847** | **rs773902** |
| **Age[a,b]** | 0.681 | 0.154 | 0.941 | 0.720 | 0.800 | 0.860 | 0.073 |
| **Daily cigarettes[a, b, c]** | 0.480 | 0.166 | 0.288 | 0.341 | 0.931 | 0.822 | 0.194 |
| **Sex[a]** | 0.447 | 0.104 | 0.987 | 0.802 | 0.350 | 0.175 | 0.618 |
| **Smoking status (ever/never)[a]** | 0.865 | 0.549 | 0.431 | 0.835 | 0.181 | 0.861 | 0.559 |
| **Smoking status (never/former/current)[b, d]** | 0.780 | 0.785 | 0.654 | 0.633 | 0.469 | 0.910 | 0.400 |
| **Highest employment status[b, d]** | 0.383 | 0.240 | 0.402 | 0.698 | 0.965 | 0.612 | 0.200 |
| **Highest qualification[b, d]** | 0.439 | 0.854 | 0.057 | 0.095 | 0.744 | 0.634 | 0.449 |

[a]*p*-value from linear regression;

[b]at first assessment;

[c]with those who smoked less than one cigarette per day given a value of -10;

[d]*p*-values from multinomial logistic regression.

### Table S17A GTEx portal results for *F2RL3* (last accessed: 04/07/2018)[1]

| Gencode Id | Gene | Variant Id | SNP Id | P-Value | NES[a] | Tissue |
|---|---|---|---|---|---|---|
| ENSG00000127533.3 | F2RL3 | 19_16997607_C_A_b37 | rs4346313 | 0.0000012 | -0.33 | Esophagus - Mucosa |
| ENSG00000127533.3 | F2RL3 | 19_16856864_C_T_b37 | rs773916 | 0.0000014 | 0.82 | Minor Salivary Gland |
| ENSG00000127533.3 | F2RL3 | 19_17011247_C_A_b37 | rs7257262 | 0.0000024 | 0.58 | Brain - Nucleus accumbens (basal |
| ENSG00000127533.3 | F2RL3 | 19_17005029_C_T_b37 | rs2227359 | 0.0000028 | 2 | Small Intestine - Terminal Ileum |
| ENSG00000127533.3 | F2RL3 | 19_17006066_C_CA_b37 | rs2227364 | 0.0000028 | 2 | Small Intestine - Terminal Ileum |
| ENSG00000127533.3 | F2RL3 | 19_17009157_A_AT_b37 | rs143120093 | 0.0000028 | 2 | Small Intestine - Terminal Ileum |
| ENSG00000127533.3 | F2RL3 | 19_16969365_C_T_b37 | rs62128036 | 0.0000029 | -0.32 | Esophagus - Mucosa |
| ENSG00000127533.3 | F2RL3 | 19_17006700_G_T_b37 | rs2981473 | 0.0000047 | 0.58 | Brain - Nucleus accumbens (basal |
| ENSG00000127533.3 | F2RL3 | 19_17224526_C_T_b37 | rs78352768 | 0.0000061 | -0.9 | Cells - EBV-transformed lymphocytes |
| ENSG00000127533.3 | F2RL3 | 19_16969855_T_C_b37 | rs2313234 | 0.0000067 | -0.31 | Esophagus - Mucosa |
| ENSG00000127533.3 | F2RL3 | 19_17006110_G_A_b37 | rs2227365 | 0.0000072 | 0.56 | Brain - Nucleus accumbens (basal |
| ENSG00000127533.3 | F2RL3 | 19_17006787_G_A_b37 | rs2227367 | 0.0000072 | 0.56 | Brain - Nucleus accumbens (basal |
| ENSG00000127533.3 | F2RL3 | 19_17011131_C_G_b37 | rs7254010 | 0.0000072 | 0.56 | Brain - Nucleus accumbens (basal |
| ENSG00000127533.3 | F2RL3 | 19_17003553_G_A_b37 | rs2227357 | 0.0000079 | -0.24 | Adipose - Subcutaneous |
| ENSG00000127533.3 | F2RL3 | 19_17000632_G_A_b37 | rs773902 | 0.000011 | -0.25 | Esophagus - Mucosa |
| ENSG00000127533.3 | F2RL3 | 19_16976289_T_C_b37 | rs2303091 | 0.000012 | -0.3 | Esophagus - Mucosa |
| ENSG00000127533.3 | F2RL3 | 19_17000632_G_A_b37 | rs773902 | 0.000018 | -0.19 | Adipose - Subcutaneous |
| ENSG00000127533.3 | F2RL3 | 19_17009372_A_G_b37 | rs2227371 | 0.000025 | -0.19 | Adipose - Subcutaneous |
| ENSG00000127533.3 | F2RL3 | 19_16966594_C_T_b37 | rs62128035 | 0.000026 | -0.3 | Esophagus - Mucosa |
| ENSG00000127533.3 | F2RL3 | 19_16950763_C_T_b37 | rs3848656 | 0.000033 | -0.3 | Esophagus - Mucosa |

[a]NES = normalized effect size

### Table S17B Blood eQTL browser results for *F2RL3* (last accessed: 04/07/2018)[2]

| *Trans*-eQTLs | | | | | | | | | | | |
|---|---|---|---|---|---|---|---|---|---|---|---|
| **P-value** | **SNP** | **SNP Chr.** | **SNP Chr. position** | **Probe** | **Probe Chr.** | **Probe Chr. position** | **SNP Alleles** | **Minor Allele** | **Z-score** | **Gene name** | **FDR** |
| 9.53E-06 | rs12485738 | 3 | 56840816 | 6250139 | 19 | 16863640 | G/A | A | -4.43 | F2RL3 | 0.39 |
| ***Cis*-eQTLs** | | | | | | | | | | | |
| **P-value** | **SNP** | **SNP Chr.** | **SNP Chr. Position** | **Probe** | **Probe Chr.** | **Probe Chr. position** | **SNP Alleles** | **Minor Allele** | **Z-score** | **Gene name** | **FDR** |
| 2.53E-07 | rs2608732 | 19 | 16869578 | 6250139 | 19 | 16863640 | G/C | C | -5.16 | F2RL3 | 0 |
| 2.67E-07 | rs773901 | 19 | 16864789 | 6250139 | 19 | 16863640 | T/G | G | -5.15 | F2RL3 | 0 |
| 1.79E-05 | rs10407613 | 19 | 16849786 | 6250139 | 19 | 16863640 | G/A | A | 4.29 | F2RL3 | 0.01 |
| 2.26E-05 | rs2227356 | 19 | 16864405 | 6250139 | 19 | 16863640 | C/T | T | -4.24 | F2RL3 | 0.01 |
| 2.79E-05 | rs1054533 | 19 | 16865049 | 6250139 | 19 | 16863640 | C/T | C | -4.19 | F2RL3 | 0.01 |
| 3.23E-05 | rs7245967 | 19 | 16872485 | 6250139 | 19 | 16863640 | T/C | C | -4.16 | F2RL3 | 0.01 |
| 5.06E-05 | rs11672791 | 19 | 16857200 | 6250139 | 19 | 16863640 | T/C | C | 4.05 | F2RL3 | 0.02 |
| 5.94E-05 | rs734568 | 19 | 16876685 | 6250139 | 19 | 16863640 | C/T | T | -4.02 | F2RL3 | 0.02 |
| 6.97E-05 | rs773863 | 19 | 16876428 | 6250139 | 19 | 16863640 | G/A | A | -3.98 | F2RL3 | 0.03 |
| 7.09E-05 | rs10415034 | 19 | 16806205 | 6250139 | 19 | 16863640 | G/A | A | 3.97 | F2RL3 | 0.03 |
| 7.09E-05 | rs8112613 | 19 | 16807648 | 6250139 | 19 | 16863640 | T/C | T | 3.97 | F2RL3 | 0.03 |
| 7.21E-05 | rs11879590 | 19 | 16821056 | 6250139 | 19 | 16863640 | G/A | A | 3.97 | F2RL3 | 0.03 |
| 1.80E-04 | rs734567 | 19 | 16876607 | 6250139 | 19 | 16863640 | T/C | C | -3.75 | F2RL3 | 0.07 |
| 1.93E-04 | rs2608743 | 19 | 16802410 | 6250139 | 19 | 16863640 | G/C | G | 3.73 | F2RL3 | 0.07 |
| 2.00E-04 | rs11671121 | 19 | 16828738 | 6250139 | 19 | 16863640 | T/C | T | 3.72 | F2RL3 | 0.07 |

### SUPPLEMENTARY TEXT

#### Materials and Methods

<u>(i) *F2RL3* epidemiology</u>

*Sample*

We studied individuals from the third wave of data collection in the Copenhagen City Heart Study, which took place between 1991 and 1994[3]. All individuals with DNA (N=9,252) who had experienced an acute myocardial infarction (AMI) (N=1,125) were selected along with 3,151 age, sex and smoking status 1:3 matched controls. Of these, 3,302 individuals were successfully assayed for DNA methylation at at least one of the four *F2RL3* CpG sites (Fig. 1B) and 3,205 had DNA methylation measured at all four positions. Characteristics of the study population are shown in Table S1.

*Acute myocardial infarction*

Participants in the Copenhagen City Heart Study were followed up prospectively from the date they were seen at the 1991-1994 data collection until 10th November 2014. AMI diagnoses were ascertained from the national Danish Patient Register, from the national Danish Register of Causes of Death, and from medical records of general practitioners and hospitals. AMI was defined according to WHO International Classification of Diseases 8th edition (ICD-8) code 410 until 1993, or 10th edition (ICD-10) codes I21-I22 from 1994 onwards[4].

*Mortality*

Information on vital status, date of death or emigration (n=2) was obtained from the national Danish Civil Registration System from date of examination until November 2014.

*Smoking*

Individuals were classified as never, former or current smokers according to their self-reported smoking status at the time of the 1991-1994 data collection. Current and former smokers were asked about age of smoking initiation, age of smoking cessation and consumption of cigarettes, cheroots, cigars and pipe tobacco. From this information, pack years of smoking were calculated; one pack year corresponds to smoking 20 cigarettes or equivalent per day for one year. Exposure to passive smoking at home was also self-reported.

*DNA collection*

DNA was isolated from frozen whole blood samples using the Qiagen Blood Kit resulting in an eluate with 5-30ng/µl. Twenty µl of this was treated with bisulfite using the EZ-96 DNA methylation Gold kit (Zymo Research). Elution volume was 60µl.

*F2RL3 DNA methylation*

PCR and pyrosequencing were performed using 96-plate format with samples from participants and EpiTect control DNA, 100% and 0% methylated (Qiagen, Manchester, UK).

Four µl of the bisulfite treated DNA was added to a 50µl PCR reaction containing 1x Hot star Taq mastermix (Qiagen, Manchester, UK), 0.2µmol /L of each oligonucleotide (one biotin-labeled) and dH2O. PCR was performed using a Senquest PCR machine (Genflow, UK). The thermal cycle was programmed for 95˚C for 15 minutes, followed by 50 cycles of 95˚C for 15 seconds, 52˚C for 30 seconds, and 72˚C for 15 seconds. There was a final extension step of 72˚C for 5 minutes before amplicons were cooled to 4˚C. PCR products were confirmed by electrophoresis at 80V for 45 min in a 1% (W/V) agarose gel, in 1 x TBE buffer with 10µl Safeview nucleic acid stain (NBS (Cambridgeshire, UK) per 100ml and examined in UV light.

Pyrosequencing assays and primers were designed using the PyroMark Assay Design Software 2.0 (Qiagen Manchester, UK). Pyrosequencing was carried out using a PyroMark 96 ID pyrosequencer (Qiagen Manchester, UK) according to the manufacturer's recommendations. Forty µl of amplicons were used for downstream single strand preparation and hybridisation of 0.2µmol/L sequencing primer using a Qiagen vacuum prep tool and workstation according to manufacturer’s instructions.

Oligo was purchased from IDT (Leuven, Belgium) and details were as follows:

| Name F2LR3 | Sequence 5'->3' |
|---|---|
| Forward oligo | GGGTTGGGTGTTTATTAGGT |
| Reverse oligo | /5BiosG/ACCAACAACAACACTAAACCATACATATA |
| Sequencing oligo | GTTTTGGTGGTGGGGTT |

CpG captures: 4. Genomic location: chr19:16,889,739-16,889,790, amplicon length: 288. CpG sites: 742, 757, 775, 786

*AHRR DNA methylation*

Full details of the methods for measurement of *AHRR* DNA methylation have been published previously[5]. DNA methylation of cg05575921 was assessed using a Taqman assay developed in house. The bisulfite treated DNA was amplified using forward and reverse PCR primers, which were designed to bind to DNA around the cg05575921 site on sequences without genetic or possible CpG DNA methylation variation. The probes detected either the unmethylated – and therefore converted T residue - or the methylated - and therefore conserved C residue.

| Name | Sequence 5'->3' |
|---|---|
| Forward primer | GGGATTGTTTATTTTTGAGAGGGTAGTTT |
| Reverse primer | CTACCAAAACCACTCCCAAAACC |
| Probe detecting unmethylated cg05575921 | VIC-AACCCAACCAAATACA |
| Probe detecting methylated cg05575921 | FAM-AACCCAACCGAATACA |

The thermal cycling profile was a conventional Taqman profile: 10 minutes at 95°C followed by 50 cycles of 15 seconds at 94°C, 60 seconds at 60°C, followed by cooling at 4°C.

After the end of the PCR reaction, the plates were read in a ViiA 7 Real-Time PCR System (Life Technologies). Samples were failed if the DNA methylation percentages from the duplicates were more than 30% from each other. Failed samples were measured again.

Therefore, valid measurements of DNA methylation were available for more than 99.8% of available DNA samples.

Each 384-well plate contained two identical internal control samples. Plates failed if the DNA methylation percentages from these were more than 30% discrepant. Imprecision was measured across plates with the use of the result of internal control samples. Mean, standard deviation, and coefficient of variation were calculated. Coefficients of variation varied from 5.0 to 6.7% for different stocks of the internal control.

*Statistical analysis*

All analyses were conducted in Stata (version 14.2)[6]. Associations between smoking status, pack years smoked, number of cigarettes smoked per day and percentage DNA methylation at *F2RL3* were assessed using linear regression. We also investigated the association between *F2RL3* DNA methylation and DNA methylation at another smoking-related DNA methylation site located on the *AHRR* gene using linear regression. Associations between *F2RL3* DNA methylation and AMI were assessed using logistic regression, adjusting for age, sex, smoking, pack years smoked and DNA methylation at *AHRR*. Individuals who were diagnosed with AMI prior to wave 3 (N=207) of data collection were excluded from the analysis. For analysis of mortality following AMI, we used Cox regression to calculate hazard ratios for the associations between DNA methylation /smoking and mortality. Individuals entered the analysis at the date of their wave 3 clinic and were censored upon death or 14th November 2014 if they were alive on this date.

We assessed the potential for collider bias induced by conditioning on AMI case status in our analysis of the association of *F2RL3* DNA methylation with mortality[7]. To do this, we investigated whether there was evidence for correlations between *F2RL3* DNA methylation and other risk factors for AMI in our case only sample, that were not present in the case control sample. We explored BMI, triglycerides and LDL cholesterol (assessed at wave 3) as measured risk factors in this analysis.
BMI was calculated as weight(kg)/(height(m)$^2$). Colorimetric and turbidimetric assays were used to measure non-fasting plasma levels of total and high-density lipoprotein (HDL) cholesterol and triglycerides[8]. LDL cholesterol was calculated using the Friedewald equation when plasma triglycerides were ≤4.0 mmol/l; if not, they were measured directly using a colorimetric assay.

Firstly, we calculated the association of each of the traits (converted into z-scores) with AMI using logistic regression (Fig. S5a). Secondly, we used linear regression to assess the association of these traits with *F2RL3* DNA methylation in the full AMI case control sample and in the sample consisting of AMI cases only (Fig. S5b).

(ii) *F2RL3* DNA methylation in two cell models

*Cell culture – Human coronary artery endothelial cells*

Human coronary artery endothelial cells (HCAEC) were purchased from PromoCell and cultured in Endothelial media MV2 (PromoCell). Cells were plated 24 hours before treatment with cigarette smoke extract (CSE)[9], 3 doses 16 hours apart, analysing cell response 16 hours after final treatment (total 48 hours)[10]. This was performed in triplicate using HCAEC from three different donors. The CSE is administered to cells as a bolus every 16 hours, because of the experimental complexity of administering smaller doses at shorter time intervals. This very approximately equates to an average person smoking 25 cigarettes[9].

*Cell culture – Acute megakaryocytic leukemia cells*

Acute megakaryocytic leukemia (CMK) cells were plated 2 hours before treatment with CSE, 4 doses 24 hours apart, analysing cell response 24 hours after final treatment (total 96 hours). CSE dosing very approximately equates to an average person smoking 25 cigarettes every 24 hours.

*F2RL3 DNA methylation*

DNA was isolated using the Norgen DNA/RNA extraction kit, resulting in an eluate with 5-50ng/µL. DNA was concentrated by ethanol precipitation and resuspension into a smaller volume where necessary. One µg of DNA was treated with bisulfite using EZ DNA methylation kit (Zymo Research, Irvine, CA) and eluted in 15µl. PCR and pyrosequencing were then performed using the same methods as described for '*F2RL3* DNA methylation' above using 96-plate format with samples from participants and EpiTect control DNA, 100% and 0% methylated (Qiagen, Manchester, UK) in duplicate. Samples were excluded from results if the DNA methylation percentages from duplicates failed or differences were more than 5%. It was possible for the methylation values of untreated samples to be lower than those treated, however runs where untreated samples were lower than the 0% negative controls were also failed and excluded.

For the CMK experiment, we performed 3 independent experiments comprised of pairs of untreated and CSE treated cells (1: n=2, 2: n=3, 3: n=4, total n=9). Data quality was assessed using the criteria described above. We included both sets of samples from experiment 1, were forced to exclude all 3 sets of samples from experiment 2 due to the controls exhibiting lower methylation recordings than the 0% negative controls and 2 of the 4 sets of samples from experiment 3 as the duplicate runs failed. This generated n=4 sets of samples which had data of sufficient quality to be included in our main analyses. These conditions were designed to only allow data of sufficient quality into analyses, however all data are available at data.bris (https://data.bris.ac.uk/data/group/health-sciences).

*F2RL3 mRNA*

Changes in HCAEC *F2RL3* mRNA expression were analysed by Quantitative PCR (qPCR). qPCR was performed using the Roche SYBR green PCR mastermix on cDNA prepared using the QuantiTect Reverse Transcription Kit (Qiagen). qPCR for *F2RL3* was performed using primers GACTGCTCCTGTGGCCCC and GTGCTGTCATCACCACCTCC with an anneal temperature of 62°C.

Changes in CMK *F2RL3* mRNA expression were analysed by qPCR performed using the ThermoFisher Scientific TaqMan PCR mastermix on cDNA prepared using Superscript IV VILO master mix (ThermoFisher Scientific). qPCR for *F2RL3* was performed using TaqMan Gene Expression Assay Hs01006385_g1 (FAM). Eukaryotic 18S rRNA endogenous control (Hs99999901_s1 (VIC)) was used in all samples. Ribosomal protein lateral stalk subunit P0 (RPLP0; Hs99999902_m1 (FAM)) was used as an additional control and was unchanged.

*Global DNA demethylation with 5-Azacytidine (HCAEC only)*

HCAEC were seeded into a 6-well plate at 1.4 x $10^5$ cells per well. 2.5 hours after seeding 5-azacytidine (5-AZA at a final concentration of 2.5μM, Sigma, UK) or DMSO control (1/4,000 final) was added to the cells. Media was replaced on day 2, 3 and 4, with fresh 5-AZA or DMSO control. Cells were analysed for gene expression (as described above) on day 5. This was performed four times using HCAEC from four different donors.

*Statistical analysis*

Analyses were conducted in Stata (version 14.2)[6]. Average *F2RL3* DNA methylation values (across the 3 cases and 3 controls) at each of the four sites were compared using two sample *t*-tests assuming unequal variances. Ratios of *F2RL3* mRNA levels in cases (n=6) compared to controls (n=4) were natural log transformed and an average taken. The average log ratio for cases was compared to the baseline value in controls, 1 (0 when log transformed), using a one sample t-test. The average log ratio was then exponentiated along with its 95% CIs. Ratios of DMSO control (n=4) and HCAEC cells treated with 5-AZA (n=4) were log transformed and averaged. These were compared to the baseline value in untreated controls, 1 (0 when log transformed), using one sample t-tests. The average log ratios were exponentiated along with their 95% CIs.

<u>(iii) Functional regulation of *F2RL3*</u>

First, we used a pGL3 reporter vector in which luciferase expression is driven by a heterologous SV40 promoter to test for the presence of an enhancer within a 184 bp fragment of *F2RL3* exon 2 containing CpG_1 to CpG_4. The pGL3 was a kind gift from Dr G Sala-Newby, University of Bristol. The 184 bp *F2RL3* exon 2 sequence (see below) was generated by PCR using the primers GATCTTAAGCTTACAGTGACACCCTGGAGC and AAGCTTAAGATCAGGTTCATCAGCAGCATG, which included *Hind*III restriction sites to enable cloning into the multiple cloning sites of the vectors.

Second, the potential mechanisms of effects on *F2RL3* expression were explored by transfecting HEK-293 cells with reporter constructs containing different fragments of *F2RL3* to drive expression of luciferase. The pCpGL vector used was a generous gift from Prof M Rehli, University of Regesnberg[11]. The *F2RL3* putative promoter (*F2RL3pro*) sequence corresponding to the 2 kB sequence immediately upstream of the *F2RL3* transcription initiation site (see sequence below) flanked by *Bam*H1 and *Hind*III restriction sites was synthesised (Eurofins). Inspection of the *F2RL3* exon 2 region with MatInspector[12] to identify candidate DNA-binding protein interaction sites[13] revealed a recognition site for CCAAT/enhancer binding protein (CEBP) at 2 bp 3’ of the CpG_1 DNA methylation site (Fig. 1A)[a]. The exon 2 ∆CCAAT fragment was generated from the intact exon 2 fragment by site directed deletion of the CCAAT sequence using the primers

[a] Of note, CEBP is a transcriptional regulator in multiple tissues including haemopoietic cells and vascular endothelium[45] in which *F2RL3* is also expressed[46]. CEBP binding to an identical CCAAT recognition site at a different locus (*MLH1*), is known to be reduced by DNA methylation of a CpG residue in an identical relative position to the CCCAT recognition sequence to that observed with CpG_1[47].

GGGCTGGCGCTGTGGGTG and CCGGCAGCCCCACCACCA (Phusion site directed mutagenesis kit, New England BioLabs).

*F2RL3 2 kB promoter sequence*

CTGGCTTTTTTTTTTTTTTTTTTTTTTTTTTGAGTCGGAGTCTCGCTCCG
TTGCCCAGGCTGGAGTGCAGTGGTGTGATCTCAGCTCACTGTAACCCCTG
CGTCCTGGGTTCAAGAGATTTTCCTGCCTCAGCCTCCTGAGGAGCTAGGA
CTACAGACGTGCGCCACCATGTCTGGCTAATTTTTGTATTTTTAGTAGAG
ACGGGTTTTCACCATGTTGGCCAGGACGGTCTCGATCTCTTGACCTCGTG
ATCCACCCACCTGGCCTCCCAAAGTGCTGGGATTACAGGCGTGAGCCATT
GCGCCCGGTCGTGCATCACTATTTTGATGCATCATCTTCAAACCACCCTG
CCCCAGCATCACTGGACTGCCGGTGTGCCCAGCCTCCCCTGCACAGCTTC
CCACTCTGCTCACAGAGAAGACGGTGGAGGGGGGAGCCCAGGCGTGGGTT
TTCCGTGTGCTGAGGGTGTCCTCAGCCACCTTTTTCCCCAAGTCATTGAC
GTGAAGCTGATTTTCTCATTGGTGGCCATGGAAACTGCCCAGAAATGGGA
GCAGGGCCTCTGAGACCCTGCCCCAGCTAGACTCTCTCTCCAGGCTTCAG
TGTCTCCTCCTGGAGCTGGGTGTTGACTCTGTCCAAGACTCTGCTCAGTC
ACTTCCTGGCTGATCCCCGAGTGCTCAAGTGTGTGATGAGATTAGGGCTC
CCCCAGGCAGGAAACGGCCACATTGGACCCCTCCCTCCCCCATCCCTAGT
TCATTTTATTTTCCCCAATCCTCATGGCTCCCAACTAGGCCCTCCTGCTC
CCAGACCCCCCCAAGCCCTGTCCCCCATCCCCTGTAGCCCAGCGCTGAGC
ACACTGGGGCCAGAACCACCCATACCAGACTCCCCCAGGGCACGACGGGT
TGAGGCCTCTCTGAGGCAGGGAGGGACAACACTGAGGGGCCTTGTGCAGA
AGAGGGAAGGGGTACCCCCCCGAGCTTCAGTTTCTTCATCTGCCCGATGA
GCCCCCATCCAATGGTACCCAGGAGTGGGGATGTGCTGGGGCTAGAGGAG
GGAGGCCCCCTGGAGGCTGGCTTGGGGACCCTTCAACCTGGCCACCTCAC
TCAGGGTCAGAGGTCAGAGCAGGCGAGCCACACACCCCGAAGTCCCCCCG
GTCCTCGGCAGGTGCTGTCCCCAGCATCACTGTCTCCCCTACAGATGCCC
GTGTGTCCCCTGGGCAAGAGTGAAGTGAGGCACTCATACAGGGGTGCCTT
GTGAGTATGGGGTCAGCCGAGGATGCTGAGGGGCTCTCGAGGTTCAGCCA
GAGTCCCTGACGCTGGGCTCGTGCTGCTCTGGGGGTCTGAAGGGACCCTC
GCTATTACCCTCTGGGAGGCGCCCATCCTTGGTTTTTTTTTTTTTTTTTG
CCGAGTCTCGCTCTGTTGCCCAGGCTGGAGTGCAGTGGCAAGATCTCGGC
TCACTGCAACCTCCACCTCCTGGGTTCAAGCAATTCTCCCACCTCAGCCT
CCTGAGTAGCTGGGACCACAGGCATCTGCCACCAAGCCCAGCTAATTTTT
GTATTTTTATGAGAGATGGGGTTTCTCCATGTTGGTCAGGCCGGTCTCAA
ACTCCCAACCTCAGGTGATTTGCCTGCCTTGGCCTCCCAAAATGTTGGGA
TTACAGCCGTGAGCCACCGTGCCTGGCCTAGCAGGGCTCTTCTTATCCTG
CTCACCAGAGATAGGAAGTCGGGGGGTGACTTCAGGAAAGGCCCAGAGCC
CGTGATGGGCGTGGGGATTGGGGTGAGGACAGGGCTGTCGTGAGAACAGT

```
GGCTGCAGATGTGAGCGGCTGGCAGGAAGTGGCCACTTGAACCGCAGATG
CTTCCTGGGCTGGTCAGGGACCGGGGGTGCTGGCTGCAGCTGGGACCCCC
CCCCCTGCATCTTGCTGGCCTGGCACCTGGGTCCCTGGGAGGCGCCACAC
TGGATATAGCCACGTGGGGCAGCCCGGTCTCCATAACCCACACTCCAGTC
```

Expression cassette analysis was performed in HEK293 cells. $5x10^4$ cells were seeded into each well of a 24-well plate and transfected with 425ng test plasmid expressing firefly luciferase with 75ng of Renilla-luciferase control using 2µl Viafect (Promega, UK). Analysis of gene expression was performed after 48 hours using Dual-Luciferase Reporter Assay System (Promega, UK) and expressed as a ratio of firefly luciferase/Renilla luciferase.

*Statistical analysis*

Statistical analyses were performed in Stata version 14.2[6]. Ratios of luciferase expression in HEK-293 cells (n=3) transfected with a pGL3 vector containing *F2RL3* exon 2 with CpG_1 to CpG_4 compared to pGL3 vector alone were log transformed and averaged. The average value was then exponentiated along with its 95% CI. Ratios of luciferase activity in HEK-293 cells transfected with different exon2 fragments relative to the promotor only construct (n=6) were log transformed and averaged. These were tested against the baseline value, 1 (0 when log transformed), using one sample t-tests. Luciferase activity in the vector containing the *F2RL3* exon 2 fragment was tested against the same fragment following deletion of the CCAAT binding site using a two-sample t-test. Average log ratios were exponentiated along with their 95% CIs.

**Chromatin immunoprecipitation (ChIP) in human coronary artery endothelial cells (HCAEC)**

*Sample preparation*

Following 5-Aza treatment of HCAEC as described above (under the subheading '*F2RL3* DNA methylation in a cell model'), cells were washed with cold PBS and fixed with 1% formaldehyde at room temperature for 10 min, before being quenched using 1/10 volume of 2.5 M glycine for 5 mins at room temp. Cells were washed twice with 20 ml of ice-cold phosphate-buffered saline. Cell Lysis was performed in 0.75 ml of SDS ChIP lysis buffer [1% SDS, 10 mM EDTA, 50 mM Tris-HCl, pH 8.1, 1 mM PMSF, 1 ug/ml leupeptin, 1 ug/ml aprotinin]. Samples were aliquoted in 300µl portions and stored at -80°C. DNA was sheared using 2.5 cycles (10x 30sec on/30sec off) in a Diagenode-Bioruptor sonicator (high power setting at 4°C) and centrifuged at 15k rpm at 4°C for 10 minutes. Protein concentration was measured using the Pierce Micro BCA Protein Assay.

*ChIP*

150µg of protein was used for ChIP, all samples were made up to the same volume using ChIP lysis buffer to ensure the SDS concentration were equivalent. Sample volume was increased with ChIP dilution buffer [1% Triton X-100, 2 mM EDTA, 150 mM NaCl, 20 mM, Tris-HCl, pH 8] with 1/100 Protease inhibitor (Sigma), to ensure that cell lysates were diluted at least 10 times. 2.5µg Rabbit anti-CEBP Beta antibody [E299] (ab32358, Abcam) or IgG control (Vector) were added and incubated overnight at 4°C on a rotary mixer.

*Magnetic bead preparation*

At same time as ChIP incubation, 25µl of Dynabeads® Protein A (30 mg/mL, #10002D, Life Technologies) were blocked at 4°C overnight with 75µl of blocking buffer [50 µg/mL Deoxyribonucleic acid from salmon sperm (#31149, Sigma), 0.5% BSA (w/v), in ChIP Dilution buffer]. Beads were then washed once with sterile PBS and twice with ChIP dilution buffer, using the magnet to pull the beads down in between each wash.

*Binding of immunoprecipitate (IP) onto beads*

The IP and non-immune control were added to the blocked and washed beads and incubated at 4°C for 3 hours on rota. The magnetic beads were then sequentially washed at 4°C for 5 minutes with 500µl of wash buffer 1 [0.1% SDS, 1% Triton X-100, 2 mM EDTA, 20 mM Tris-HCl, pH 8.1, 150 mM NaCl]; 500µl of wash buffer 2 [0.1% SDS, 1% Triton X-100, 2 mM EDTA, 20 mM Tris-HCl, pH 8.1, 500 mM NaCl]; 500µl of wash buffer 3 [0.25 M LiCl, 1% Nonidet P-40, 1% deoxycholate, 1 mM EDTA, 10 mM Tris-HCl, pH 8.1]; and twice with 500µl of TE [10mM Tris, 1mM EDTA, pH 8.0]. The TE was removed and the beads were incubated with two sequential 15 minute elution steps (2 x 75µl of elution buffer [1% SDS, 100mM NaHCO3]) at 65°C, vortexing briefly every 5 minutes. A 'total input' sample was also prepared containing 1/16$^{th}$ of original input in 150µl elution buffer.

*Reverse cross link and sample extraction*

6µl of 5M NaCl (200mM final) was added to each tube (IP, non-immune and total input), mixed and incubated overnight at 65°C. Samples were vortexed vigorously (2 minutes for each tube) and 150µl of 10mM Tris pH 8 added to each tube. Samples were phenolchloroform extracted, DNA precipitated and dissolved in 10mM Tris pH8. 1 μl of sample was used in each PCR reaction.

*Quantitative PCR:*

Quantitative PCR was performed using Roche SYBRgreen 2x mastermix with an anneal temperature of 67°C using primers SW855F (5' –GACAGTGACACCCTGGAGCTCCC) and SW858R (5'- CCAGCACCCACAGCGCCAGC). The relative occupancy of the *F2RL3* exon 2 CEBP recognition site was estimated as the ratio of the intensity of the specific IP band to that of the mock IP band in gel electrophoresis.

*Statistical analysis:*

Ratios of the % occupancy of the *F2RL3* exon 2 CEBP recognition site with CEBP-β in 5-AZA treated cells compared to controls were calculated for each batch of cells (n=5). Ratios were natural log transformed and the mean taken. The mean was compared to 0 (log of the value in controls) using a one sample t-test. The log ratio and its 95% CI were exponentiated.

<u>(iv) Differential platelet function in a human experiment</u>

*Study Design*

A recall by phenotype design was implemented in the Avon Longitudinal Study of Parents and Children (ALSPAC). ALSPAC is a trans-generational prospective birth cohort that began with the recruitment of 14,541 pregnant women resident in Avon, UK with expected dates of delivery 1st April 1991 to 31st December 1992. Since then, the health and development of mothers and their children has been followed across the life-course[14]. Further details of the cohort are available below in "Supplementary Text: The Avon Longitudinal Study of Parents and Children (ALSPAC)" and the study website contains details of all the data that is available through a fully searchable data dictionary (http://www.bristol.ac.uk/alspac/researchers/access/). Participants were recruited based on DNA methylation at *F2RL3* as measured by Illumina Infinium HumanDNA methylation 450 BeadChip (450 K) array[15].

*Ethical considerations and informed consent*

Details of the ethics approvals relevant to the ALSPAC cohort in general can be found on the study website (http://www.bristol.ac.uk/alspac/researchers/research-ethics/). Ethical approval for this study was obtained from the ALSPAC Ethics and Law Committee and the

NHS South West - Frenchay Research Ethics Committee (REC reference 14/SW/1099). All participants received a participant information sheet prior to being recruited to the study and were given the opportunity to ask questions both at the telephone screening and at their appointment. All participants were asked to complete a written consent form. Participants were appropriately reimbursed for their time and effort (as judged against other contemporary study initiatives) and all travel expenses reimbursed.

*Participant recruitment*

Between May 2015 and January 2016, we recalled young people from the ALSPAC cohort who had had DNA methylation assessed as part of the ARIES project (total N=1,007)[15]. Genome-wide DNA methylation profiling was conducted in these individuals from tissue taken at birth (cord blood), childhood (average age 7) and adolescence (age 15 or 17 years) (peripheral blood) using the Illumina Infinium HumanDNA methylation 450K BeadChip assay[15] (see “Supplementary Text: The ALSPAC Cohort” for further details). In this study, data from the childhood and adolescence datasets (release v2[16]) were used post-normalisation by a pipeline described by Touleimat and Tost[17]. In addition, we adjusted DNA methylation values for cell composition (using estimated fractions of CD8 T cells, CD4 T cells, NK cells, B cells and monocytes), sex, clinic attended in the case of the adolescent measures (whether age 15 or age 17) and the technical artefacts of chip row and plate; a mixed model was fitted in which all factors were fitted as fixed effects except plate which was treated as a random effect.

In order to be eligible for invitation to the study, individuals had to have DNA methylation data available at both time points (childhood and adolescence) and have genome-wide genotype data (details of the SNP genotyping, imputation, processing and quality control procedures carried out in ALSPAC are in “Supplementary Text: The ALSPAC Cohort”). As we were interested in *F2RL3* DNA methylation independent of smoking, we excluded individuals who reported; (i) smoking daily or weekly when they were age 13.5; or (ii) having smoked more than 100 cigarettes in their lifetime at age 15; or (iii) being daily smokers at age 15. The 731 individuals that remained eligible for invite were ranked firstly according to DNA methylation at age seven and then according to DNA methylation at age 17; the average of these two rankings was then used to prioritise individuals for invitation to the study, with those with the highest average ranking and those with the lowest average ranking selected for invite. Initially, the top 50 and bottom 50 were invited to participate; this was extended to the top and bottom 100 during the course of the study to meet target recruitment numbers. Some restrictions were imposed by the cohort (related to participant's commitment to other studies running concurrently with ours) such that the final number of invitations issued was n=147. The two groups, selected on the basis of high and low DNA methylation, did not differ substantively based on other relevant characteristics measured in the cohort (Table S5). Researchers and fieldworkers remained blind to participant status (high versus low DNA methylation group) throughout the recruitment, data collection and laboratory analysis phases of the study.

Individuals responding positively to the invitation were screened in a telephone interview and invited to attend clinic if they: 1) had never been a regular smoker; 2) did not have any haemostatic or cardiovascular disorders; 3) were not anaemic; 4) were not on insulin treatment; 5) had not had a major illness in the past; 6) did not regularly take anticoagulants, antiplatelet drugs or NSAIDS; 7) were not dependent on any substance apart from caffeine. Individuals eligible to take part were asked not to take ibuprofen or aspirin for seven days prior to their scheduled clinic visit; clinic visits were rescheduled in cases where such drugs had been taken in the seven days preceding their clinic visit. At the clinic, individuals were screened again for the exclusion criteria.

A total of 49 individuals were recruited to the study and had blood samples taken. Eight participants were excluded due to not meeting exclusion criteria or technical issues during laboratory analysis. Individuals recruited to the study and whose data was retained for analysis (n=41) did not appear to differ from those not recruited based on other relevant characteristics measured in the eligible cohort (Table S6).

*Sample Collection*

Participants provided a non-fasting blood sample (4.5mL) collected according to standard procedures. This was stored at room temperature in a vacutainer containing 1:9 v/v 4% trisodium citrate (BD Biosciences, Oxford, UK) and transported to the laboratory for analysis within two hours of being taken (range 21 – 79 minutes).

*Sample analysis*

*Preparation of platelet-rich plasma (PRP):* Haematologic parameters were tested using a Horiba Pentra ES60 Plus haematology analyser (Horiba UK Ltd, Northampton, UK), prior to centrifugation at 180 x g for 17 minutes. The PRP layer was removed and a platelet count was performed on a Z1 Coulter Particle Counter (Beckman Coulter, High Wycombe, UK). PRP was diluted in a modified HEPES-Tyrode's buffer (145 mM NaCl, 2.9 mM KCl, 10 mM

HEPES, 1 mM $MgCl_2$, 5 mM glucose, pH 7.3) to $3x10^7$ platelets/mL. The buffy layer containing white blood cells was removed from the centrifuged whole blood and snap frozen in liquid nitrogen and stored at -80°C for subsequent DNA extraction and pyro-sequencing (see '*F2RL3 DNA methylation*' below).

*Flow Cytometry:* For assessing $\alpha_{IIb}\beta_3$ integrin activation and P-selectin exposure, diluted PRP at a platelet count of $3x10^7$ plts/mL was stimulated for 10 minutes at room temperature with either the PAR1-specific activating peptide, SFLLRN (0.25, 0.5, 1.0, 2.0, 3.0, 5.0 and 10.0 μM), or the PAR4-specific activating peptide, AYPGKF (20, 30, 50, 75, 100, 200 and 400 μM), in 50 µl reactions in the presence of 5 µl FITC conjugated mouse anti-human PAC-1 and 2.5 µl PE-conjugated mouse anti-human CD62P (BD Biosciences, Oxford, UK) per reaction. Unstimulated basal samples in the presence or absence of antibodies were used as controls. Unstimulated basal samples in the presence of 5 mM EDTA were included to give a measure of basal integrin activation. Samples were then fixed with an equal volume of 2% paraformaldehyde prior to analysis on a FACSCanto flow cytometer (BD Biosciences, Oxford, UK).

For assessing platelet surface receptor expression, diluted PRP at a platelet count of $3x10^7$ plts/mL was incubated in 50 µl reactions in the presence of PE-conjugated mouse anti-human CD41 (Thermo Fisher (Invitrogen), Paisley, UK), CD61 or IgG isotype control (both from BD Biosciences, Oxford, UK) for 10 minutes at room temperature. Samples were then fixed with an equal volume of 2% paraformaldehyde prior to analysis on a FACSCanto flow cytometer (BD Biosciences, Oxford, UK).

*F2RL3 DNA methylation*

To provide a measure of contemporary DNA methylation at the locus, previously separated (and frozen) buffy layer was thawed, red blood cells lysed and DNA extracted using a standard manual guanidine HCl/chloroform extraction method. DNA was normalised at 50ng/µl. 1µg of DNA was treated with bisulfite using EZ DNA methylation Kit (Zymo Research, Irvine, CA) and eluted in 15µl. PCR and pyrosequencing were then performed using the same methods as described for '*F2RL3* DNA methylation' above. PCR and pyrosequencing were performed using 96-plate format with samples from participants and EpiTect control DNA, 100% and 0% methylated (Qiagen, Manchester, UK) in duplicate. Samples were failed if the DNA methylation percentages from duplicates were more than 5%.

*Statistical analysis*

For assessing $\alpha_{IIb}\beta_3$ integrin activation and P-selectin exposure, dose response curves of activation responses versus PAR4 agonist concentration were obtained from non-linear regression of log[agonist concentration] versus response performed using GraphPad Prism (version 7.00 for Windows, GraphPad Software, La Jolla California USA, www.graphpad.com). A by-group comparison of dose response curves was carried out by two-way ANOVA. By-group comparisons of $\alpha_{IIb}\beta_3$ integrin activation and P-selectin exposure $EC_{50}$ and maximum response values, measured platelet surface receptors and haematological parameters were compared by two-sample Wilcoxon rank-sum (Mann-Whitney) test after the removal of outliers identified on the basis of a robust regression of $EC_{50}$ (or maximum response) on DNA methylation group (high/low).

The contemporary (pyrosequencing-based) DNA methylation values were used to explore the linear relationship between DNA methylation at each of the positions and platelet reactivity. Linear regression models of the outcome (integrin activation $EC_{50}$ or p-selectin exposure $EC_{50}$) on DNA methylation (at each position) were fitted within selection group. At CpG_1, the beta coefficient for integrin activation $EC_{50}$ was similar in the two selection groups so we also performed a combined analysis with DNA methylation group (high/low) included as a covariate in the model. Fitting measured confounders in the model showed no strong evidence of association between DNA methylation and sex, age, time from blood draw to assay, laboratory technician carrying out the assay or platelet count.

*Genetic Analysis:* Pre-existing genetic data (details of the SNP genotyping, imputation, processing and quality control procedures carried out in ALSPAC are in “Supplementary Text: The ALSPAC Cohort”) were used to determine if the genetic variant rs773902 in *F2RL3* was associated with DNA methylation. A by-group comparison of allele frequency at rs773902 (A/G) was performed using a chi-squared test both in those selected for invite (n=200, Table S5) and in those actually recruited (n=41, Table S11). A linear model of DNA methylation (%) on rs773902 genotype was fitted first using CpG_1 DNA methylation measured in this study (n=33 with data) and secondly using pre-existing DNA methylation at cg03636183 (CpG_3) measured in the entire ARIES v2 collection (n=731)[15,16]. Finally, a linear model of outcome (integrin activation $EC_{50}$ or p-selectin exposure $EC_{50}$) on rs773902 genotype was fitted to assess the association between genotype and platelet reactivity.

(v) Two-step epigenetic Mendelian randomization

A two-step epigenetic Mendelian randomization (MR) strategy[18,19] was used to interrogate the causal relationship between smoking, DNA methylation and CVD outcomes (see Fig. S1).

*Step one*

In the first step, the causal impact of smoking (a modifiable risk factor) on the epigenetic signature at *F2RL3* was evaluated. This analysis was performed using data from the Copenhagen City Heart Study (as described above). A genetic variant in the *CHRNA5-A3-B4* gene cluster, rs1051730, which is robustly associated with tobacco consumption amongst smokers (i.e., cigarettes smoked per day, plasma cotinine levels)[20,21] was used as the instrument. The rs1051730 genotype of the nicotinic acetylcholine receptor gene (*CHRNA3)* was determined with a Taqman assay using methods described previously[22]. We assessed the association between the rs1051730 SNP and cigarettes per day (in current smokers) and pack years (in ever smokers) using linear regression, assuming an additive genetic model and adjusting for age and sex (Table S12). Beta coefficients were expressed per additional copy of the minor allele of rs1051730. In Mendelian randomisation analysis, we calculated the association of rs1051730 with *F2RL3* DNA methylation using linear regression, stratified by smoking status (never, former, current and ever (former + current) smokers) (Table S13).

*Step two*

In the second step, we evaluated the causal impact of *F2RL3* DNA methylation on CVD as an outcome, and more specifically on ischaemic heart disease (IHD) (also known as coronary heart disease (CHD)), myocardial infarction (MI) and AMI as subcategories of CVD involving coronary thrombosis in a two sample MR framework[23,24]. Genetic instruments for *F2RL3* DNA methylation were *cis*-DNA methylation quantitative trait loci (*cis*-mQTLs) for cg03636183 (CpG_3) identified in a recent (unpublished) meta-GWAS of DNA methylation from 36 European cohorts (including the ARIES collection[15], a subgroup of ALSPAC with 450k DNA methylation data) performed by the GoDMC consortium (see Supplementary Text below for further details of the GoDMC analysis). In this analysis, 110 *cis*-variants were associated with cg03636183 (CpG_3) at $p<1e-5$ in at least one cohort. Using results from the meta-analysis, linkage disequilibrium (LD)-based clumping was performed in Plink[25,26] within the region (chr19:15854486-18021066) to give a set of index variants to use as instruments. Clumps were formed around index variants with $p<0.0001$ such that any sites within 5000 kb of an index variant, with an $r^2$ larger than 0.1 with it and with an association p-value of <0.01 were assigned to that index variant's clump. Clumping revealed six clumps with index variants: rs3848657, rs2227349, rs55961639, rs112791293, rs773919 and rs189158451. We used these SNPs to instrument *F2RL3* DNA methylation.

We also performed the MR analysis using rs773902 as an instrument as this variant has previously been linked to platelet function[27,28] and is associated with DNA methylation both in the ARIES collection and in the sub-group of the ALSPAC cohort recalled for our human experiment. For each SNP (n=7), the effect size (and standard error) expressed in standard deviation (SD) units for the DNA methylation decreasing allele was extracted from the GoDMC meta-analysis results (Table S14). A proxy SNP, rs812847, was used in place of rs773919 because rs773919 is palindromic with a minor allele frequency close to 0.5 introducing uncertainty during the harmonization process. rs812847 was in the same clump as rs773919 and has $r^2$=0.98 with rs773919 (based on correlations extracted from 1000 Genomes data using the 'TwoSampleMR package'[29]).

SNP-outcome associations for the seven SNPs were extracted for CHD and MI from CARDIoGRAMplusC4D consortium GWAS results[30] using the R package, TwoSampleMR which uses data available via MR-Base[29]. For each SNP, SNP to disease effect, expressed as an odds ratio (OR) per additional DNA methylation decreasing allele can be observed in Fig. S7.

SNP-outcome associations for the seven SNPs were calculated in UK Biobank for AMI, IHD and CVD (see "Supplementary Text: UK Biobank" for more details). Analyses were conducted using N=487,409 individuals with imputed genotype data (release date 20th July 2017) and phenotype data downloaded on 2nd October 2018 under research application 15825 (N=502,591). Quality control was undertaken in line with and using exclusion lists generated during quality control filtering of the UK Biobank data conducted by R.Mitchell, G.Hemani, T.Dudding and L.Paternoster, as described in the published protocol[31]. There were 487,371 with both imputed genotype data and phenotypic data available. 79,447 individuals were excluded on the basis of genetic relatedness such that the remaining subset is a maximal set of unrelated individuals, derived using an algorithm applied to all the related pairs provided by UK Biobank which preferentially removes individuals related to the greatest number of other individuals until no related pairs remain; in addition, nine apparently highly related individuals not included in the kinship inference analysis conducted by UK Biobank analysts were excluded. A further 736 individuals were excluded from the analysis based on standard exclusion criteria as defined by UK Biobank analysts (sex mismatch, putative sex chromosome aneuploidy, outliers in heterozygosity and missing rates). Thirty-five of the remaining individuals were removed in line with consent withdrawals. This left data for N=407,144 individuals to be taken forward into the analysis. In order to designate individuals as cases or controls for CVD, IHD and AMI, we used information (in the form of ICD-10 and ICD-9 codes) from six data fields: cause of death (f40001), secondary cause of death (f40002), main diagnosis (across all episodes in hospital) (f41202, f41203) and secondary diagnosis (across all episodes in hospital) (f41204, f41205). Individuals were designated as CVD cases if they had an ICD-10 code beginning with "I" and/or an ICD-9 code in the range 309-459 ('Diseases of the circulatory system') or a code beginning with "G45" ('Transient cerebral ischaemic attacks and related syndromes'), IHD cases if they had an ICD-10 code in the range I20-I25 and/or an ICD-9 code in the range 410-414 ('Ischaemic heart diseases') and AMI cases if they had an ICD-10 code I21 and/or ICD-9 code 410 ('Acute myocardial infarction') or an ICD-10 code I22 and/or ICD-9 code 412 ('Subsequent myocardial infarction') across any of the six data fields. Individuals not designated as CVD cases were designated as controls; the same control set was used in the analysis of all three outcomes. Where cause of death differed across instances, values were set to missing. One individual whose date of death differed across instances and two individuals who had cause of death data but no date of death were excluded from subsequent analyses leaving N=407,141 individuals with genotype and phenotype data for the analysis. The association of each of the seven SNPs with disease outcome (AMI, IHD, CVD) was calculated using logistic regression assuming an additive model; sex and ten principal components (as calculated by UK Biobank analysts – data field 22009) were fitted as covariates in the model (see below for further explanation regarding the model that was fitted). For each SNP, SNP to disease effect, expressed as an odds ratio (OR) per additional DNA methylation decreasing allele can be observed in Fig. S7.

The causal effect of *F2RL3* DNA methylation on disease outcome based on the multi-SNP instrument was estimated using a generalized weighted linear regression approach implemented in the R package 'MendelianRandomization'[24]. In this approach, correlation between SNPs in the set due to linkage disequilibrium is accounted for by fitting a matrix of correlations in the model (Table S15) (correlations were extracted from the UK Biobank dataset used in the main analysis, n=407,141)[32]. In addition, estimates were generated for individual SNPs using a Wald Ratio method implemented in Stata (version 14.2)[6] (Fig. S8A) and in a leave one out analysis where each SNP from the set was removed in turn (Fig. S8B).

*Additional analyses:* To explore the potential impact of survivorship bias on our results, the UK Biobank analysis was repeated such that cases were restricted to individuals who had the relevant ICD10 codes recorded in the primary or secondary cause of death variables (i.e. incident mortality). Results from these analyses can be found in Fig. S9 under 'Fatal AMI', 'Fatal IHD' and 'Fatal CVD'.

In UK Biobank, the SNPs used as instruments in the analysis described above were associated with a number of confounders, indicative of confounding caused by population stratification (Table S16A); this is expected due to differences in allele frequencies (as well as confounders) across populations. In sensitivity analyses, removing the non-White Europeans (defined as those lying outside the largest cluster following an in-house k-means cluster analysis performed using the first four principal components provided by UK Biobank in the statistical software environment R) (Table S16B) or adjusting for principal components one through ten (Table S16C) largely eliminated the association of genotype with measured confounders. Therefore, in the main analysis non-White European individuals were left in the data and ten principal components fitted in the model to account for population stratification in the sample. In sensitivity analyses, restricting the data to participants designated either as White European or as White British did not substantially alter the SNP-outcome associations (Fig. S10). In further sensitivity analyses, fitting array (either UK Biobank or UK BiLEVE) as a covariate did not substantially alter the SNP-outcome associations (Fig. S10).

The primary analysis conducted in UK Biobank (as described above) was repeated in a stratified analysis, whereby the SNP-outcome association was estimated separately for 'ever smokers' and 'never smokers'. Individuals were assigned to groups on the basis of smoking status (as defined in data field 20116) such that those with status 'former' or 'current' were designated as 'ever smokers'. Results of the instrumental variable analyses conducted using the multi-SNP instrument and rs773902 only are presented in Fig. S11 and Fig. S12, respectively.

The CARDIoGRAMplusC4D consortium GWAS from which betas were extracted for the primary MR analysis included participants of non-European ancestry (13% South Asian and 6% East Asian). In sensitivity analyses to explore possible confounding due to population structure, the same analysis was run using results from a previous iteration of CARDIoGRAM that only included European individuals[33]. This dataset was considerably smaller than CARDIoGRAMplusC4D and none of the SNPs in the multi-SNP set were present. Suitable proxies ($r^2$>0.8) were identified for four of the SNPs (rs2608739 for rs112791293; rs2227356 for rs2227349; rs2303091 for rs3848657; rs8108315 for rs55961639). The only phenotype available in this CARDIoGRAM dataset was CHD. Results of the instrumental variable analyses conducted using the multi-SNP instrument and compared to the CARDIoGRAMplusC4D result are shown in Figure S13.

### Additional results

Differential platelet function in a human experiment

*Genetic analysis:* A previous study has shown that the single nucleotide polymorphism (SNP), rs773902, located at 16,889,821 bp on chromosome 19 (Fig. 1B) is associated with platelet function. Specifically, the A-allele at the locus which encodes threonine (Thr) at residue 120 in transmembrane domain 2 (as opposed to alanine (Ala)) was associated with greater PAR4-induced human platelet aggregation and a higher level of Ca+ flux[27]. Using existing genetic data in the ALSPAC cohort, we explored the potential impact of this variant both on methylation and platelet reactivity. The A-allele at rs773902 occurred more frequently in the low DNA methylation group (Table S11) and was associated with decreased *F2RL3* DNA methylation both in this sub-study (CpG_1 4.25% (95% CI: 2.44, 6.06), $p<0.001$, n=33) per A-allele) and in the entire ARIES collection (CpG_3 (cg03636183) 1.16% (95% CI: 0.52, 1.81), $p<0.001$, n=731). When fitted in a linear model (assuming an additive genetic model) genotype was also associated with increased platelet reactivity. Each additional A-allele was associated with a 16.1 μM (95% CI: 8.4, 23.9, $p<0.001$, n=41) decrease in integrin activation $EC_{50}$ and a 20.4 μM (95% CI: 4.9, 35.9, $p=0.01$, n=39) decrease in P-selectin exposure $EC_{50}$.

### Cohort information

#### The Avon Longitudinal Study of Parents and Children (ALSPAC)

*ALSPAC: Description of study numbers*

ALSPAC recruited 14,541 pregnant women resident in Avon, UK with expected dates of delivery 1st April 1991 to 31st December 1992[14,34]. 14,541 is the *initial* number of pregnancies for which the mother enrolled in the ALSPAC study and had either returned at least one questionnaire or attended a "Children in Focus" clinic by 19/07/99. Of these *initial* pregnancies, there was a total of 14,676 fetuses, resulting in 14,062 live births and 13,988 children who were alive at 1 year of age. Detailed information on these children, their mothers and mothers' partners has been collected ever since via attendance at clinics and postal/online questionnaires.

When the oldest children were approximately 7 years of age, an attempt was made to bolster the initial sample with eligible cases who had failed to join the study originally. As a result, when considering variables collected from the age of seven onwards (and potentially abstracted from obstetric notes) there are data available for more than the 14,541 pregnancies mentioned above. The total sample size for analyses using any data collected after the age of seven is therefore 15,247 pregnancies, resulting in 15,458 fetuses. Of this **total sample** of 15,458 fetuses, 14,775 were **live births** and 14,701 were **alive at 1 year of age**.

*ALSPAC: Genotyping description*

ALSPAC children were genotyped using the Illumina HumanHap550 quad chip genotyping platforms by 23andme subcontracting the Wellcome Trust Sanger Institute, Cambridge, UK and the Laboratory Corporation of America, Burlington, NC, US. The resulting raw genome-wide data were subjected to standard quality control methods. Individuals were excluded on the basis of sex mismatches; minimal or excessive heterozygosity; disproportionate levels of individual missingness (>3%) and insufficient sample replication (IBD < 0.8). Population stratification was assessed by multidimensional scaling analysis and compared with Hapmap II (release 22) European descent (CEU), Han Chinese, Japanese and Yoruba reference populations; all individuals with non-European ancestry were removed. SNPs with a minor allele frequency of < 1%, a call rate of < 95% or evidence for violations of Hardy-Weinberg equilibrium (P < 5E-7) were removed. Cryptic relatedness was measured as proportion of identity by descent (IBD > 0.1). Related subjects that passed all other quality control thresholds were retained during subsequent phasing and imputation. 9,115 subjects and 500,527 SNPs passed these quality control filters.

ALSPAC mothers were genotyped using the Illumina human660W-quad array at Centre National de Génotypage (CNG) and genotypes were called with Illumina GenomeStudio. PLINK (v1.07)[35] was used to carry out quality control measures on an initial set of 10,015 subjects and 557,124 directly genotyped SNPs. SNPs were removed if they displayed more than 5% missingness or a Hardy-Weinberg equilibrium P-value of < 1E-6. Additionally, SNPs with a minor allele frequency of less than 1% were removed. Samples were excluded if they displayed more than 5% missingness, had indeterminate X chromosome heterozygosity or extreme autosomal heterozygosity. Samples showing evidence of population stratification were identified by multidimensional scaling of genome-wide identity by state pairwise distances using the four HapMap populations as a reference, and then excluded. Cryptic relatedness was assessed using an IBD estimate of more than 0.125 which is expected to correspond to roughly 12.5% alleles shared IBD or a relatedness at the first cousin level. Related subjects that passed all other quality control thresholds were retained during subsequent phasing and imputation. 9,048 subjects and 526,688 SNPs passed these quality control filters.

*ALSPAC: Imputation description*

477,482 SNP genotypes in common between the sample of mothers and sample of children were combined. SNPs with genotype missingness above 1% due to poor quality were removed (11,396 SNPs removed). 321 subjects were removed due to potential ID mismatches. This resulted in a dataset of 17,842 subjects containing 6,305 duos and 465,740 SNPs (112 were removed during liftover and 234 were out of HWE after combination). Haplotypes were estimated using ShapeIT (v2.r644) which utilises relatedness during phasing. A phased version of the 1000 genomes reference panel (Phase 1, Version 3) was obtained from the Impute2 reference data repository (phased using ShapeIt v2.r644, haplotype release date Dec 2013). Imputation of the target data was performed using IMPUTE V2.2.2[36,37] against the reference panel (all polymorphic SNPs excluding singletons), using all 2,186 reference haplotypes (including non-Europeans). This gave 17,842 mothers and children eligible for study with available genotype data. Subsequent consent withdrawals have left 17,825 individuals for study.

*ALSPAC: ARIES description*

Samples were drawn from the Avon Longitudinal Study of Parents and Children[14,34]. Blood from 1018 mother–child pairs (children at three time points and their mothers at two time points) were selected for analysis as part of the Accessible Resource for Integrative Epigenomic Studies (ARIES, http://www.ariesepigenomics.org.uk/) (Relton 2015). Following DNA extraction, samples were bisulphite converted using the Zymo EZ DNA Methylation™ kit (Zymo, Irvine, CA, USA). Following conversion, genome-wide methylation was measured using the Illumina Infinium HumanMethylation450 (HM450) BeadChip. The arrays were scanned using an Illumina iScan, with initial quality review using GenomeStudio. ARIES consists of 5469 DNA methylation profiles obtained from 1022 mother-child pairs measured at five time points (three time points for children: birth, childhood and adolescence; and two for mothers: during pregnancy and at middle age). Full details of the pre-processing and normalization of ARIES v2 has been described previously[15,16].

**UK Biobank**

UK Biobank is a population-based health research resource consisting of approximately 500,000 people, aged between 38 years and 73 years, who were recruited between the years 2006 and 2010 from across the UK[38]. Particularly focused on identifying determinants of human diseases in middle-aged and older individuals, participants provided a range of information (such as demographics, health status, lifestyle measures, cognitive testing, personality self-report, and physical and mental health measures) via questionnaires and interviews; anthropometric measures, BP readings and samples of blood, urine and saliva were also taken (data available at www.ukbiobank.ac.uk). A full description of the study design, participants and quality control (QC) methods have been described in detail previously[39]. UK Biobank received ethical approval from the Research Ethics Committee (REC reference for UK Biobank is 11/NW/0382).

*Genotyping and imputation*

The full data release contains the cohort of successfully genotyped samples (n=488,377). 49,979 individuals were genotyped using the UK BiLEVE array and 438,398 using the UK Biobank axiom array. Pre-imputation QC, phasing and imputation are described elsewhere[40]. In brief, prior to phasing, multiallelic SNPs or those with MAF ≤1% were removed. Phasing of genotype data was performed using a modified version of the SHAPEIT2 algorithm[41]. Genotype imputation to a reference set combining the UK10K haplotype and HRC reference panels[42] was performed using IMPUTE2 algorithms[37]. The analyses presented here were restricted to autosomal variants within the HRC site list.

### Consortia information

#### The Genetics of DNA Methylation Consortium (GoDMC) Meta-GWAS

GoDMC was established with the view of bringing together researchers with an interest in studying the genetic basis of DNA methylation variation, to consolidate as many resources and expertise as possible and thereby expedite this field of research. One of the first aims of the consortium was to carry out a meta-GWAS of DNA methylation, as measured on Illumina 450k or EPIC Beadchips. Results from a meta-GWAS involving 36 cohorts (N = 27,750) were used here. Results from the GoDMC consortium analyses are available on request from http://www.godmc.org.uk/projects.html. Here we provide a brief description of the analysis.

*Genotype data:* Genotype data of all autosomes and chromosome X (if available) was imputed to 1000G and above using hg19/build37. Genotype data was filtered on an info score of 0.8 and a minor allele frequency (MAF) of 0.01. Genotype data was converted to bestguess data without a probability cut-off.

*DNA methylation data:* DNA methylation was measured in whole blood or cord blood using Illumina 450k or EPIC Beadchips in at least 100 European individuals. Normalized beta values were used, preferable normalized with the R package meffil [35]. Most analysts used meffil to quality control and normalize the DNA methylation data using functional normalization. Protocols can be found here: https://github.com/perishky/meffil/wiki.

A github pipeline was implemented to run the analyses locally. For the genotype data, several standard sample QC steps were performed including a sex check, removal of samples with >5% missingness, and the identification and exclusion of ethnic outliers. In datasets of ostensibly unrelated individuals, those that were found to be related (identity by state > 0.125) were excluded.

The pipeline then residualised the normalized methylation betas by replacing outliers that were 10 standard deviations from the mean (3 iterations) with the probe mean, rank transforming the normalized beta values and regressing out age, sex, predicted cell counts, predicted smoking, genetic principal components and non-genetic methylation principal components. In family-based cohorts, genetic relatedness matrices were constructed and relatedness adjusted for using the GRAMMAR approach[43]. Genomic lambdas were checked by performing a GWAS of cg07959070. These residualised methylation measurements were used in all analyses.

*Association analysis:* First, every study performed a full analysis of all candidate mQTL associations, returning only associations at a threshold of p<1e-5. All candidate mQTL associations at p<1e-5 were combined to create a unique 'candidate list' of mQTL associations. In total, 102,965,711 candidate mQTL associations in cis (p<1e-5, SNP located within 1Mb of the methylation site) and 710,638,230 candidate mQTL associations in trans were identified in at least one dataset. To avoid computational burden, we included cis associations found in at least one dataset and trans associations in at least two datasets.The candidate list (n=120,212,413) was then sent back to all cohorts and the association estimates obtained for every mQTL association on the candidate list.

*Meta analyses:* Meta analyses were run using a modified version of METAL[44] using 962 chunks. Candidate mQTL associations were meta-analysed using fixed effects, additive random effects and multiplicative random effects models. 36 datasets from European origin were included in the meta-analyses.
